# Scute structure reveals sex of juvenile and young sturgeon in caviar aquaculture technology

**DOI:** 10.1101/286401

**Authors:** Nikolai V. Barulin

## Abstract

Sturgeons are valued as specialty black caviar, which is very expensive. Only females are used in the technology of caviar aquaculture. Universal method of sex determination has not yet been developed. Most of known methods are not sufficiently accurate, or used at a relatively late age, or difficult to use. Perfect early determination of sex is considered to be impossible. Because of the dark color of most sturgeon, important morphological differences, which fish of almost all ages have, were overlooked. We first found that the scute structure of Sterlet and Siberian sturgeon depends on the sex, such a pattern is typical for sturgeon of all ages. We found that sex determination is possible at very early stages. Dependencies found with the help of machine learning method open a possibility for creation of sex determination equipment using the artificial intelligence. Our results open a perspective for creation of the sex determination methods for other 23 sturgeon species, which can increase the efficiency of caviar aquaculture and sturgeon restoration in natural waters.

## INTRODUCTION

Sturgeon caviar is a worldwide famous and very expensive delicacy (Bronzi, 2014). Caviar price can reach up to 300 US dollars / ounce and more, because most sturgeon is in the phase of full extinction. Furthermore, the first maturation period of sturgeon can reach up to 10-20 years (Williot et. al., 2017). Caviar is healthy food, especially for children and pregnant women. Therefore, the caviar price should be lower. Early sex determination to cull out males is one of the ways to reduce caviar price.

However, sturgeons have no reliable external sexual characteristics, as previously thought (Chebanov, Galich, 2013). Sex determination allow the use of hormonal (Mosyagina, Zelennikov, 2016; Du et al., 2016; Wheeler et al., 2016), biochemical (Barulin, 2015), histological (Golpour et al., 2017) methods and biopsy (Falahatkar et al., 2014), laparoscopy (Falahatkar et al., 2014), endoscopy (Munhofen et al., 2014). Above methods have difficulties with mass application in aquaculture and can have a high error (the first two methods) or can traumatize fish (the last four methods). Furthermore, a sturgeon sex gene has not been found, unlike most fish species (Khodaparast et al., 2014), however, a study in this area are still under way (Vizziano-Cantonnet et al., 2016; Havelka et al., 2017). Currently, the ultrasound diagnostics is the most successful method of early sturgeon sex determination (Memiş et al., 2016) and can determine sex starting from 2 years for Sterlet sturgeon (*Acipenser ruthenus*), 2-3 years for Siberian (*A. baerii*) and Russian (*A. gueldenstaedtii*) sturgeons, 3-4 years for White sturgeon (*A. transmontanus*) and in 4-5 years in Beluga (*Huso huso*) (Chebanov, Galich, 2013). However, the ultrasonic diagnostics requires the expensive equipment and high qualification of fish farmers.

The aim of this work is to perform morphometrical study of dorsal scutes in sturgeon, to determine the correlation of their structure with sex, and to develop a system for intravital sex identification.

## MATERIALS AND METHODS

### 1. Test species

Sterlet sturgeon (*Acipenser ruthenus*) descendants from the Volga-river population and Siberian sturgeon (*A. baerii*) descendants from the Lena-river population at the age of 3 months (juvenile), 1 year (young) and 2 years (adult) kept in aquaculture, were studied. Average length of Sterlet sturgeon was: adults – 61.2±2.3 cm, young 24.8±1.5 cm, juveniles 70.3±3.6 mm. Average length of Siberian sturgeon was: adults – 73.2±2.7 cm, young 34.8±1.9 cm, juveniles 11.5±4.2 cm.

### 2. Aquaculture management

The research was carried out from 2011 to 2017 at the laboratory and Aquaculture farm of the Department of Ichthyology and Pisciculture, Belarusian State Agricultural Academy (Gorki, Mogilev region, Belarus), and the Aquaculture farms: ‘Vasilek’ (Minsk region, Belarus), “Selets” (Brest region, Belarus) and “Remona” (Mogilev region, Belarus).

All fish farms used the Recirculation Aquaculture Technology with plastic fish tanks, mechanical and biological filtrations, ozonation, aeration or oxygenation. The production systems met the basic technological requirements adopted in the modern practice of sturgeon aquaculture (Chebanov, Galich, 2013). High-protein feed from BioMar, Coppens, and Aller aqua was used.

### 3. Data generation

#### 3.1. Data collection

200 three –months-old juveniles of sterlets and Siberian sturgeon were marked with microdecimal CWT tags. These fish were re-marked with PIT tags at the age of 1 year. Then the sex and maturity stages of sturgeon gonads were checked at the age of 2 years using ultrasound diagnostics, biopsy, histological and visual methods (Chebanov, Galich, 2013) (Fig.1).

**Fig. 1.**
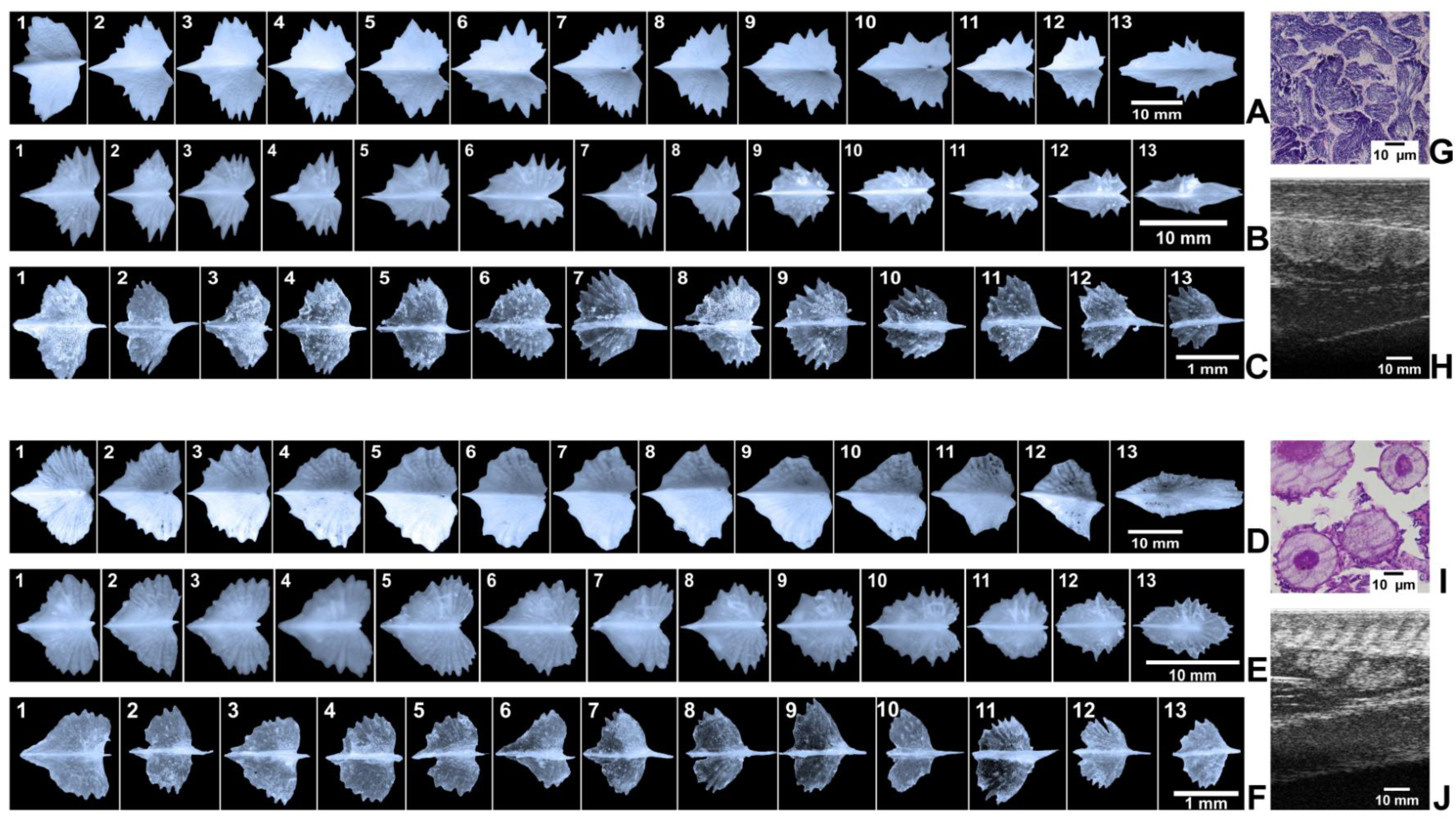
Visual morphological sex differences observed on dorsal scutes of *Acipenser ruthenus*. Showing cleared series of adults (fish length 61.2±1.3 cm, 2 years-old (A, D)), young (fish length 24.8±1.5 cm, 1 year-old (B, E)) and juveniles (fish length 70.3±3.6 mm, 3 months-old(C, F)) dorsal scutes from head (1) to dorsal fin (10); origin: Belarus, RAS of Belarusian State Agricultural Academy. Backgrounds were cleaned and contrast was enhanced using Adobe Photoshop. Used biopsy, histology and ultrasound for sex identification of adults sterlet. Showing histology (G, I) and ultrasound (H, J) images of testes (G, H) and ovaries (I, J)

We collected scute images of each marked sturgeon at the age of 3 months, 1 year and 2 years for further analysis and research. We used digital SLR camera in macro mode with enhanced flash.

#### 3.2. Sex determination

The Trusov’s classification system (Trusov, 1964; Chebanov, Galich, 2013) was used to identify the gonad maturity stages.

Non invasive ultrasound express examination was conducted in the frontal and transverse planes. During examination the transducer was pressed against the body in the region of the 3rd– 4th ventral scutes (counting from the pelvic fins), so that one edge of the transducer is located above the scutes (Chebanov, Galich, 2013). With frontal scanning, the skin, muscle tissue, serous membrane of the abdominal cavity, gonads, intestines were visualized. Male (testes) and female (ovaries) gonads were visualized depending on their echostructure. The testicular part was hyperechoic and has distinct margins. The fat part was under- or slightly developed from the medial side and practically not visible. The margins of the gonads were smoothly curved, while the bright hyperechoic tunic of the testis was clearly seen. The ovarian tissue appeared as a grainy “cloud-like” structure of mixed echogenicity with uneven boundaries without tunic. The fatty portion of the ovary was slight and was visualized in the shape of the darker areas as distinct from the lighter ovarian tissue (Chebanov, Galich, 2013).

Ultrasound examination was carried out with the help of MindrayDP-6600 ultrasound portable scanner by the author of the article who has been using ultrasound diagnostics for more than 5 years (more than 20 000 viewed sturgeons).

Formalin-preserved gonadal tissues were dehydrated in alcohol and embedded in paraffin. The embedded tissues were then sectioned at 5 µm and subsequently stained with hematoxylin and eosin (H&E).

A visual assessment of the gonads was performed in the euthanised fishes. The euthanasia of fish was carried out in accordance with the principles of humane treatment of animals. All sturgeon adults were destined for slaughter for commercial purposes.

#### 3.3. Scutes collection and measurement

Layer of dorsal, lateral and abdominal scutes was cut from head to dorsal fin of the euthanized sturgeons for studying the morphological parameters. The first dorsal scute was not cut due to extraction complexity. Small dorsal scutes after LSDR (last big scute of dorsal row) were not taken into account. After cutting the scutes were cleaned, photographed by digital SLR camera used in macro mode with enhanced flash (Fig. 2).

**Fig. 2.**
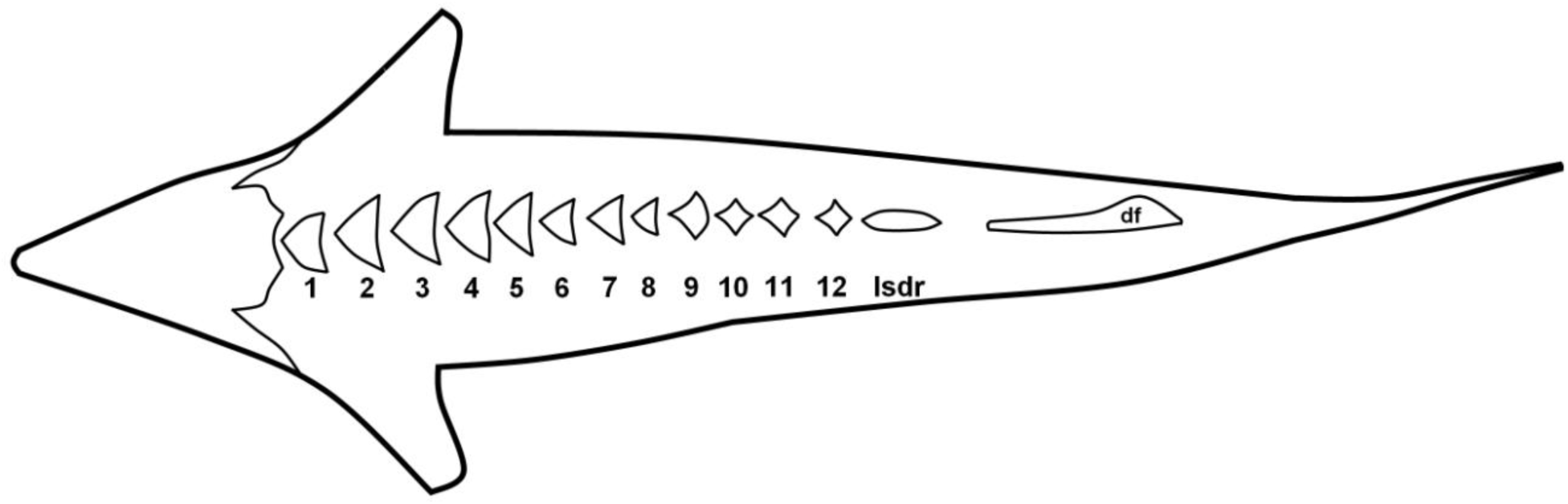
Numbering scheme of dorsal scutes. df - dorsal fin, lsdr- last big scute of dorsal row

Images were measured in ImageJ software using the following tools: “Straight Line”, “Segmented Line”, “Elliptical selections” and the Trust Canvas Widescreen Tablet. The following parameters of scutes were measured: the scute length; the scute width; left and right blade length of scute; the scute area; diameter of the conditional circle in which the scute was placed; the teeth number, the teeth length and the teeth width of scute (Fig.3).

**Fig. 3.**
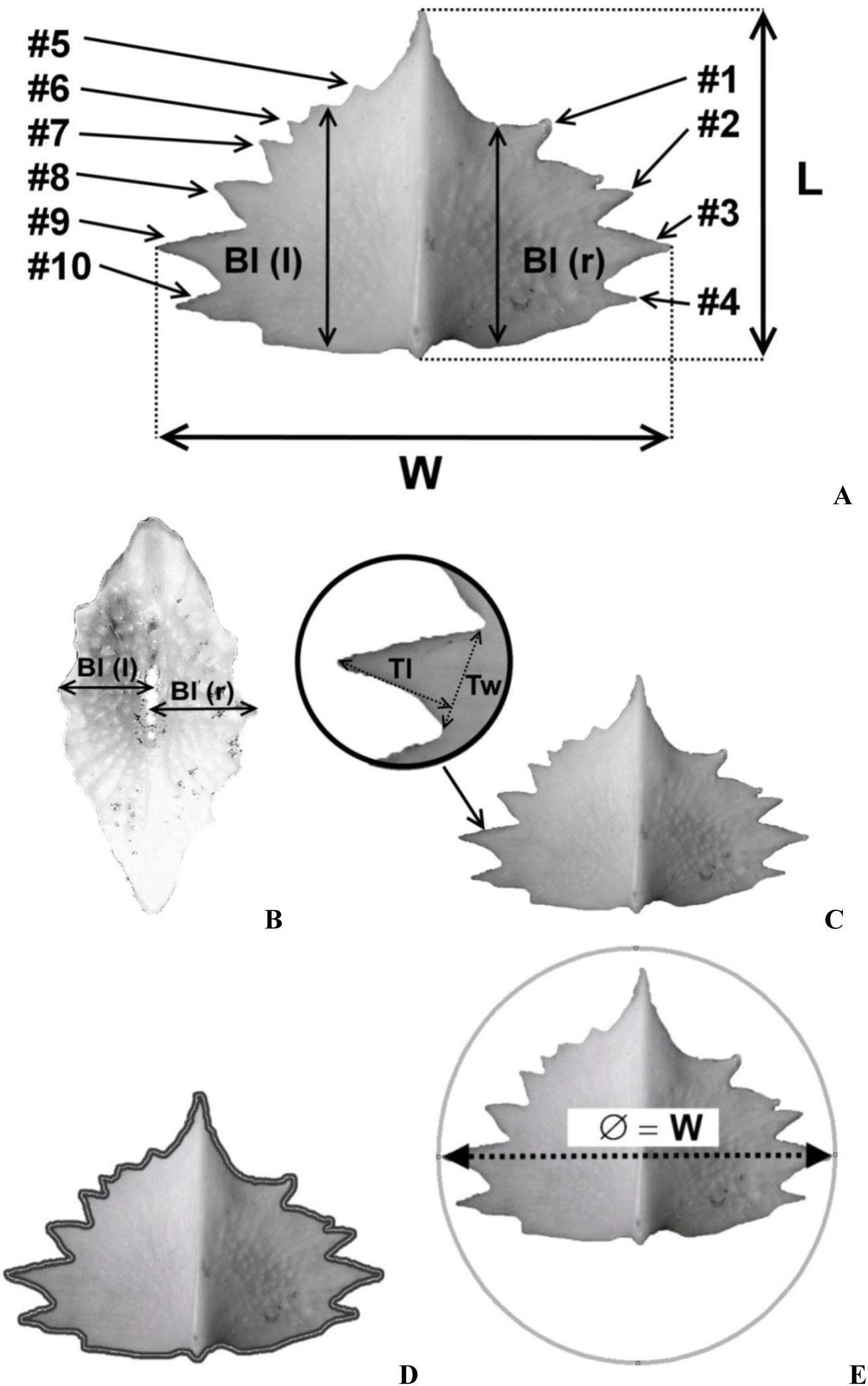
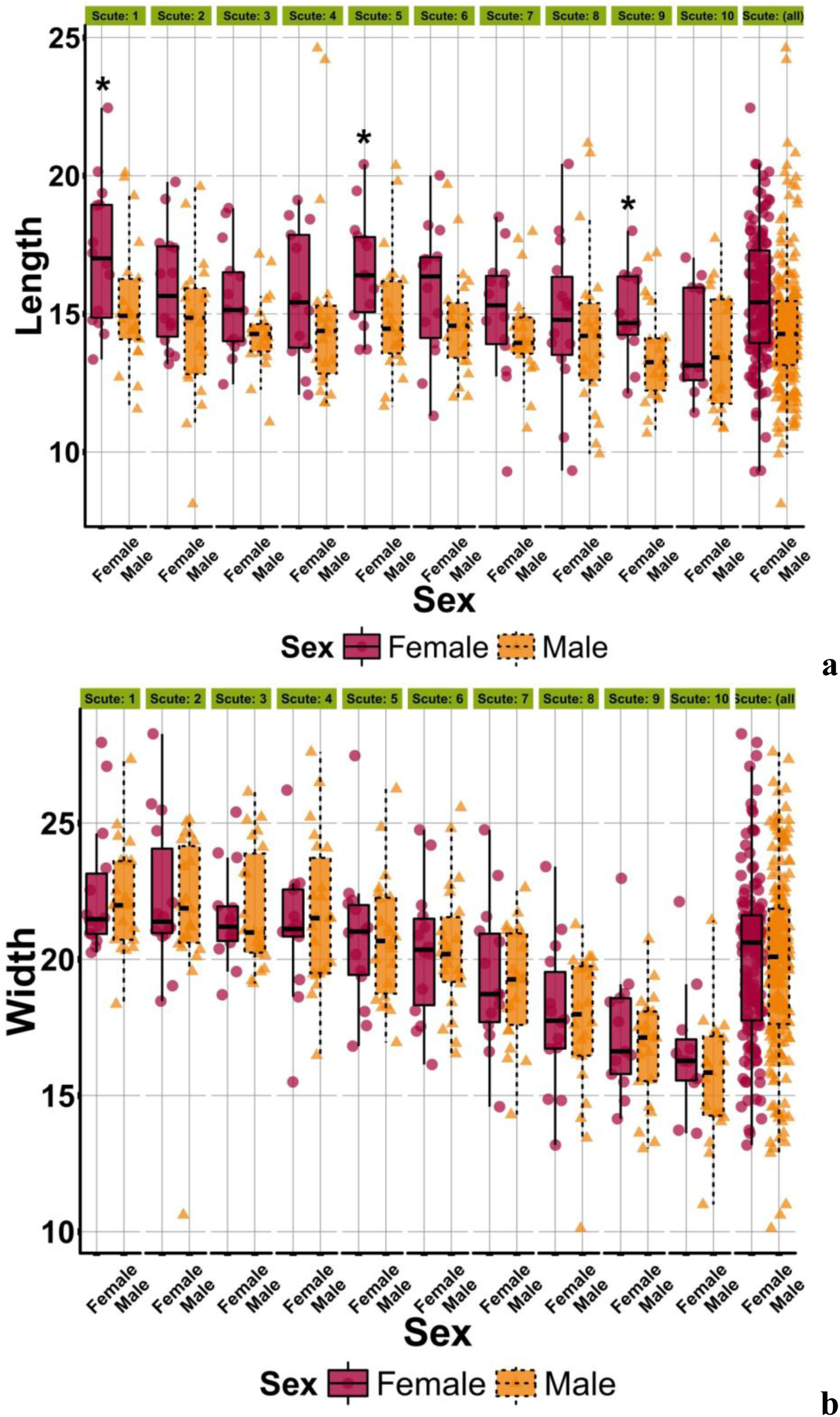
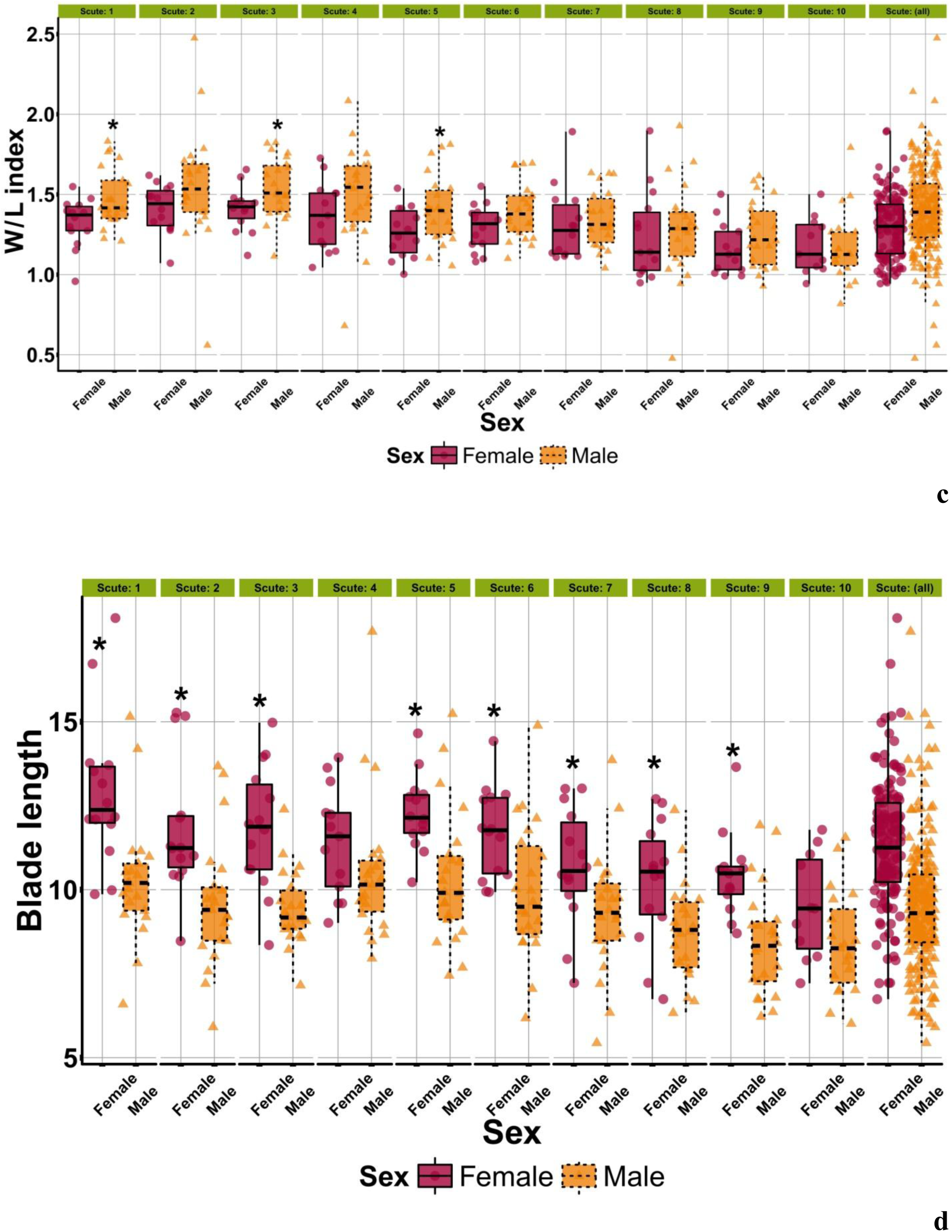
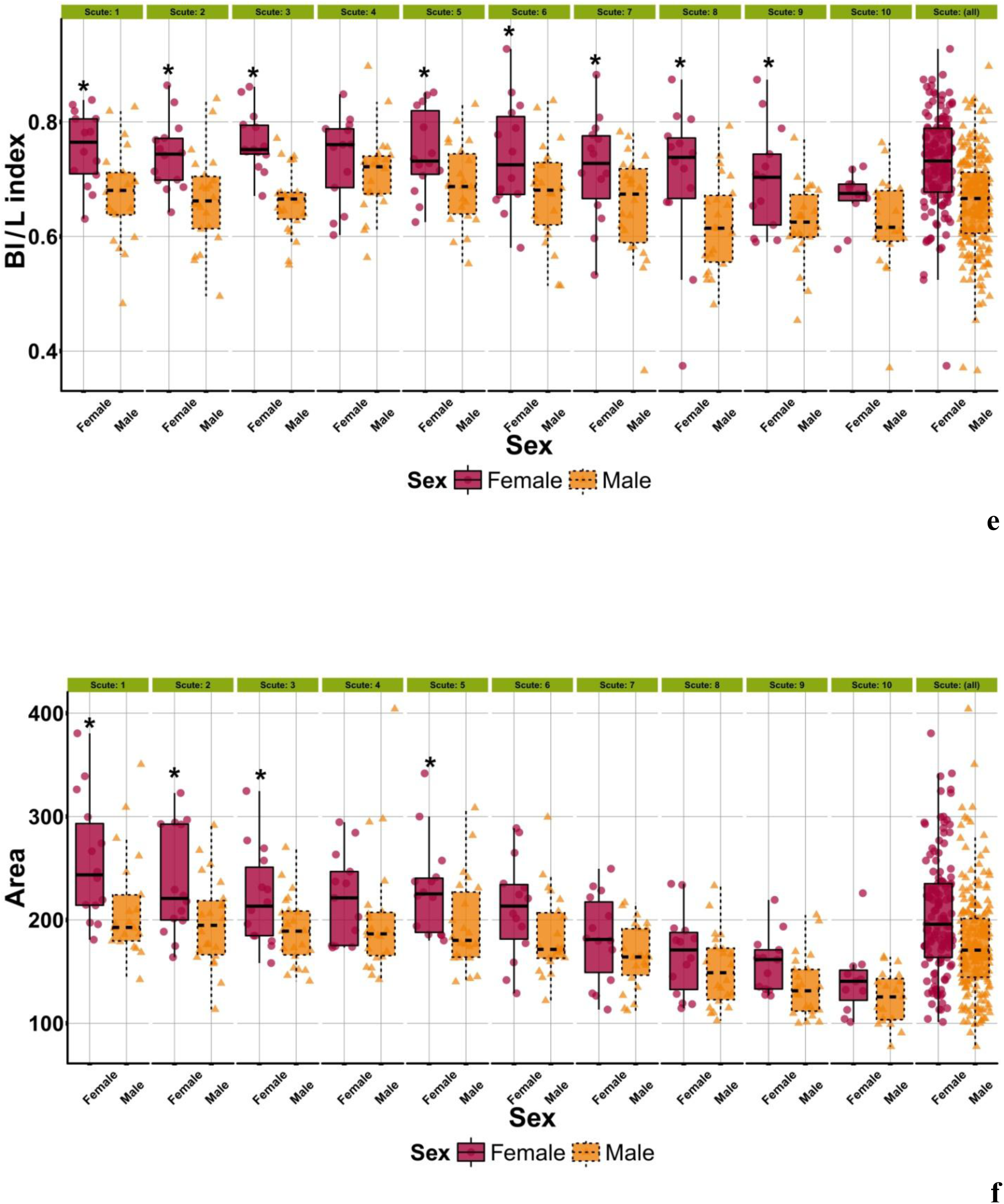
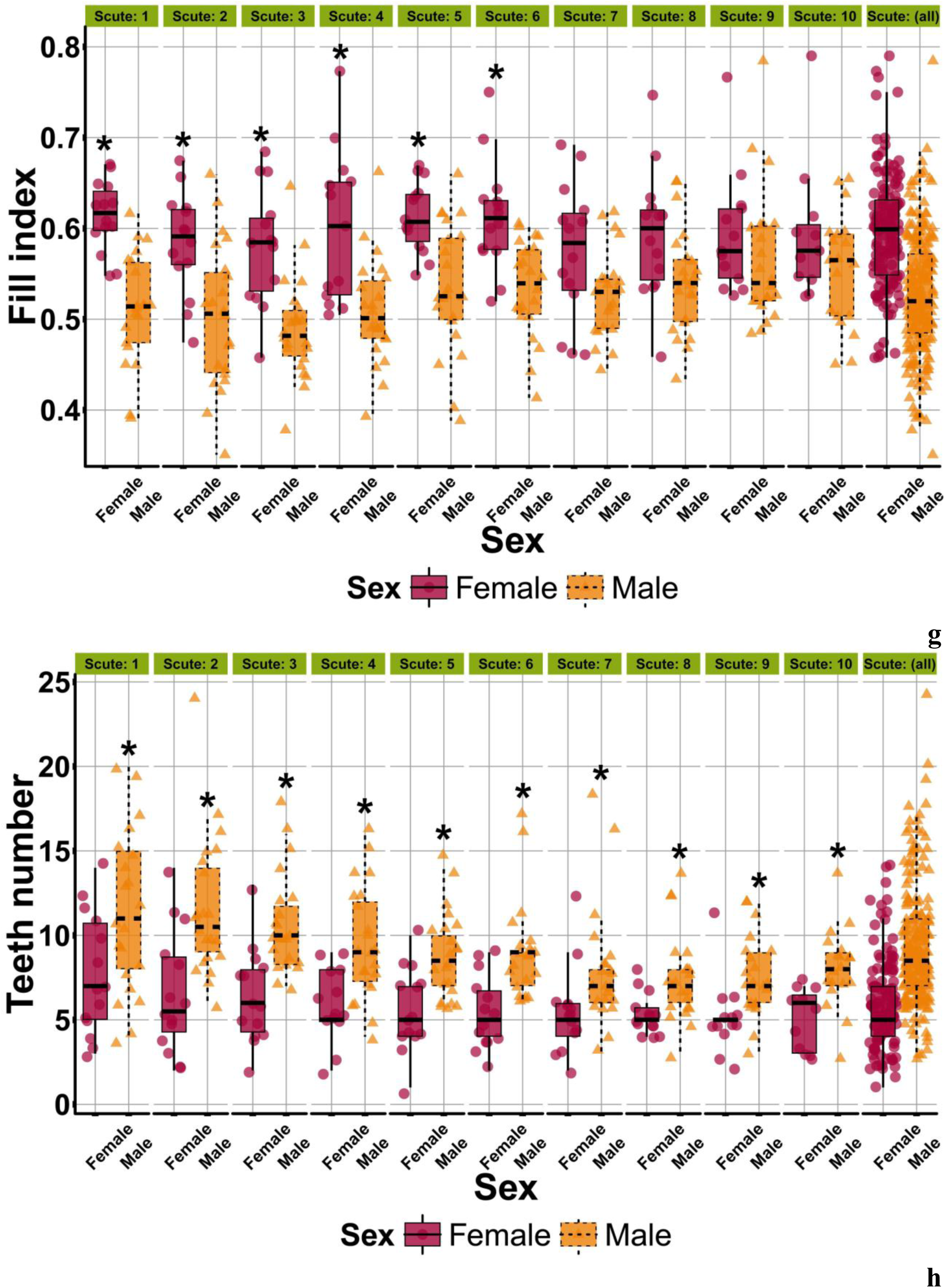
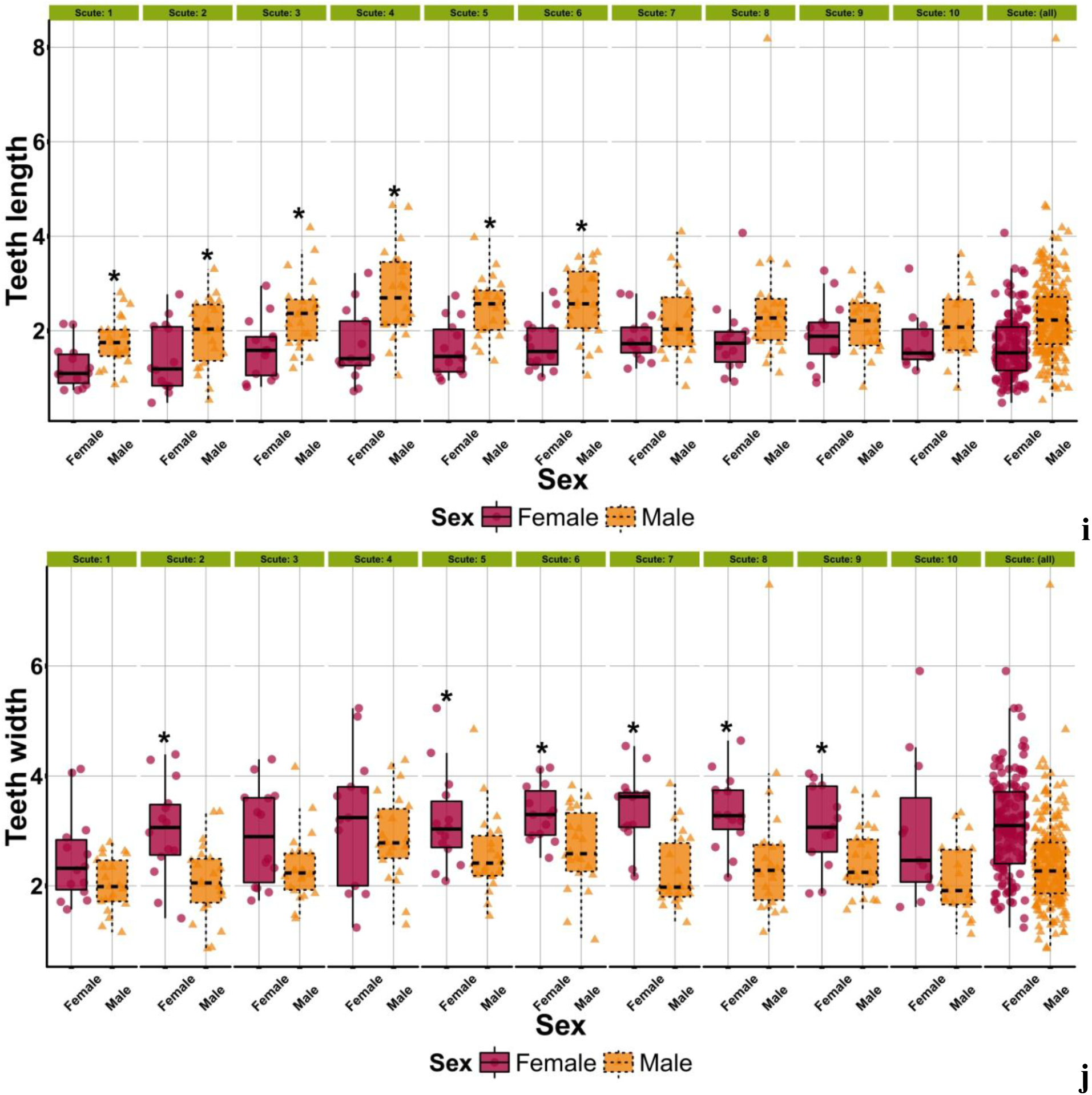
Measurement scheme of the sterlet dorsal scute using tool “Straight Line” of ImageJ software (A): “L” - scute length; “W” - scute width; “Bl(r)” - length of right blade; “Bl(l)” - length of left blade; #1-#10 – principle of scute teeth numbering. Measurement scheme of the last scute of dorsal row (B). Measurement scheme of the dorsal scute teeth (C). Measurement scheme of the dorsal scute area using tool “Segmented Line” of ImageJ (D). Measurement scheme of diameter of the conditional circle in which the scute was placed using tool “Elliptical selections” of ImageJ (E).

Based on the measurements obtained, the following indicators were calculated: scute width / scute length (W/L) index, average blade length / scute length (Bl/L) index, fill index (scute area / diameter of the conditional circle); teeth length / scute width (Tl/W) index, teeth length / teeth width (Tl/Tw) index.

After finding the sex specific structure of scutes, the morphological parameters of cleaned scutes were re-researched for juvenile and young sturgeon (Fig. 1).

### 4. Statistical analysis

The R (R Core Team, 2018) software was used for statistical processing of the results, including R packages: R Commander (Fox, 2005), PMCMR (Pohlert, 2014), MASS (Venables, Ripley, 2002), corrplot (Wei, Simko, 2016), randomForest (Liaw, Wiener, 2002), randomForestExplainer (Paluszynska, Biecek, 2017), ggplot2 (Wickham, 2016), circlize (Gu et al., 2014), reshape2 (Wickham, 2007), caret (Kuhn et al., 2017), vegan (Oksanen et al., 2017), rpart.plot (Milborrow et al., 2017), party (Hothorn et al., 2006), Boruta (Kursa, Rudnicki, 2010), neuralnet (Fritsch, Guenther, 2016). The distribution and homogeneity of data were evaluated using the Shapiro–Wilk, F– and Levene tests. To determine the statistical significance, parametrical (Student’s and Tukey’s tests) and nonparametrical (Mann-Whitney U– and Newman-Keuls tests) methods were used. To determine the presence of a relationship between several predictors (variables) the multicollinearity correlation matrix was used. To evaluate the qualitative characteristics the χ2 criterion (”chi-square”) was used.

For selected morphological parameters indices for sex determination: created sex determination models, the machine learning methods (neural networks, random forest method, boruta algorithm, building trees method, cumulative link model, binary matrix and binary discriminant analysis with DA algorithm) were used. The transition criterion from 1 in 0 in the binary matrix were the results obtained when using constructing trees method based on a recursive partition.

Random Forest implements Breiman’s random forest algorithm (based on Breiman and Cutler’s original FORTRAN code) for classification and regression can also be used in unsupervised mode for assessing proximities among data points (Liaw, Wiener, 2002).

Boruta is an all relevant feature selection wrapper algorithm, capable of working with any classification method that output variable importance measure (VIM); by default, Boruta uses Random Forest. The method performs a top-down search for relevant features by comparing original attributes’ importance with significance achievable at random, estimated using their permuted copies, and progressively eliminating irrelevant features to stabilise that test (Kursa, Rudnicki, 2010).

The party package aims at providing a recursive partitioning laboratory assembling various high- and low-level tools for building tree-based regression and classification models. This includes conditional inference trees (ctree), conditional inference forests (cforest) and parametric model trees (mob). At the core of the package is ctree, an implementation of conditional inference trees which embed tree-structured regression models into a well defined theory of conditional inference procedures. This non-parametric class of regression trees is applicable to all kinds of regression problems, including nominal, ordinal, numeric, censored as well as multivariate response variables and arbitrary measurement scales of the covariates (Hothorn et al., 2006).

Cumulative link models are powerful model class for such data since observations are treated rightfully as categorical, the ordered nature is exploited and the flexible regression framework allows in-depth analyses (Christensen, 2015).

## RESULTS

The sterlet sturgeon descendants from the Volga-river population and the siberian sturgeon descendants from the Lena-river population at the age of 3 months, 1 year and 2 years kept in aquaculture were studied. Various morphological and relative parameters (indices) that described the scutes form (structure) (Fig.1), were measured.

The average scute length in males decreases from 15.28±0.51 mm to 13.54± 0.38 mm; that in females decreases from 17.15±0.70 mm to 14.20±0.45 mm (Fig. 4a). Difference in the values of this parameter between scute nos. 1–10 in males and females varies in the range from 3.35 to 12.2 % (р < 0.05). The average scute width in males decreases from 22.29± 0.44 mm to 15.64±0.53 mm; that in females decreases from 22.48± 0.65 mm to 16.62± 0.56 mm (Fig. 4b). Difference in the values of this parameter between scute nos. 1–10 in males and females varies in the range from 0.8 to 6.3 % (р > 0.05). The average W/L index in males decreases from 1.55±0.08 to 1.17±0.05; that in females decreases from 1.41±0.04 to 1.16±0.05 (Fig. 4c). Difference in the values of this index between scute nos. 1–10 in males and females varies in the range from 0.80 to 11.50 % (р < 0.05). The average blade length of scute in males decreases from 10.49± 0.48 mm to 8.40± 0.36 mm; that in females decreases from 12.92±0.61 mm to 9.50 ± 0.33 mm (Fig. 4d). Difference in the values of this parameter between scute nos. 1–10 in males and females varies in the range from 9.0 to 25.8 % (р < 0.05). The average Bl/L index in males decreases from 0.71 ± 0.02 to 0.62 ± 0.02; that in females decreases from 0.77 ± 0.01 to 0.67± 0.01 (Fig. 4e); difference in the values of this index between males and females varies in the range from 3.9 to 15.7 % (р < 0.05). The average scute area in males decreases from 211.76±11.10 mm^2^ to 124.23 ± 5.93 mm^2^; that in females decreases from 256.82± 16.15 mm^2^ to 141.06± 6.95 mm^2^ (Fig. 4f); difference in the values of this parameter between males and females varies in the range from 8.8 to 21.3 % (р < 0.05). The average fill index in males varies in the range from 0.49 ± 0.01 to 0.57 ± 0.02; that in females varies in the range from 0.57 ± 0.02 to 0.61 ± 0.01 (Fig. 4g); difference in the values of this index between scute nos. 1–10 in males and females varies in the range from 4.7 to 19.8 % (р < 0.05). The average teeth number in males decreases from 11.55± 0.87 to 7.50± 0.55; that in females varies from 7.64± 0.96 to 5.00± 0.50 (Fig. 4h). Difference in the values of this parameter between scute nos. 1–10 in males and females varies in the range from 33.9 to 77.6 % (р < 0.05). The average teeth length in males varies in the range from 1.77± 0.12 mm to 2.81± 0.21 mm; that in females varies in the range from 1.24± 0.12 mm to 1.93± 0.22 mm (Fig. 4i). Difference in the values of this parameter between scute nos. 1–10 in males and females varies in the range from 12.7 to 68.8 % (р < 0.05). The average teeth width in males varies in the range from 2.03± 0.11 mm to 2.86± 0.17 mm; that in females varies in the range from 2.50± 0.22 mm to 3.40± 0.18 mm (Fig. 4j). Difference in the values of this parameter between scute nos. 1–10 in males and females varies in the range from 13.0 to 51.8 % (р < 0.05). The average Tl/W index in males increases from 0.08 ± 0.01 to 0.18 ± 0.01; that in females increases from 0.05 ± 0.01 to 0.11 ± 0.01 (Fig. 4k); difference in the values of this index between males and females varies in the range from 0.5 to 66.2 % (р < 0.05). The average Tl/Tw index in males varies in the range from 0.89 ± 0.06 to 1.06 ± 0.09; that in females varies in the range from in the range 0.46 ± 0.05 to 0.65 ± 0.09 (Fig. 4l). Difference in the values of this index between males and females varies in the range from 43.0 to 105.4 % (р < 0.05).

**Fig. 4.**
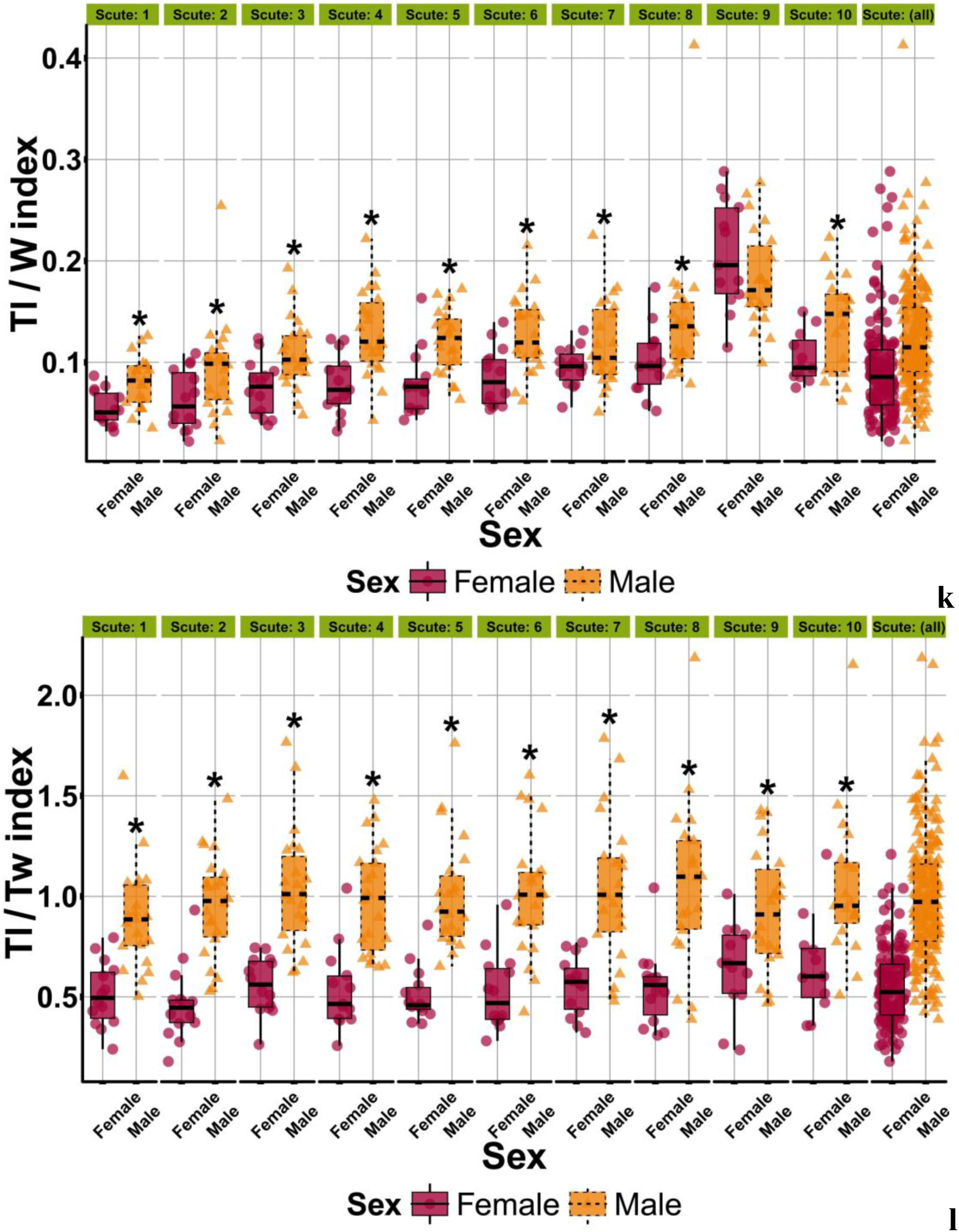
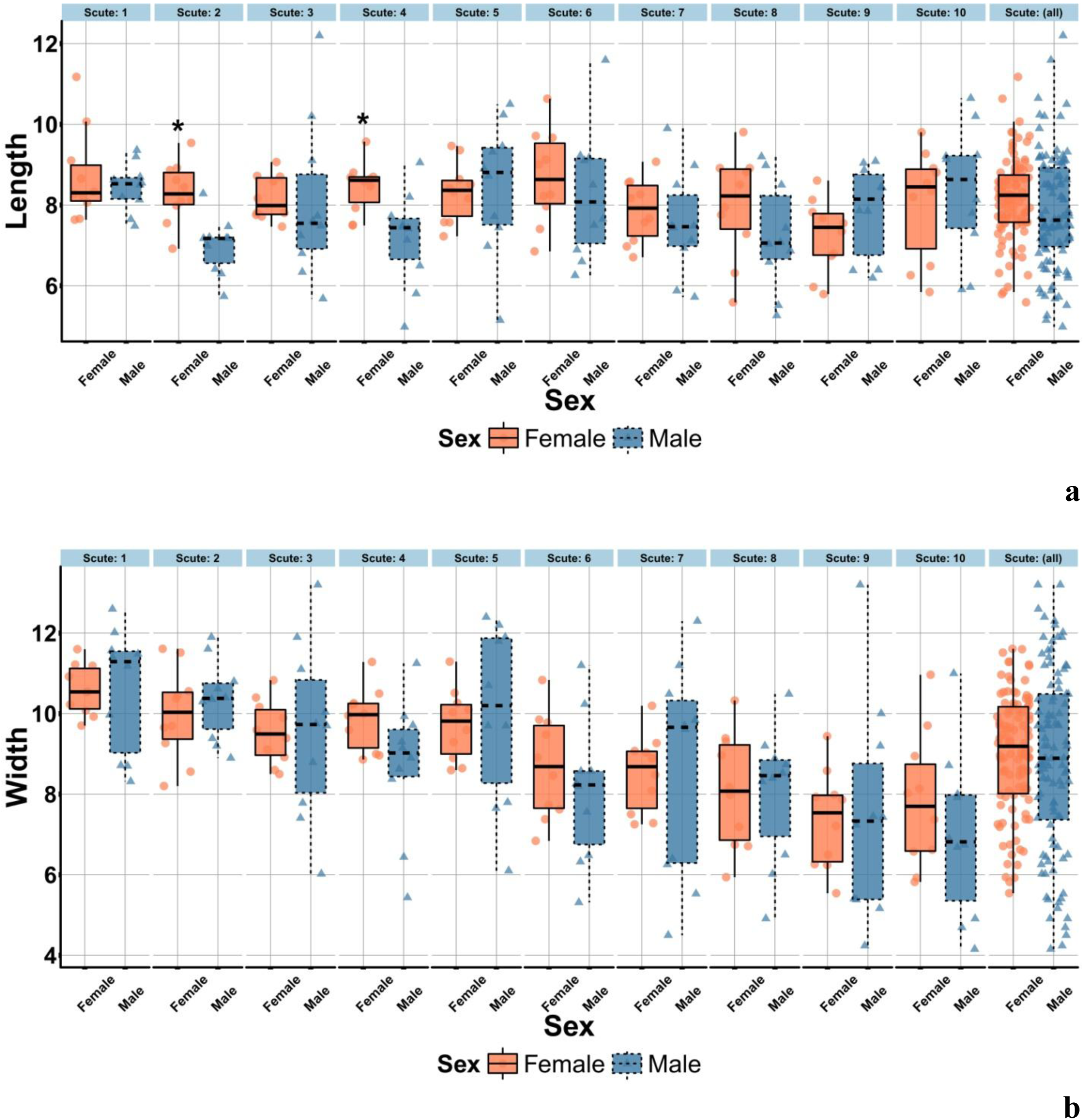
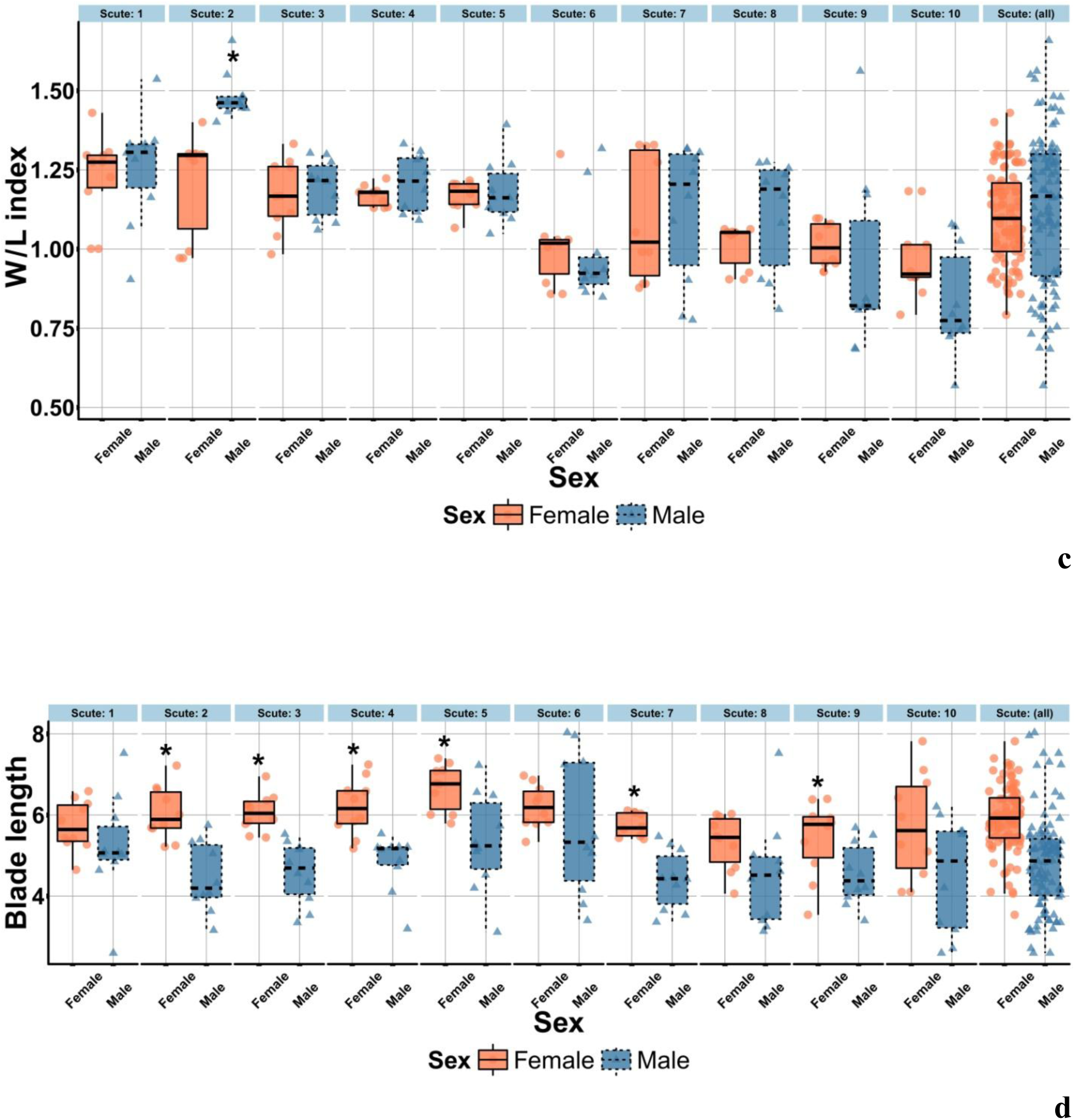
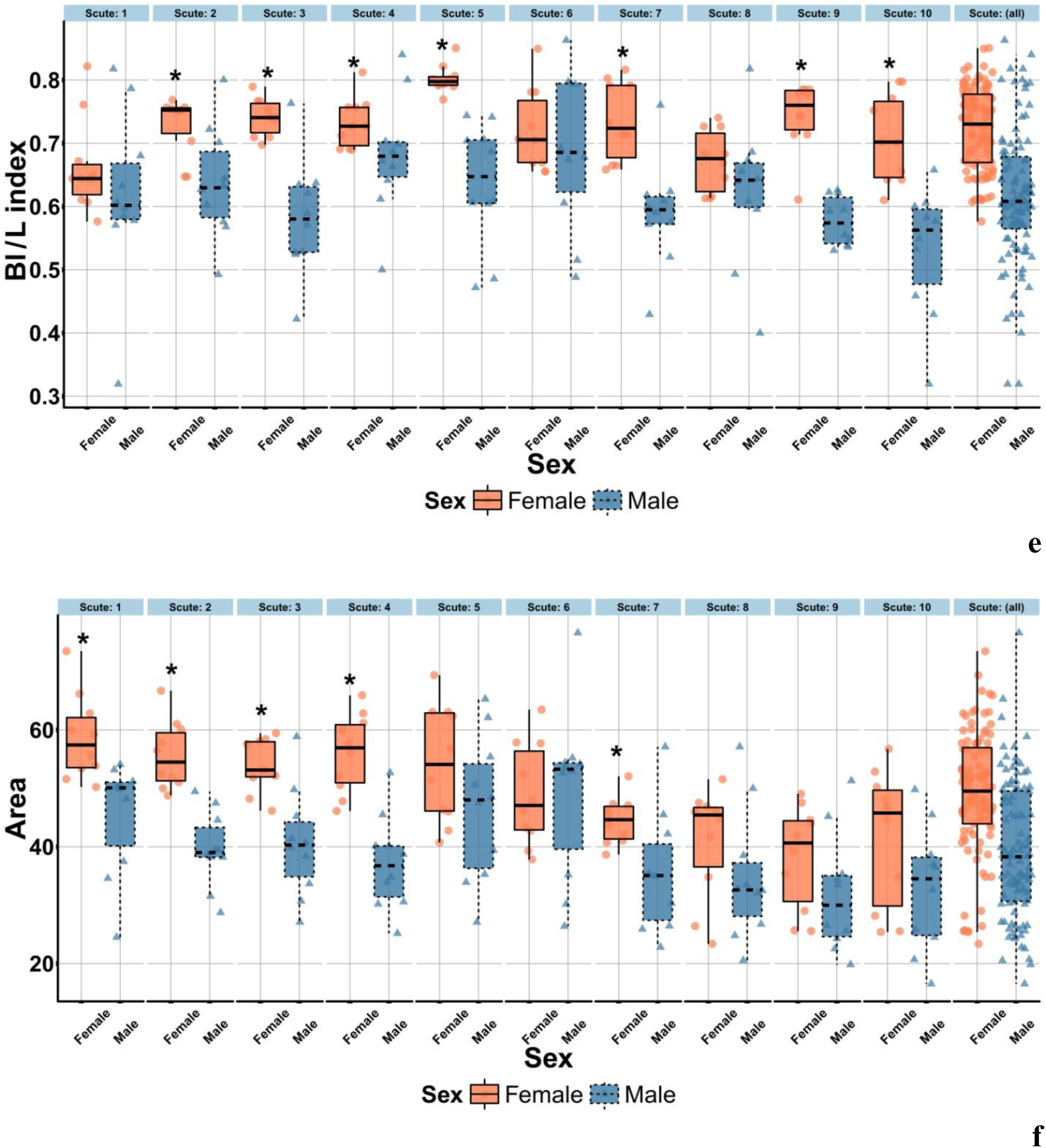
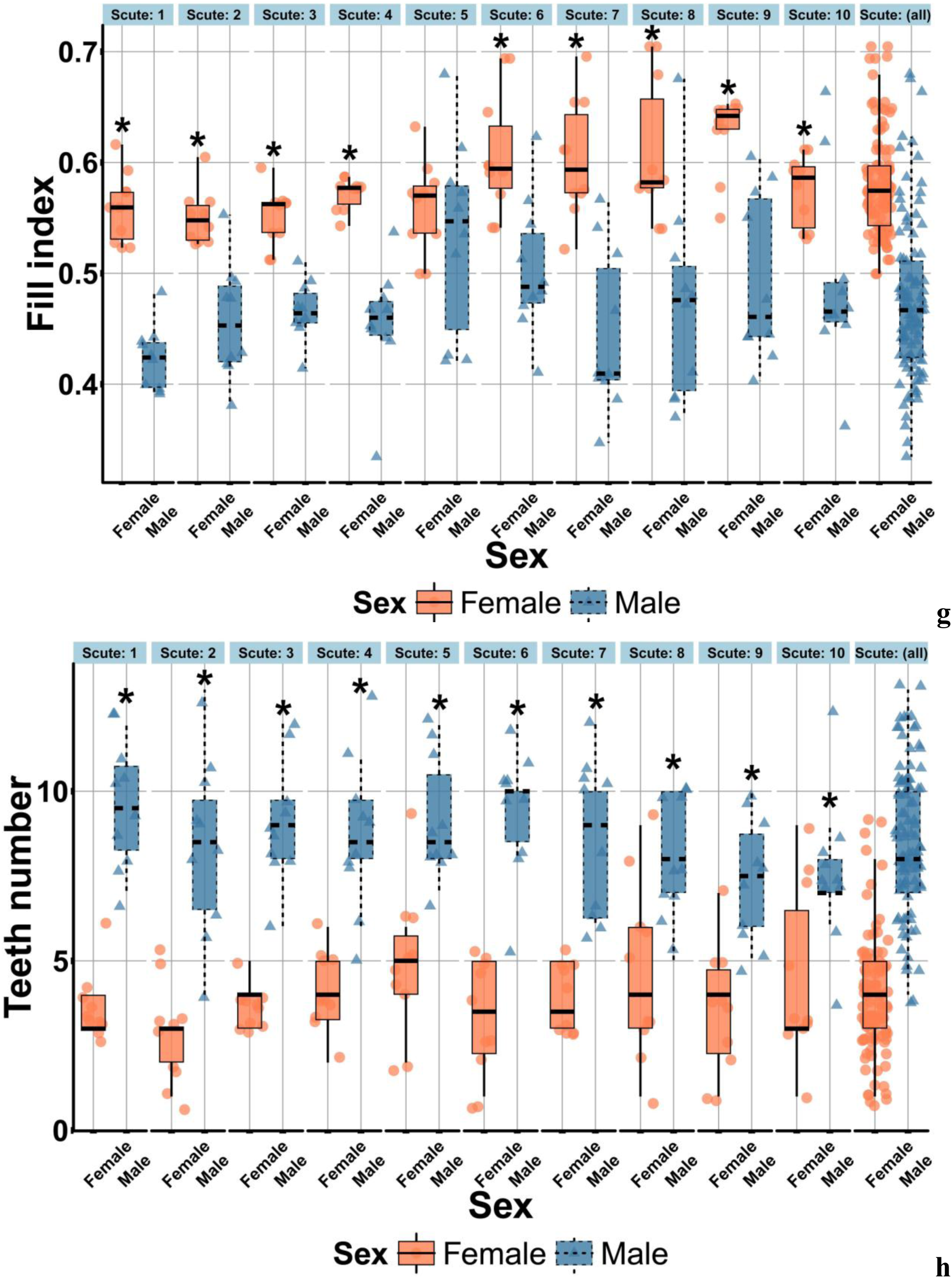
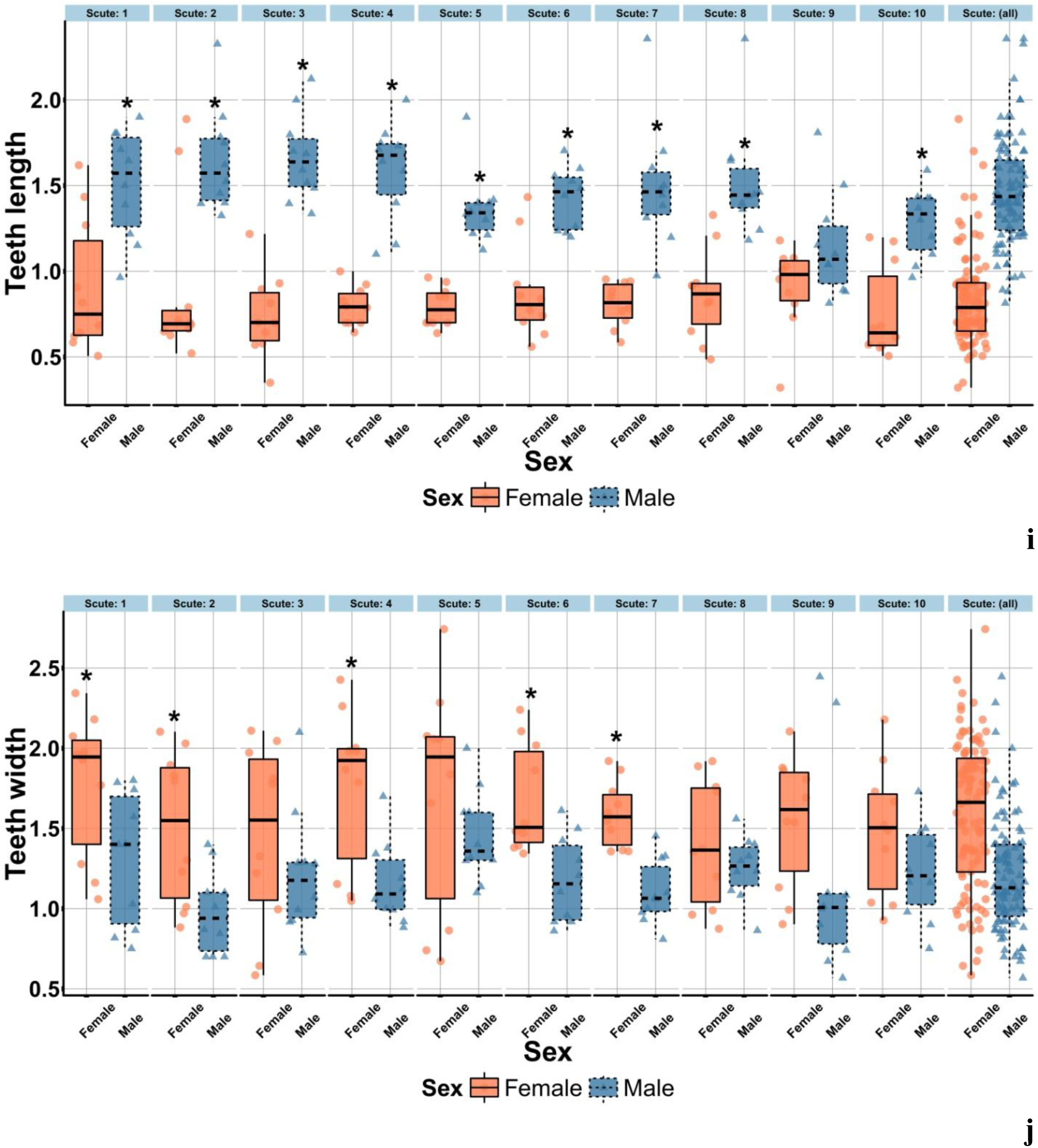
Statistical sex morphological differences between the dorsal scutes of adults *Acipenser ruthenus*. Boxplot with jitter show the sex morphological differences between median, percentile boundaries (0.25 and 0.75), interquartile range (IQR), min (max x1, Q3 + 1.5 x IQR), min (max x1, Q1 - 1.5 x IQR) and data distribution of the scute length (a), the scute width (b), the W/L index (c), the blade length of scute (d), the Bl/L index (e), the scute area (f), the fill index (g), the teeth number (h), the teeth length (i), the teeth width (j), the Tl/W index (k), the Tl/Tw index (l). Symbol (*) shows the statistical significance (p < 0.05) (Mann-Whitney U–test). Number of fish treated = 37. Number of scutes treated = 370.

The average scute length in males of young decreases from 8.44 ± 0.19 mm to 6.99± 0.22mm; that in females of young decreases from 8.72±0.36mm to 7.24±0.29 mm (Fig. 5a). Difference in the values of this parameter between scute nos. 1–10 in males and females of young varies in the range from 1.4 to 18.8 % (р < 0.05). The average scute width in males of young decreases from 10.60± 0.49mm to 6.97±0.66 mm; that in females of young decreases from 22.48± 0.65 mm to 16.62± 0.56 mm (Fig. 5b). Difference in the values of this parameter between scute nos. 1–10 in males and females of young varies in the range from 0.8 to 6.3 % (р > 0.05). The average W/L index in males of young decreases from 1.48±0.02 to 0.83±0.05; that in females of young decreases from 1.23±0.04 to 0.97±0.04 (Fig. 5c). Difference in the values of this index between scute nos. 1–10 in males and females varies in the range from 1.08 to 22.39 % (р < 0.05). The average blade length of scute in males of young decreases from 5.76± 0.55 mm to 4.41± 0.23 mm; that in females of young decreases from 6.67±0.18 mm to 5.32± 0.21 mm (Fig. 5d). Difference in the values of this parameter between scute nos. 1–10 in males and females of young varies in the range from 7.7 to 35.4 % (р < 0.05). The average Bl/L index in males of young decreases from 0.69± 0.04 to 0.53± 0.03; that in females of young decreases from 0.80± 0.01 to 0.66± 0.02 (Fig. 5e); difference in the values of this index between males and females of young varies in the range from 4.3 to 33.2 % (р < 0.05). The average scute area in males of young decreases from 49.15±4.67mm^2^ to 31.84±3.22 mm^2^; that in females of young decreases from 58.65± 2.30 mm^2^ to 38.22± 2.80mm^2^ (Fig. 5f); difference in the values of this parameter between males and females of young varies in the range from 0.6 to 50.7 % (р < 0.05). The average fill index in males of young varies in the range from 0.53± 0.03 to 0.42± 0.01; that in females of young varies in the range from 0.63± 0.01 to 0.55± 0.01 (Fig. 5g); difference in the values of this index between scute nos. 1–10 in males and females of young varies in the range from 5.0 to 35.4 % (р < 0.05). The average teeth number in males of young decreases from 9.60±0.54 to 7.40± 0.60; that in females of young decreases from 4.80± 0.65 to 2.80± 0.44 (Fig. 5h). Difference in the values of this parameter between scute nos. 1–10 in males and females of young varies in the range from 66.7 to 200.0 % (р < 0.05). The average teeth length in males of young varies in the range from 1.66± 0.08 mm to 1.15±0.10 mm; that in females of young varies in the range from 0.91± 0.08 mm to 0.74± 0.08 mm (Fig. 5i). Difference in the values of this parameter between scute nos. 1–10 in males and females of young varies in the range from 27.3 to 124.7 % (р < 0.05). The average teeth width in males of young varies in the range from 1.45±0.09 mm to 0.98± 0.08 mm; that in females of young varies in the range from 1.77± 0.14 mm to 1.41± 0.13mm (Fig. 5j). Difference in the values of this parameter between scute nos. 1–10 in males and females of young varies in the range from 12.2 to 54.0 % (р < 0.05). The average Tl/W index in males of young increases from 0.14± 0.01 to 0.20± 0.01; that in females of young increases from 0.08 ± 0.01 to 0.12 ± 0.01 (Fig. 5k); difference in the values of this index between males and females of young varies in the range from 33.1 to 129.7 % (р < 0.05). The average Tl/Tw index in males of young varies in the range from 0.95± 0.04 to 1.75± 0.11; that in females of young varies in the range from in the range 0.49± 0.05 to 0.69± 0.13 (Fig. 5l). Difference in the values of this index between males and females of young varies in the range from 67.6 to 192.5 % (р < 0.05).

**Fig. 5.**
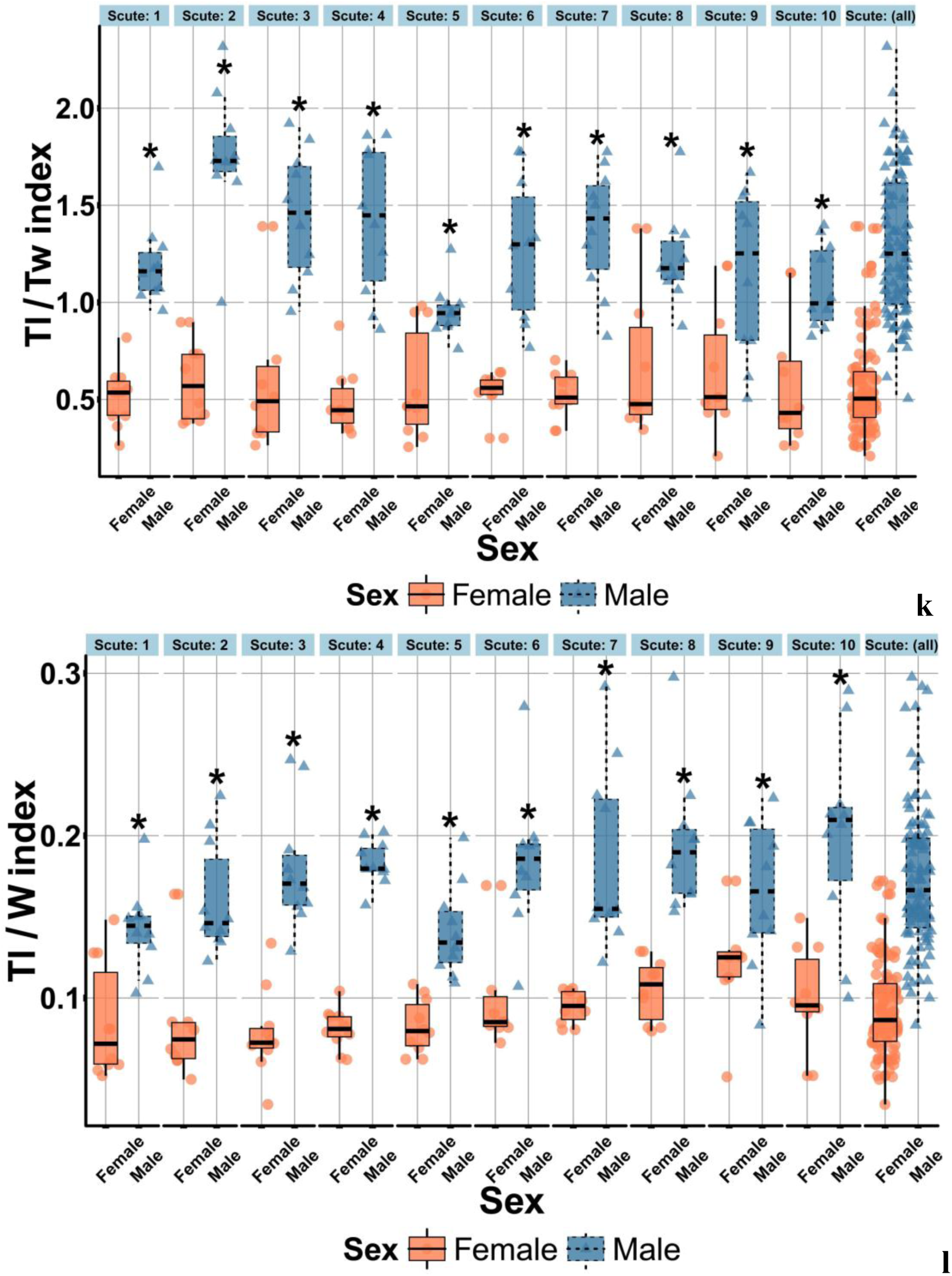
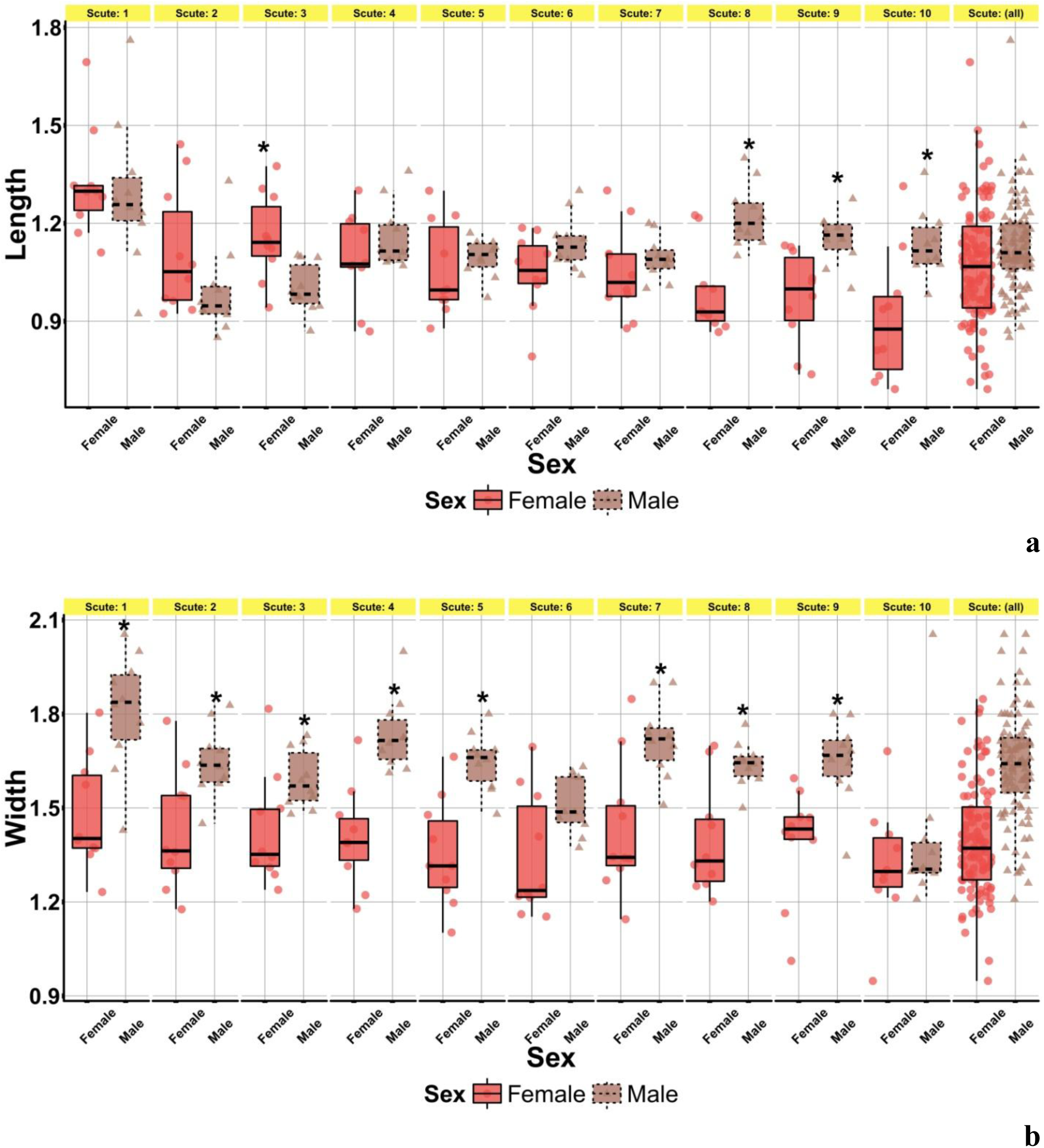
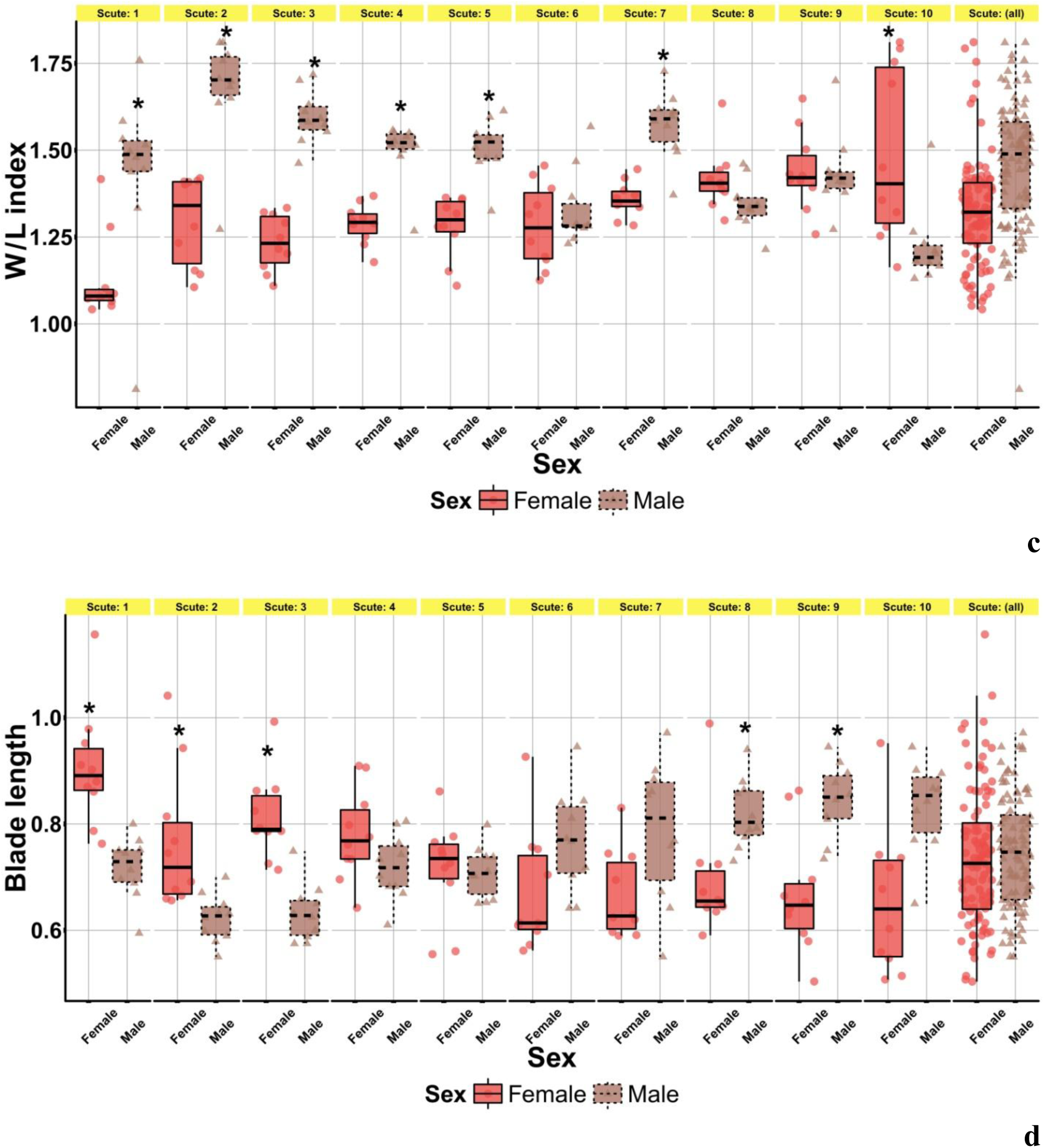
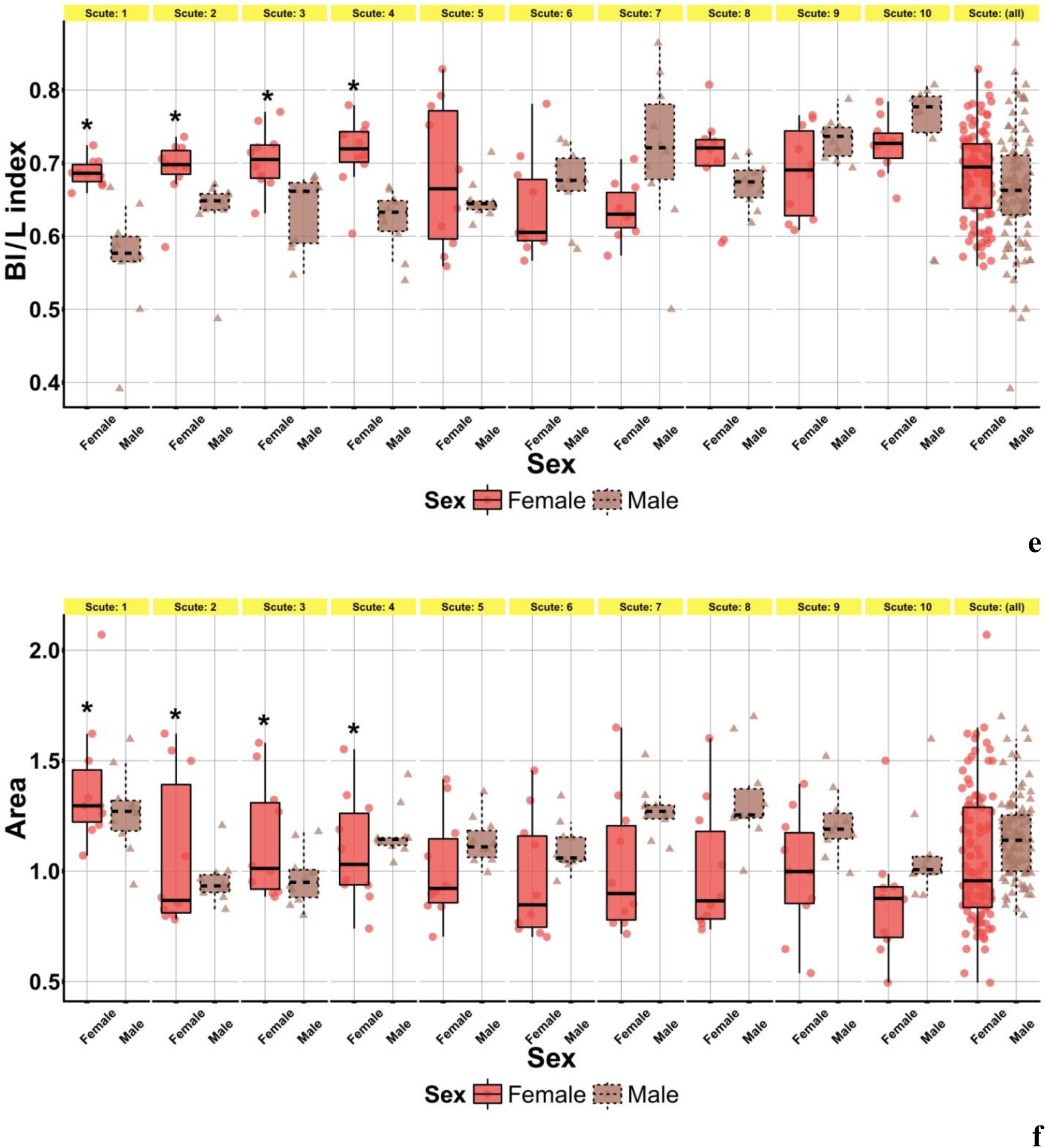
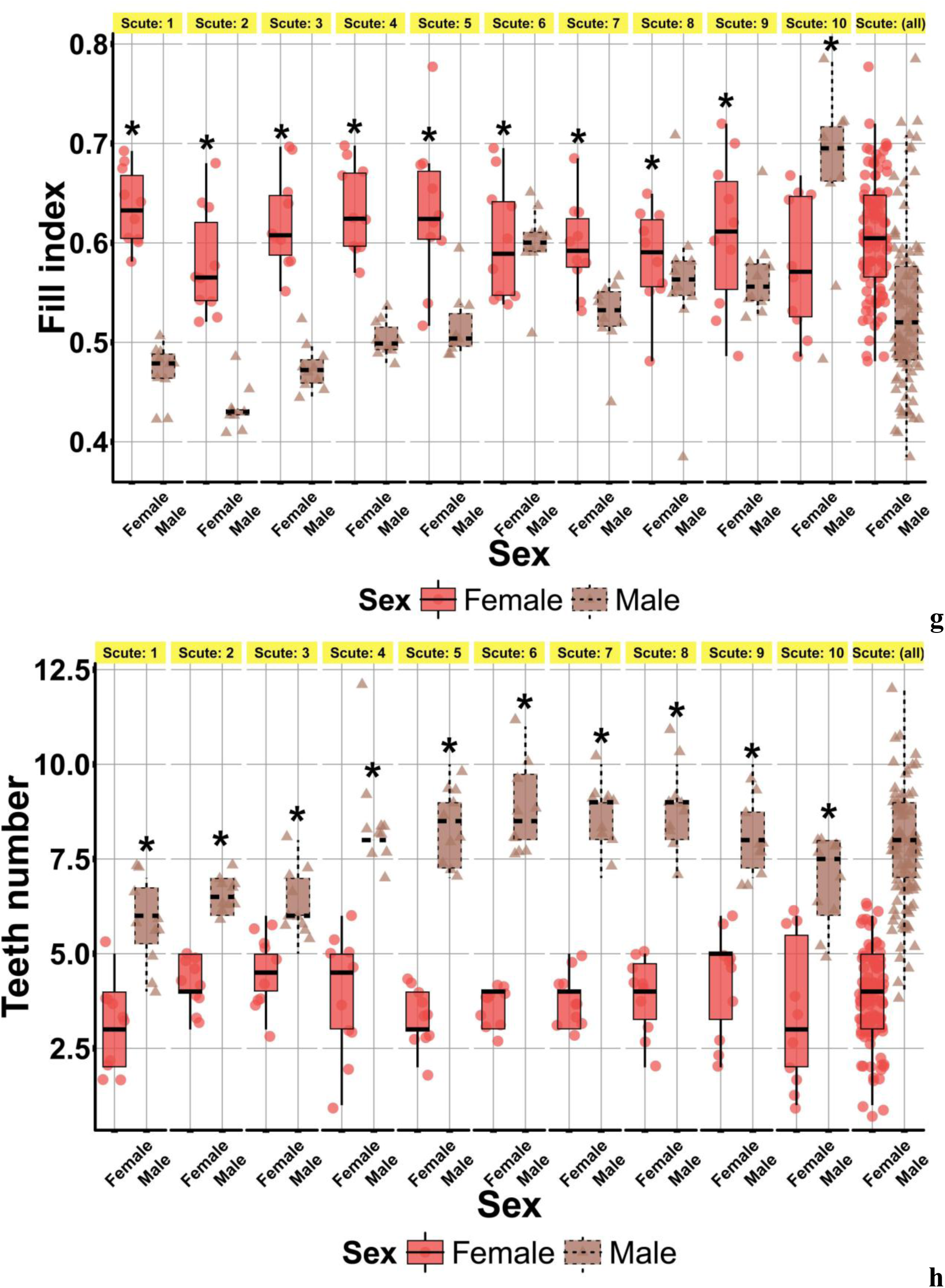
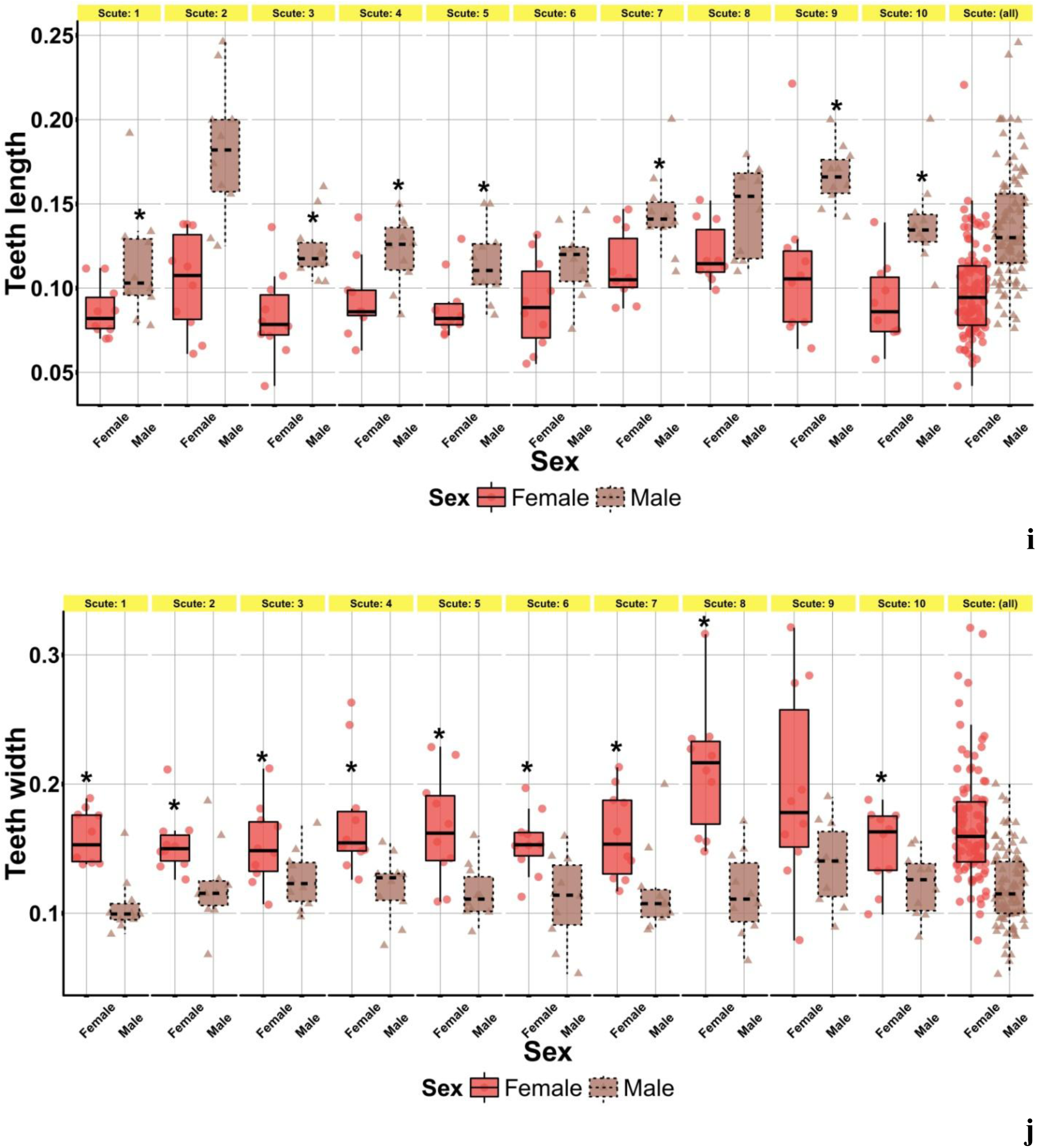
Statistical sex morphological differences between the dorsal scutes of young *Acipenser ruthenus*. Boxplot with jitter show the sex morphological differences between median, percentile boundaries (0.25 and 0.75), interquartile range (IQR), min (max x1, Q3 + 1.5 x IQR), min (max x1, Q1 - 1.5 x IQR) and data distribution of the scute length (a), the scute width (b), the W/L index (c), the blade length of scute (d), the Bl/L index (e), the scute area (f), the fill index (g), the teeth number (h), the teeth length (i), the teeth width (j), the Tl/W index (k), the Tl/Tw index (l). Symbol (*) shows the statistical significance (p < 0.05) (Mann-Whitney U–test). Number of fish treated = 20. Number of scutes treated = 200.

The average scute length in males of juveniles decreases from 1.29± 0.07mm to 0.99± 0.04mm; that in females of juveniles decreases from 1.32±0.05 mm to 0.91±0.06 mm (Fig. 6a). Difference in the values of this parameter between scute nos. 1–10 in males and females of juveniles varies in the range from 2.4 to 25.3 % (р < 0.05). The average scute width in males of juveniles decreases from 1.81± 0.06 mm to 1.39±0.08 mm; that in females of juveniles decreases from 1.48± 0.06 mm to 1.32± 0.06 mm (Fig. 6b). Difference in the values of this parameter between scute nos. 1–10 in males and females of juveniles varies in the range from 5.7 to 23.4 % (р < 0.05). The average W/L index in males of juveniles decreases from 1.68±0.05 to 1.22±0.03; that in females of juveniles decreases from 1.49±0.08 to 1.13±0.04 (Fig. 6c). Difference in the values of this index between scute nos. 1–10 in males and females varies in the range from 0.3 to 29.5 % (р < 0.05). The average blade length of scute in males of juveniles decreases from 0.84±0.02 mm to 0.62± 0.01 mm; that in females of juveniles decreases from 0.91±0.03 mm to 0.66±0.04 mm (Fig. 6d). Difference in the values of this parameter between scute nos. 1–10 in males and females of juveniles varies in the range from 0.67 to 28.65 % (р < 0.05). The average Bl/L index in males of juveniles varies in the range from 0.74 ± 0.03 to 0.57 ± 0.02; that in females of juveniles varies in the range from 0.72 ± 0.01 to 0.64 ± 0.02 (Fig. 6e); difference in the values of this index between males and females of juveniles varies in the range from 1.8 to 21.0 % (р < 0.05). The average scute area in males of juveniles decreases from 1.32 ± 0.07 mm^2^ to 0.96 ± 0.03 mm^2^; that in females of juveniles decreases from 1.38 ± 0.09 mm^2^ to 0.86 ± 0.09 mm^2^ (Fig. 6f); difference in the values of this parameter between males and females of juveniles varies in the range from 7.2 to 32.2 % (р < 0.05). The average fill index in males of juveniles varies in the range from 0.67 ± 0.03 to 0.43 ± 0.01; that in females of juveniles varies in the range from 0.64 ± 0.01 to 0.58± 0.02 (Fig. 6g); difference in the values of this index between scute nos. 1–10 in males and females of juveniles varies in the range from 0.3 to 35.1 % (р < 0.05). The average teeth number in males of juveniles decreases from 8.90 ± 0.35 to 5.80 ± 0.36; that in females of juveniles decreases from 4.60 ± 0.31 to 3.10 ± 0.35 (Fig. 6h). Difference in the values of this parameter between scute nos. 1–10 in males and females of juveniles varies in the range from 39.1 to 151.5 % (р < 0.05). The average teeth length in males of juveniles varies in the range from 0.18 ± 0.01 mm to 0.11 ±0.01 mm; that in females of juveniles varies in the range from 0.12± 0.01 mm to 0.08± 0.01 mm (Fig. 6i). Difference in the values of this parameter between scute nos. 1–10 in males and females of juveniles varies in the range from 20.9 to 75.4 % (р < 0.05). The average teeth width in males of juveniles varies in the range from 0.14 ± 0.01 mm to 0.11± 0.01 mm; that in females of juveniles varies in the range from 0.21 ± 0.02 mm to 0.15± 0.01 mm (Fig. 6j). Difference in the values of this parameter between scute nos. 1–10 in males and females of juveniles varies in the range from 20.9 to 83.3 % (р < 0.05). The average Tl/W index in males of juveniles varies in the range from 0.06± 0.01 to 0.11± 0.01; that in females of juveniles varies in the range from 0.06 ± 0.01 to 0.09 ± 0.01 (Fig. 6k); difference in the values of this index between males and females of juveniles varies in the range from 2.1 to 50.8 % (р < 0.05). The average Tl/Tw index in males of juveniles varies in the range from 1.64 ± 0.21 to 0.99± 0.05; that in females of juveniles varies in the range from in the range 0.72± 0.05 to 0.55±0.03 (Fig. 6l). Difference in the values of this index between males and females of juveniles varies in the range from 78.4 to 140.0 % (р < 0.05).

**Fig. 6.**
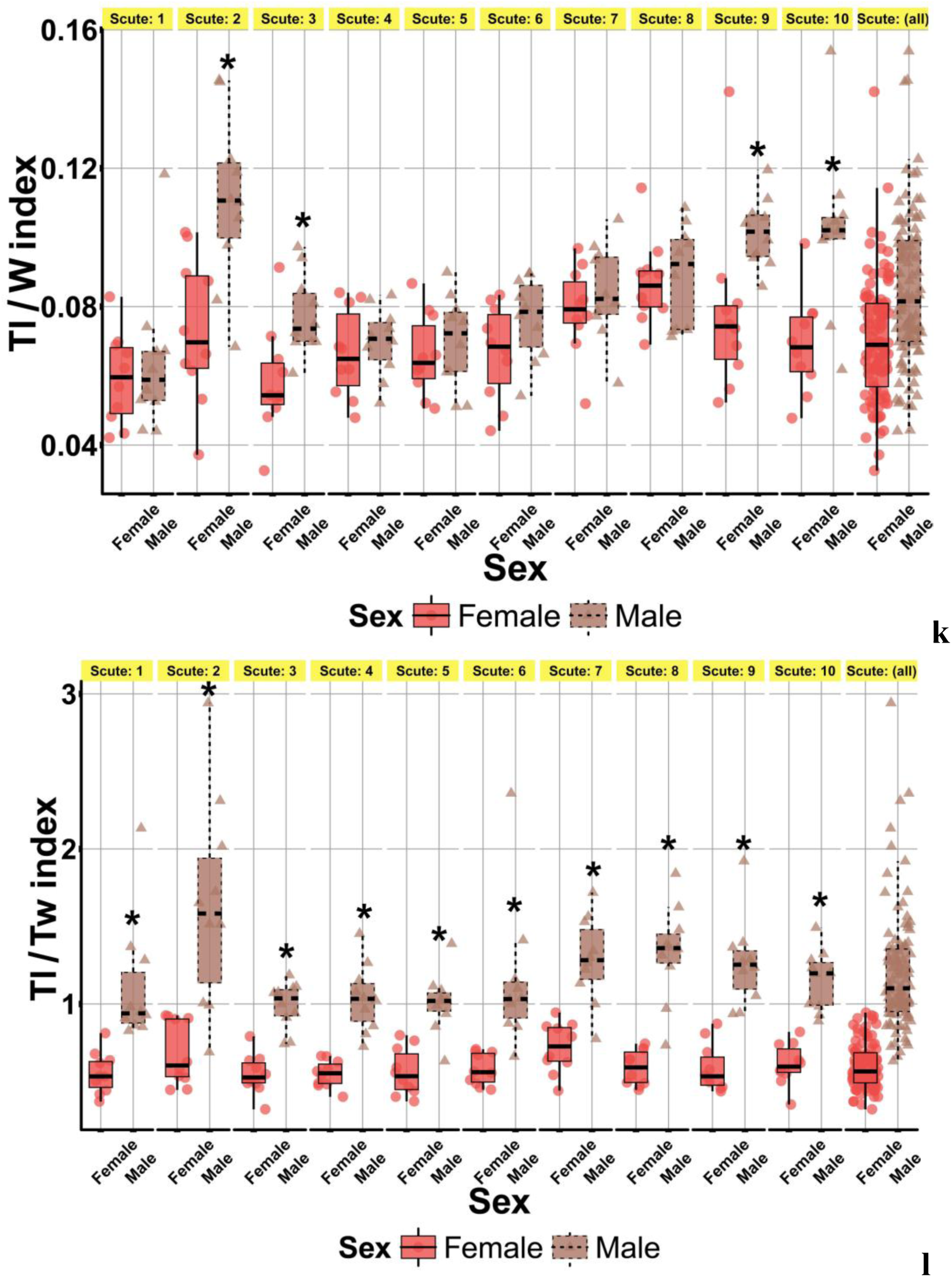
Statistical sex morphological differences between the dorsal scutes of juveniles *Acipenser ruthenus*. Boxplot with jitter show the sex morphological differences between median, percentile boundaries (0.25 and 0.75), interquartile range (IQR), min (max x1, Q3 + 1.5 x IQR), min (max x1, Q1 - 1.5 x IQR) and data distribution of the scute length (a), the scute width (b), the W/L index (c), the blade length of scute (d), the Bl/L index (e), the scute area (f), the fill index (g), the teeth number (h), the teeth length (i), the teeth width (j), the Tl/W index (k), the Tl/Tw index (l). Symbol (*) shows the statistical significance (p < 0.05) (Mann-Whitney U–test). Number of fish treated = 20. Number of scutes treated = 200.

We found the correlation in morphological parameters and indices in adult sterlet (Fig. 7, 10). Strong correlation (from ±0.7 to ±1) found between the scute area and the blade length of scute, between the scute area and the scute length, between the blade length of scute and the scute length in adults of males and females.

**Fig. 7.**
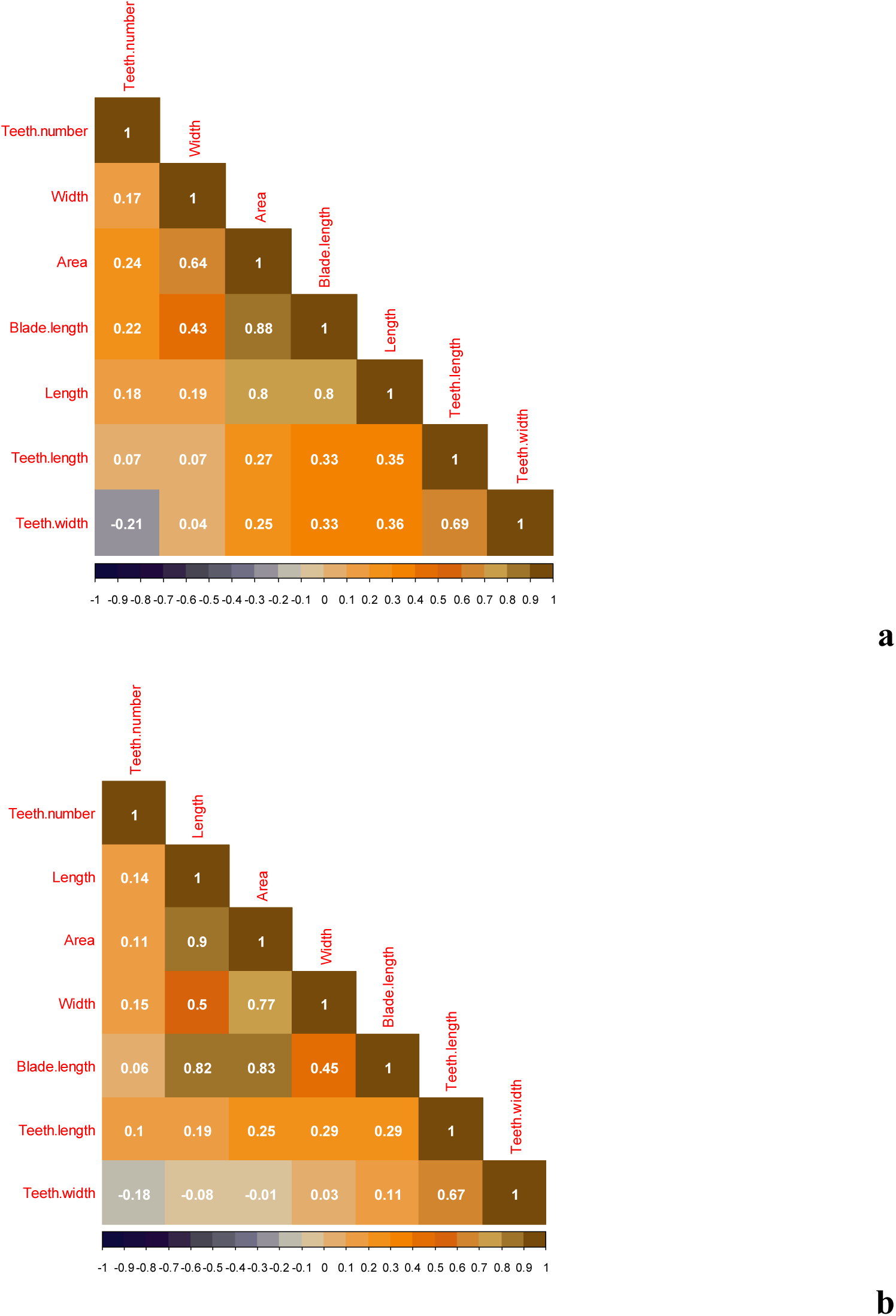
Correlation matrix show multicollinearity between morphological parameters of the first to fifth dorsal scutes of adults males (a) and females (b) *Acipenser ruthenus* (fish length – 61.2±1.3 cm, 2 years-old). Number of fish treated = 37. Number of scutes treated = 185.

We found the correlation in morphological parameters and indices in young sterlet (Fig. 8, 10). Strong correlation found between the scute area and the blade length of scute, between the scute area and the scute length, between the blade length of scute and the scute length, between the scute width and the scute length in young of males. Strong correlation found between the scute area and the scute length in young of females.

**Fig. 8.**
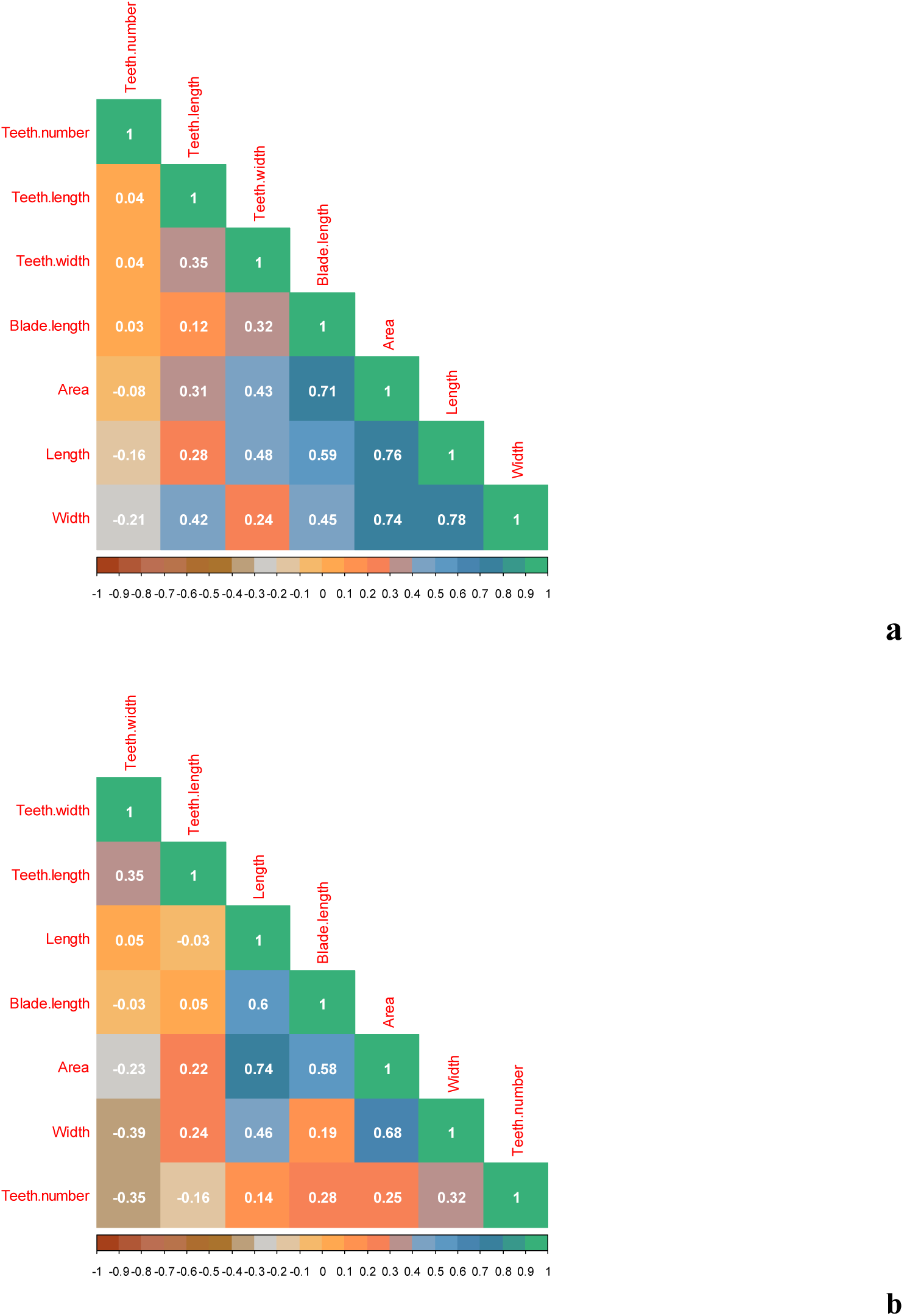
Correlation matrix show multicollinearity between morphological parameters of the first to fifth dorsal scutes of young males (a) and females (b) *Acipenser ruthenus* (fish length – 24.8±1.5 cm, 1 year-old). Number of fish treated = 20. Number of scutes treated = 100.

We found the correlation in morphological parameters and indices in juveniles sterlet (Fig. 9, 10). Strong correlation found between the scute area and the scute width, between the blade length of scute and the scute area in juveniles of males. Strong correlation found between the scute area and the scute width, between the blade length of scute and the scute width, between the scute length and the scute width, between the blade length of scute and the scute area, between the blade length of scute and the scute length, between the scute area and the scute length in juveniles of females.

**Fig. 9.**
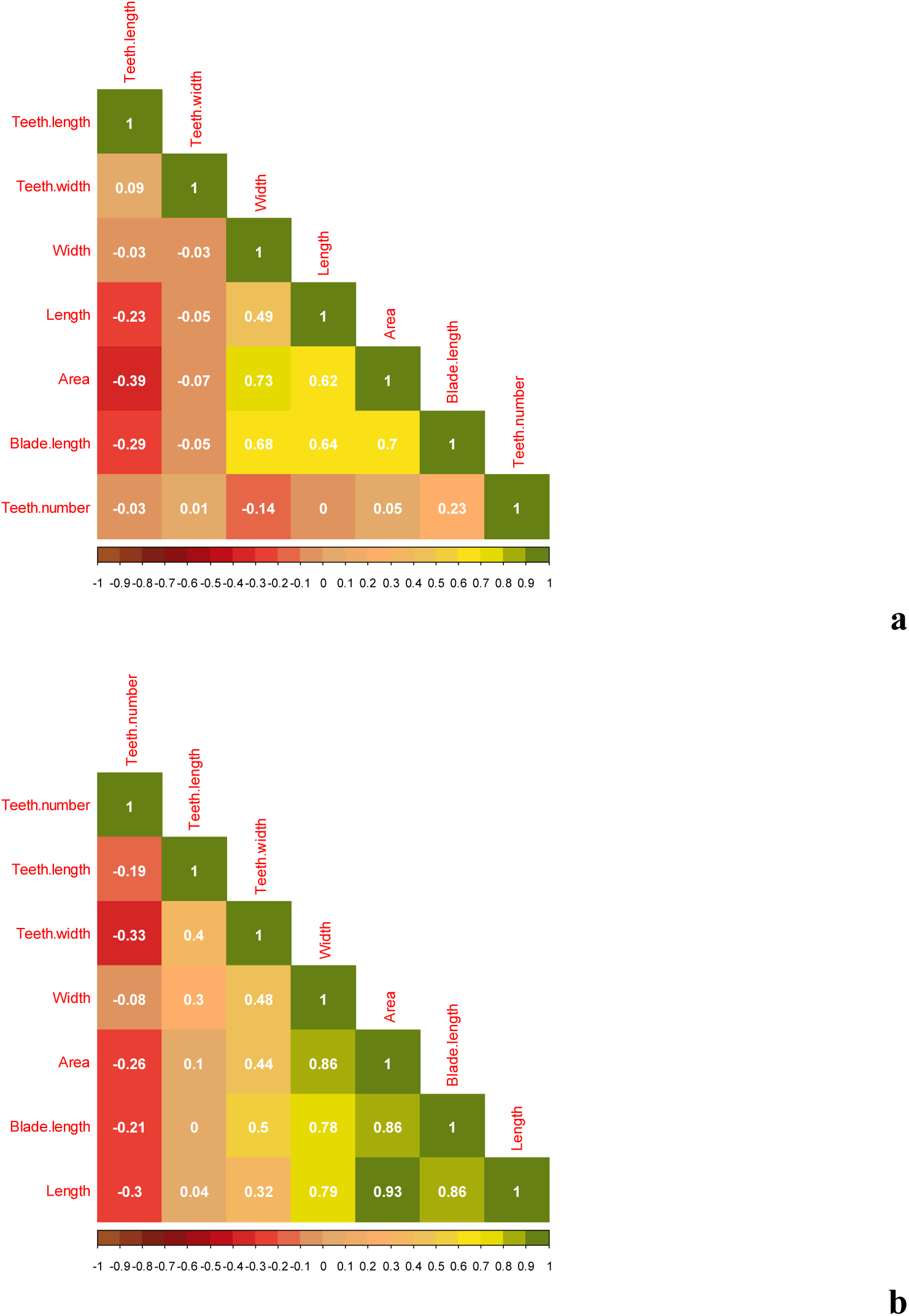
Correlation matrix show multicollinearity between morphological parameters of the first to fifth dorsal scutes of juveniles males (a) and females (b) *Acipenser ruthenus* (fish length – 70.3±3.6 mm, 3 months-old). Number of fish treated = 20. Number of scutes treated = 100.

**Fig. 10.**
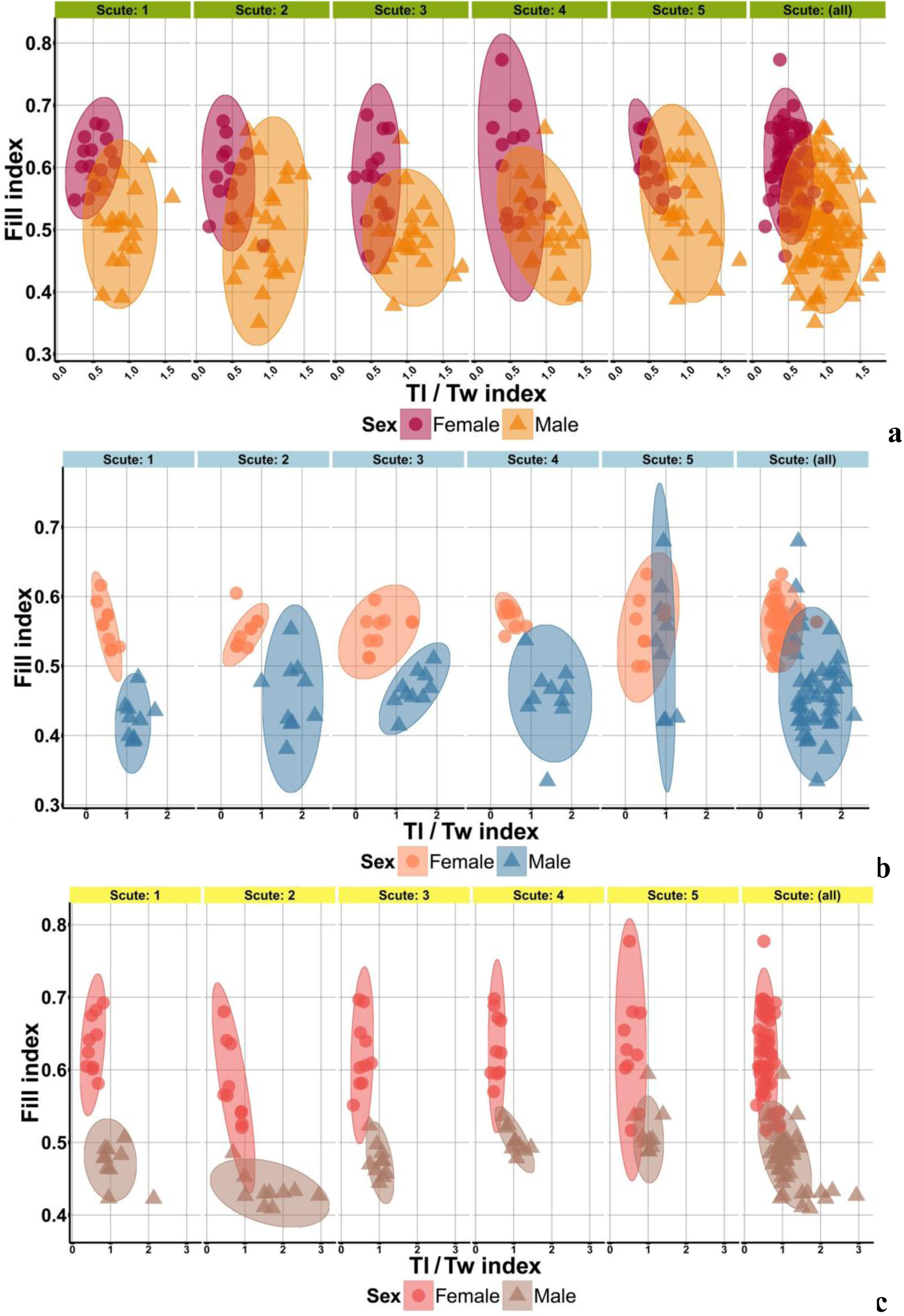
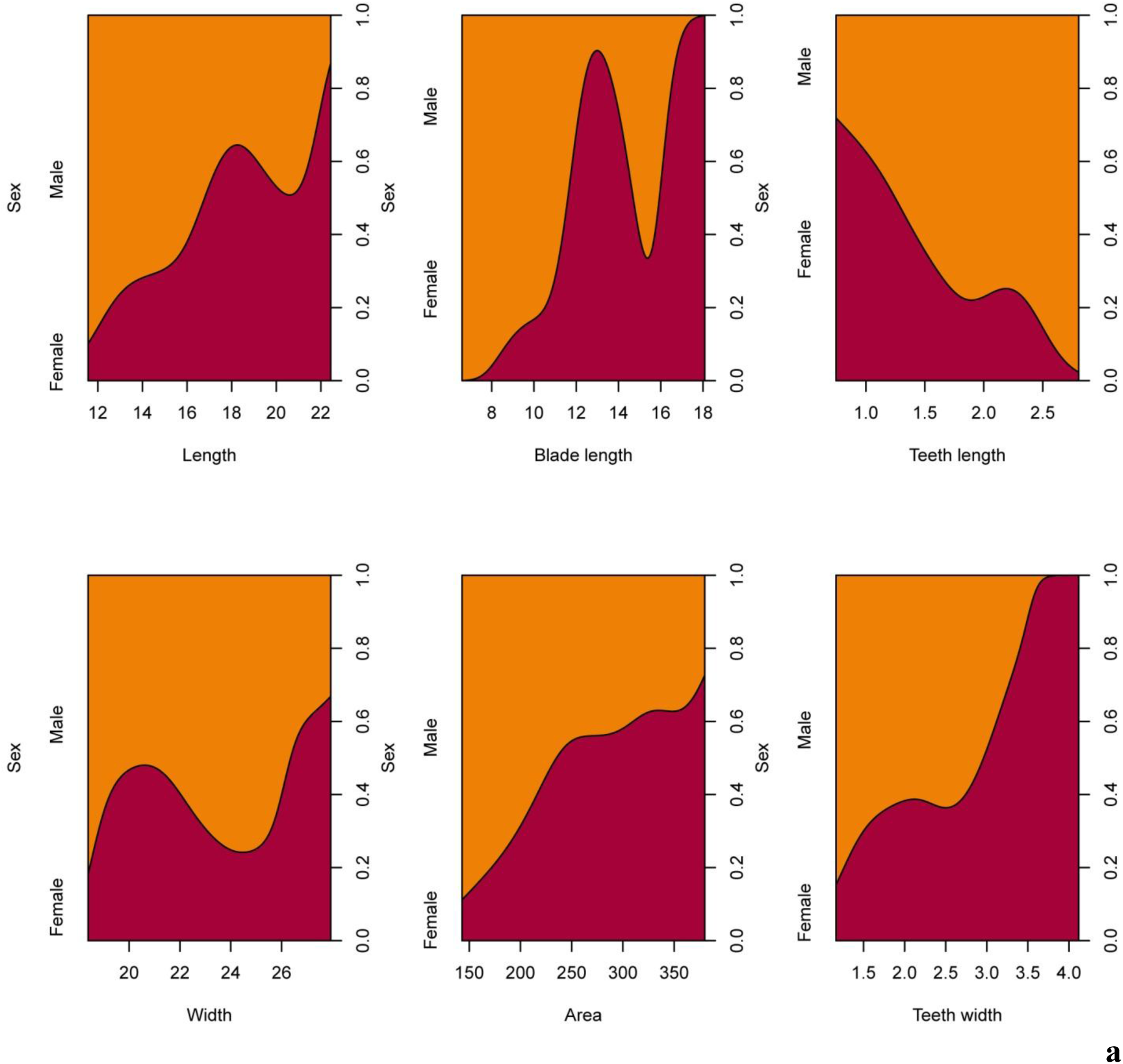
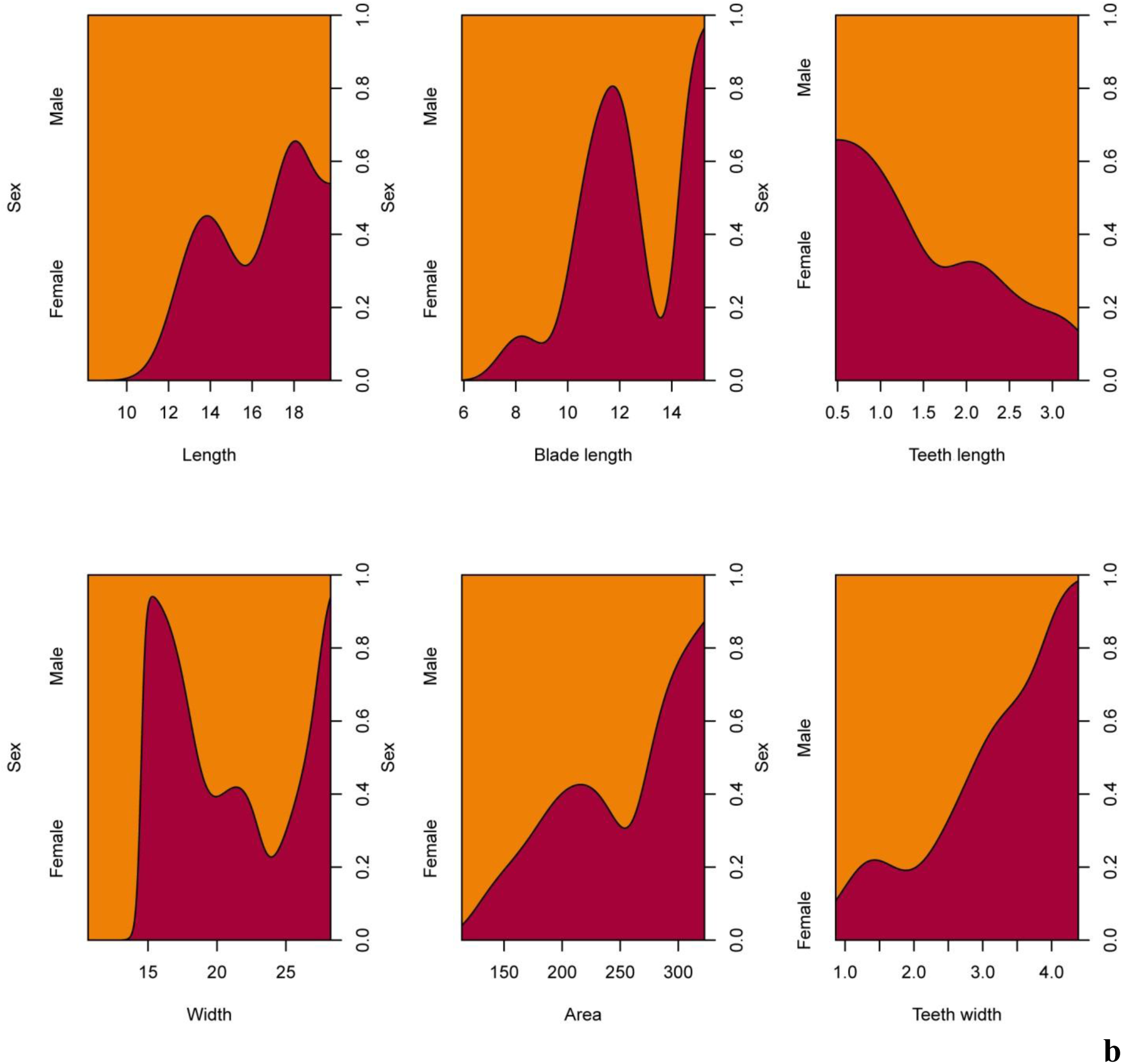
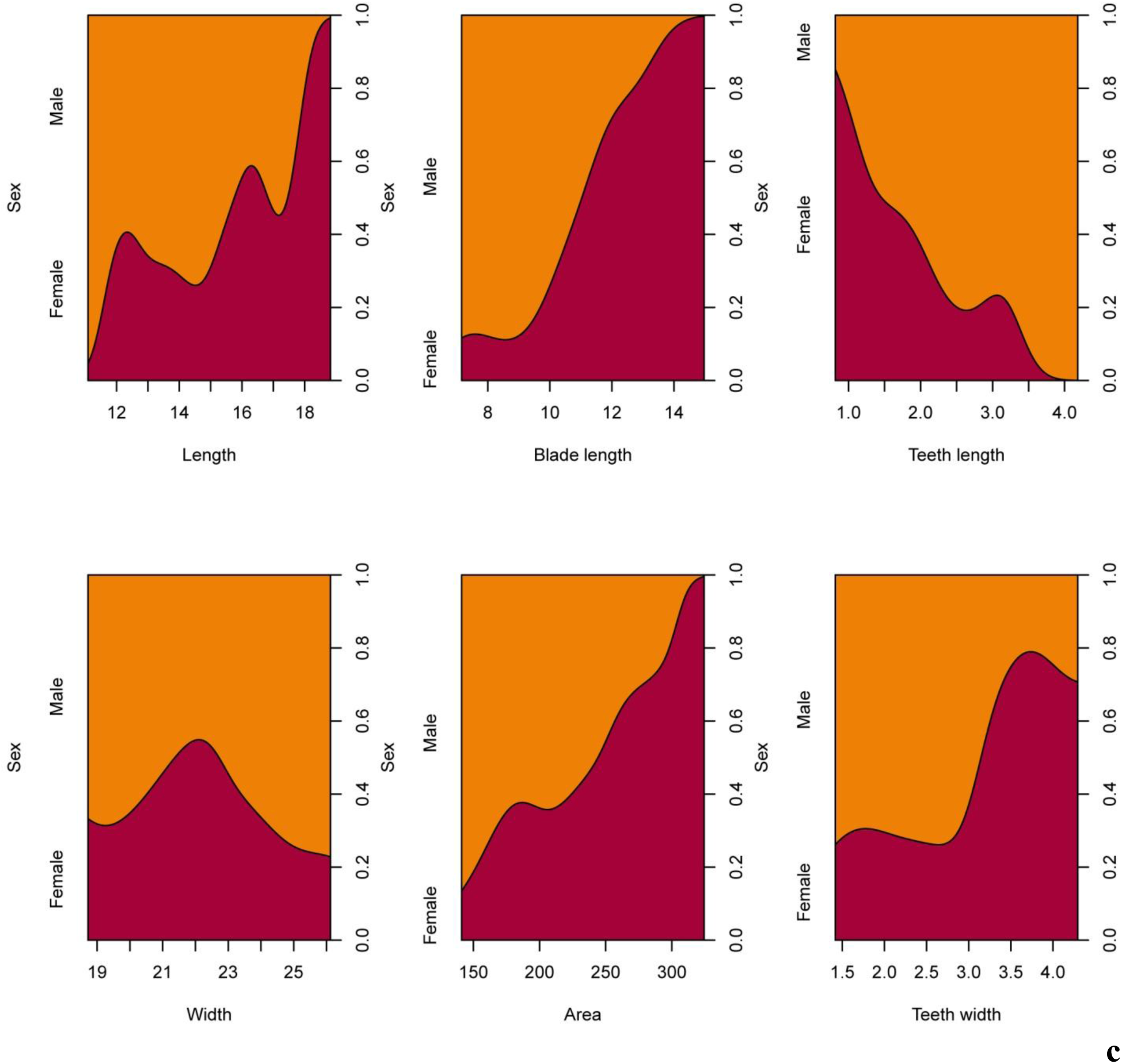
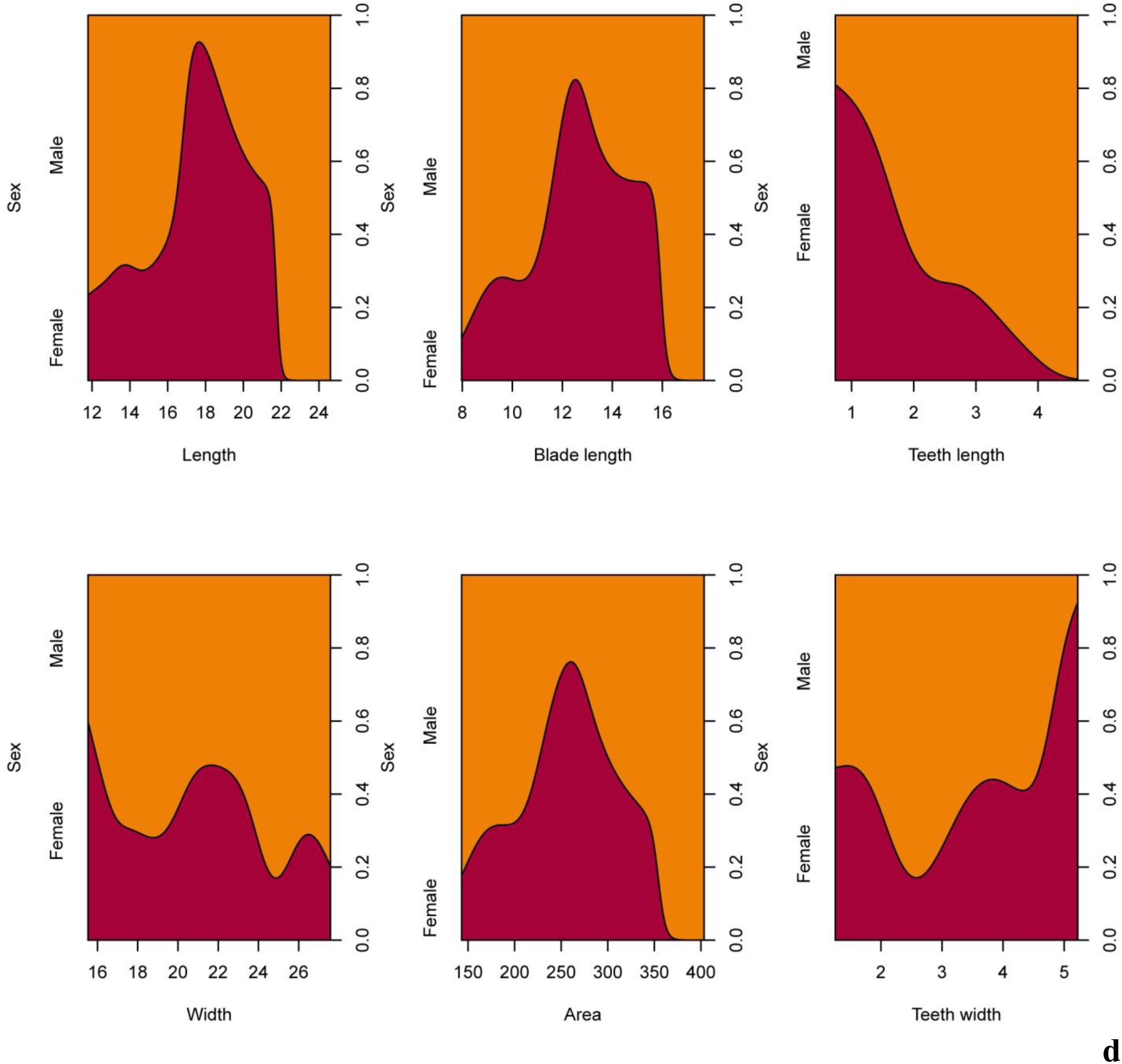
Results of distribution points show a sex-class separation between important morphological indices for adults (a), young (b) and juveniles (c) depending on scute number. Number of fish treated = 37 (adults), 20 (young), 20 (juveniles). Number of scutes treated = 370 (adults), 200 (young), 200 (juveniles).

We found relationships in morphological parameters and indices between males and females of adult sterlet (Fig. 11, 12). Increase (the scute area, the teeth width, the blade length, the fill index, the Bl/L index) and decrease (the teeth length, the W/L index, the Tl/Tw index, the Tl/W index, the teeth number) of the morphological parameter and index size of the first five dorsal scute increase the sex identification probability.

**Fig. 11.**
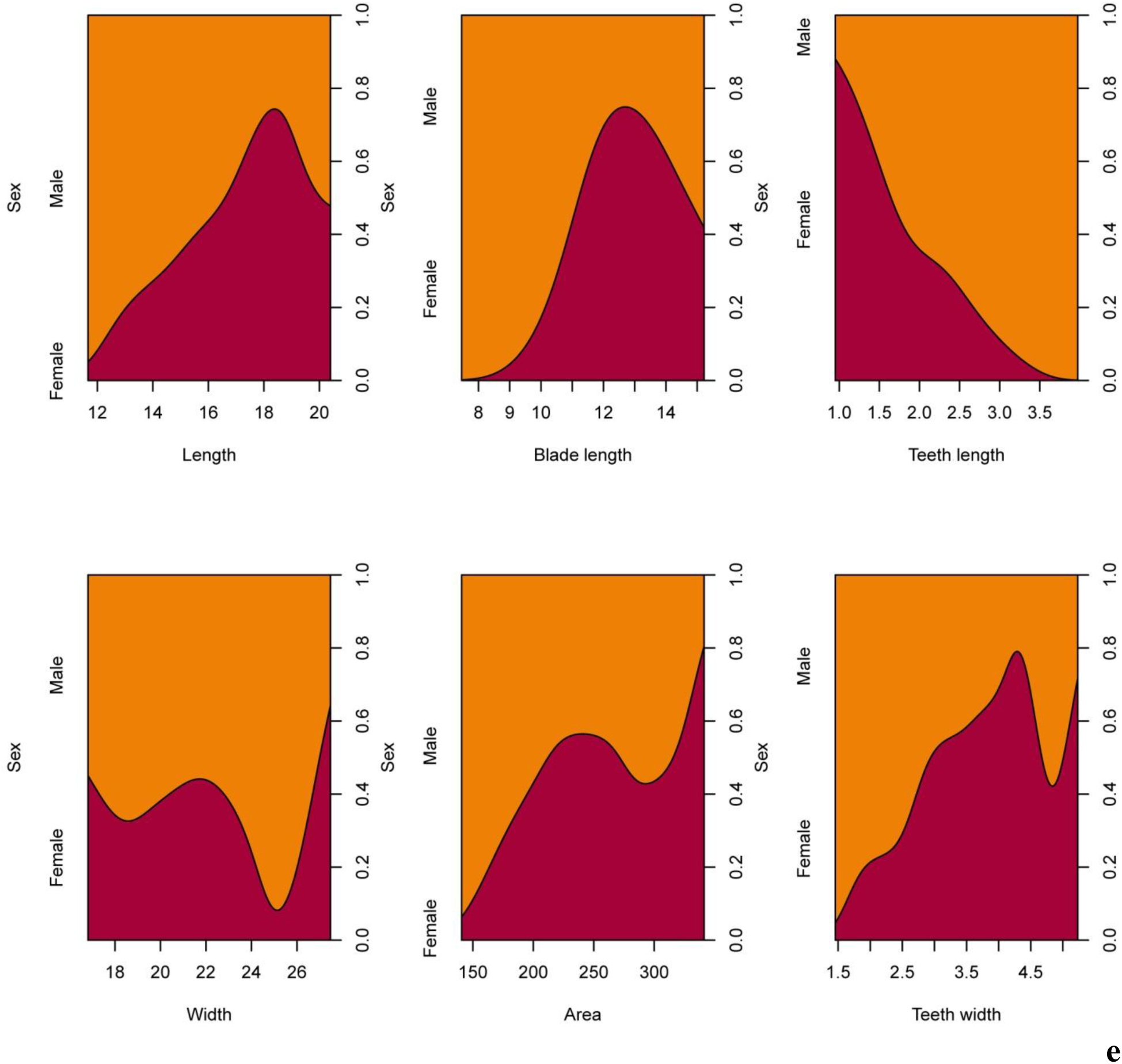
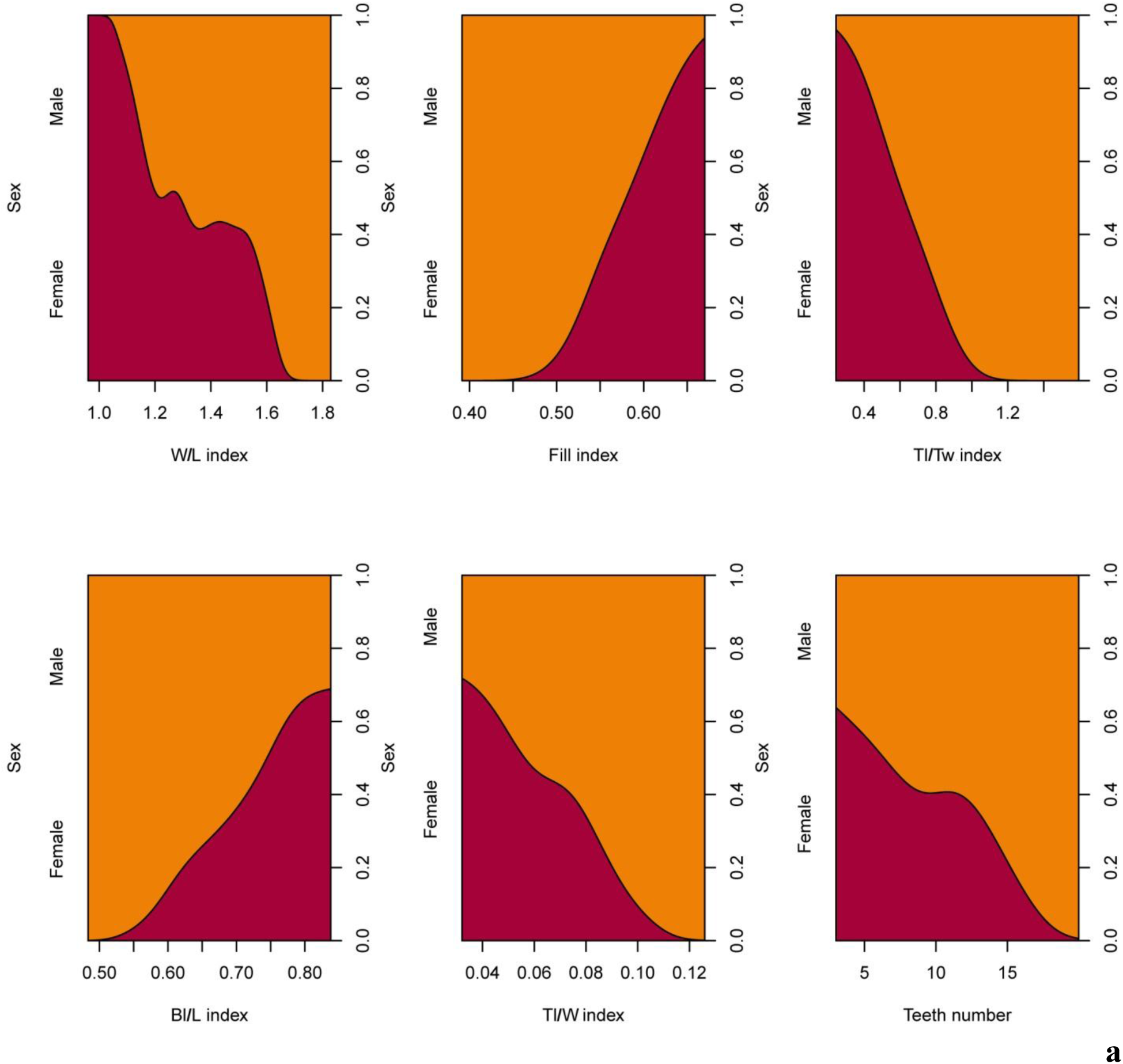
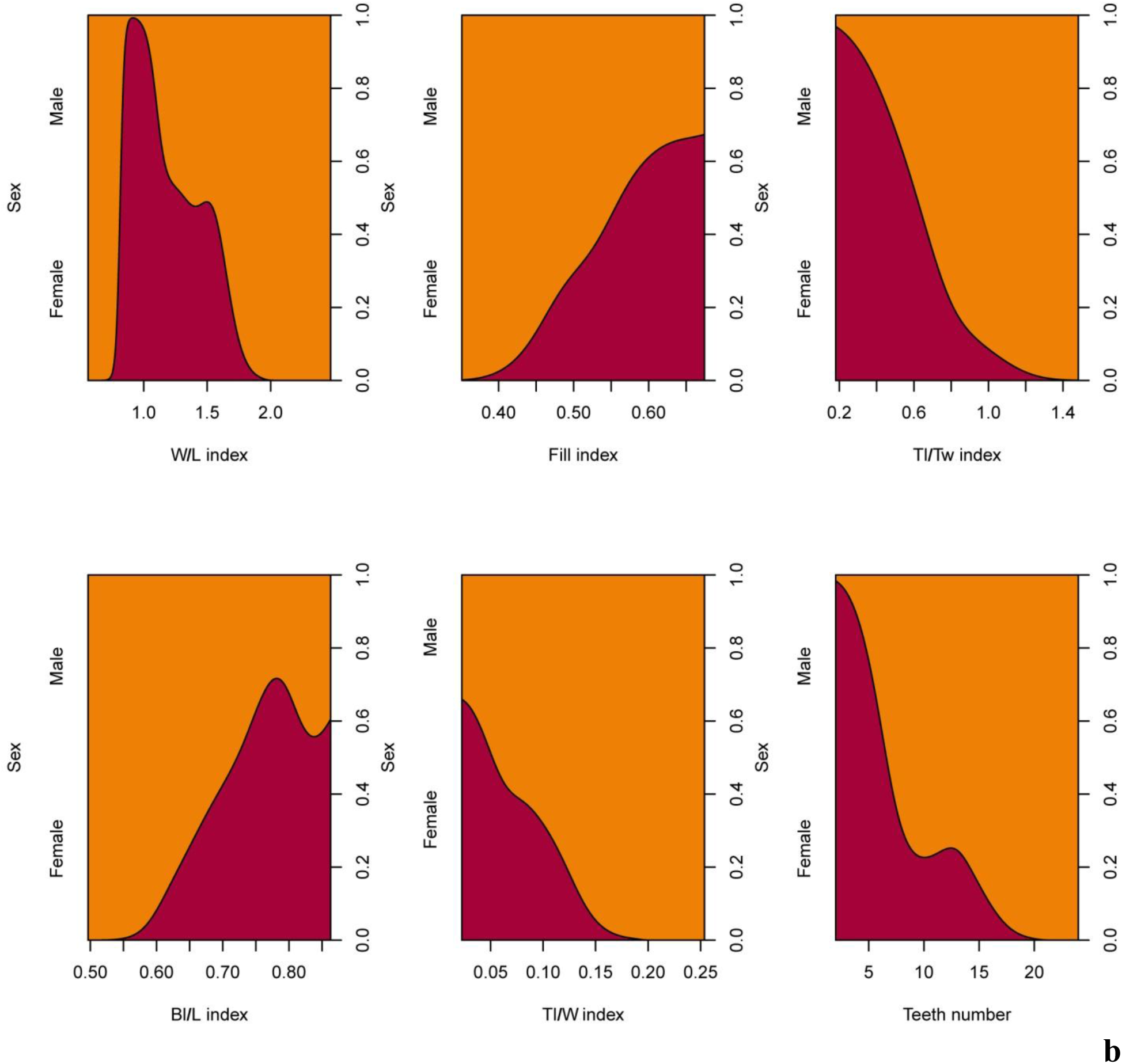
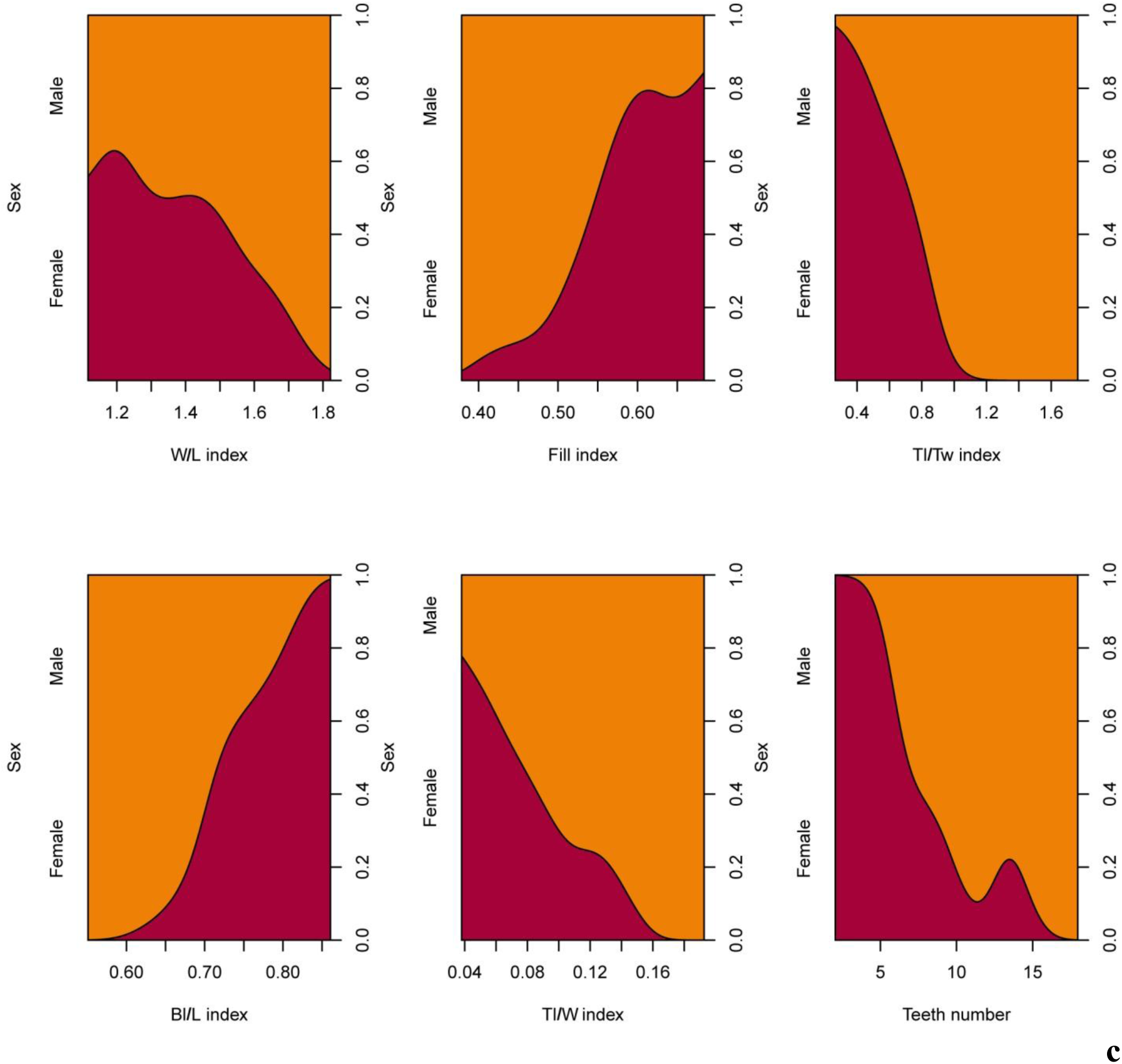
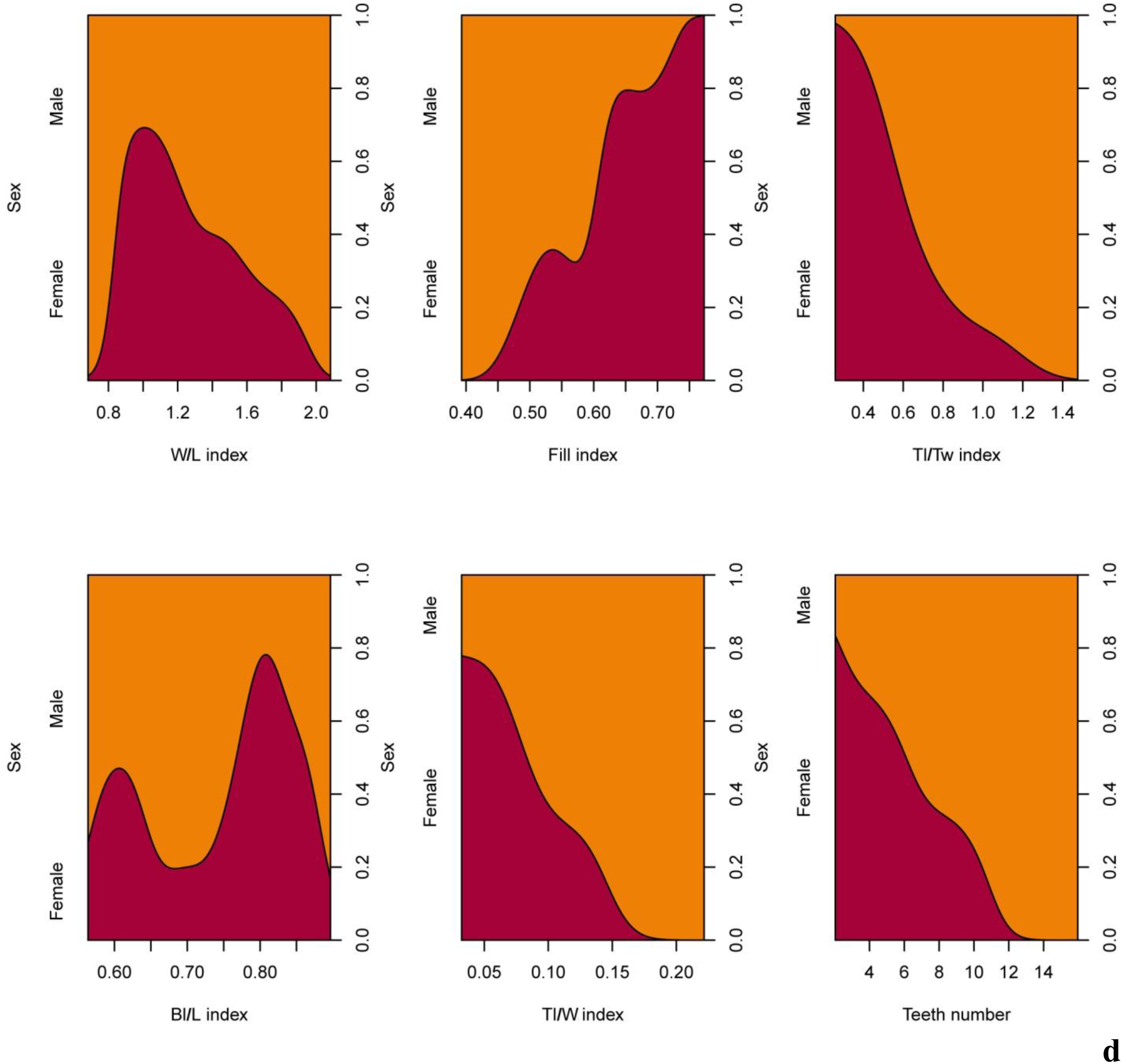
Conditional density plots of morphological parameters of the first (a), second (b), third (c), fourth (d), fifth (e) dorsal scutes of adults *Acipenser ruthenus*. Number of fish treated = 37. Number of scutes treated = 185.

**Fig. 12.**
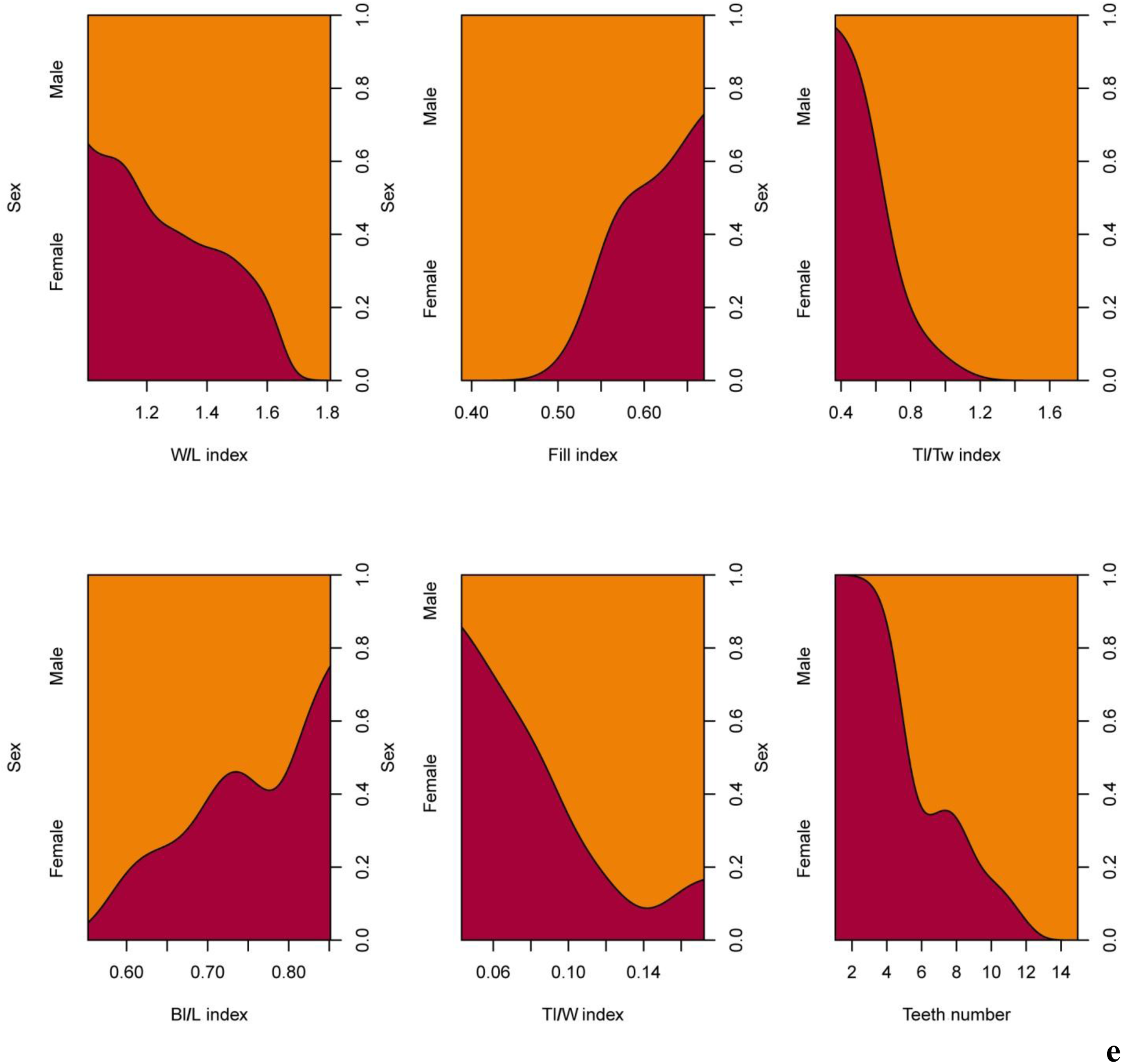
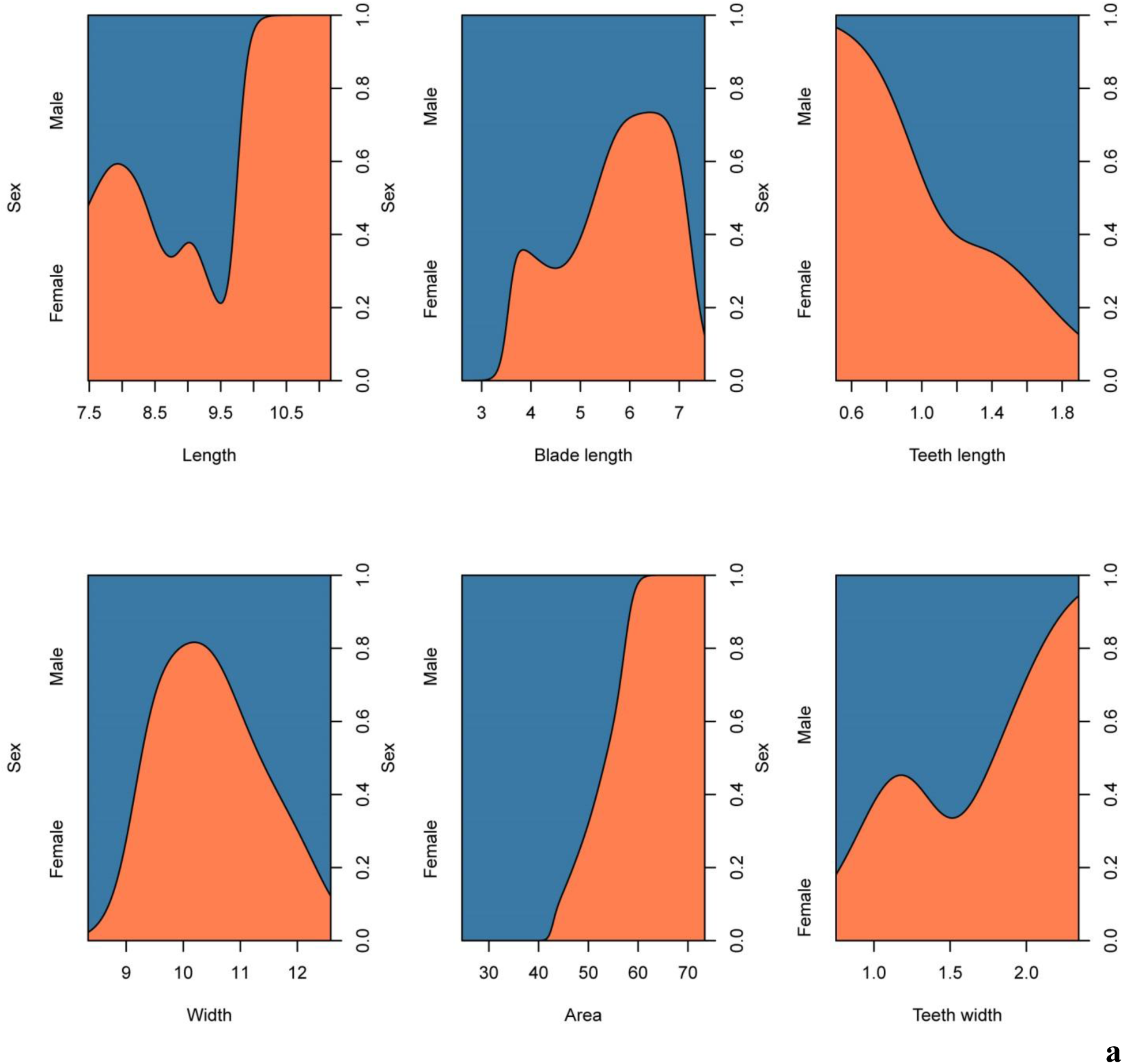
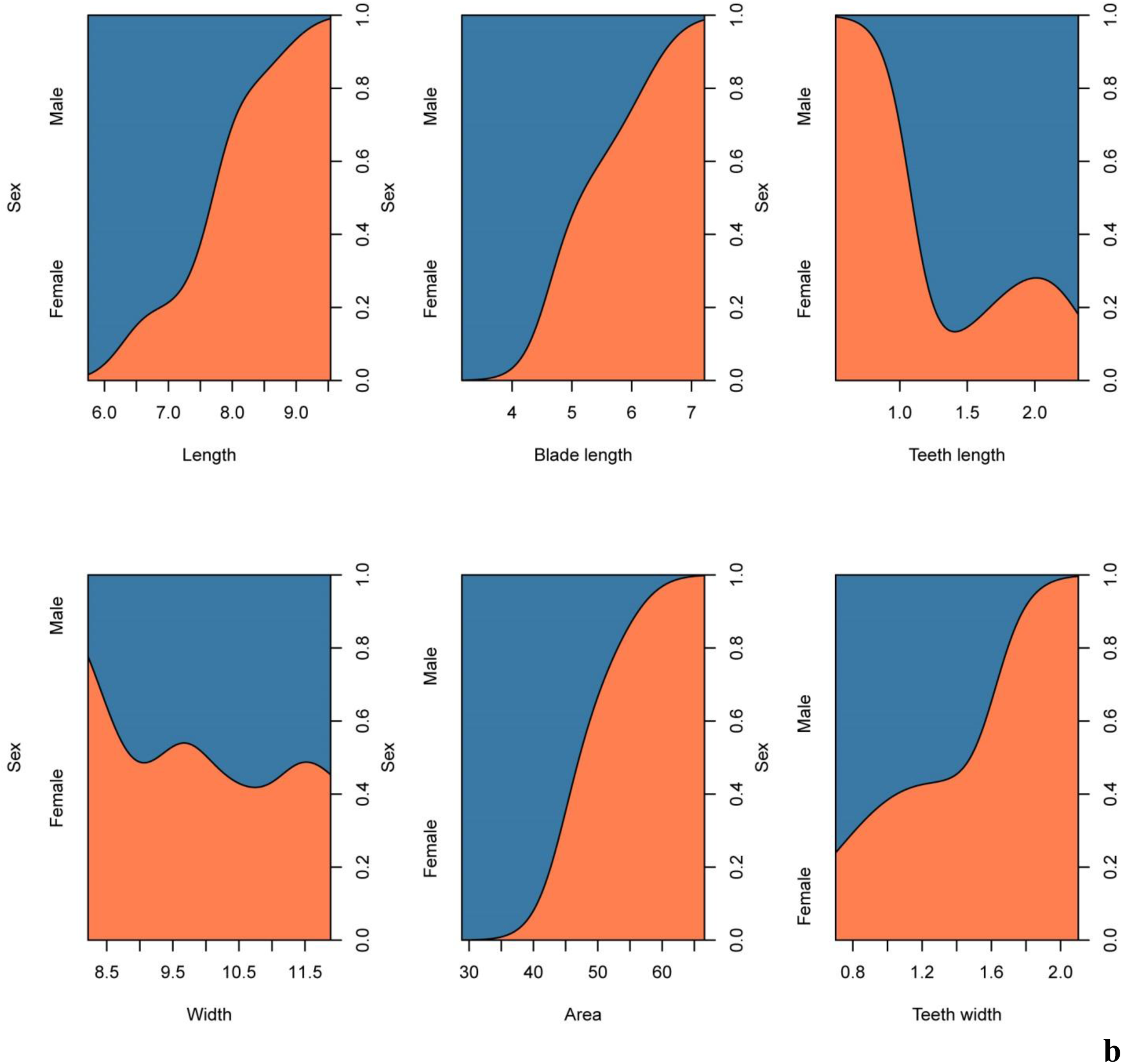
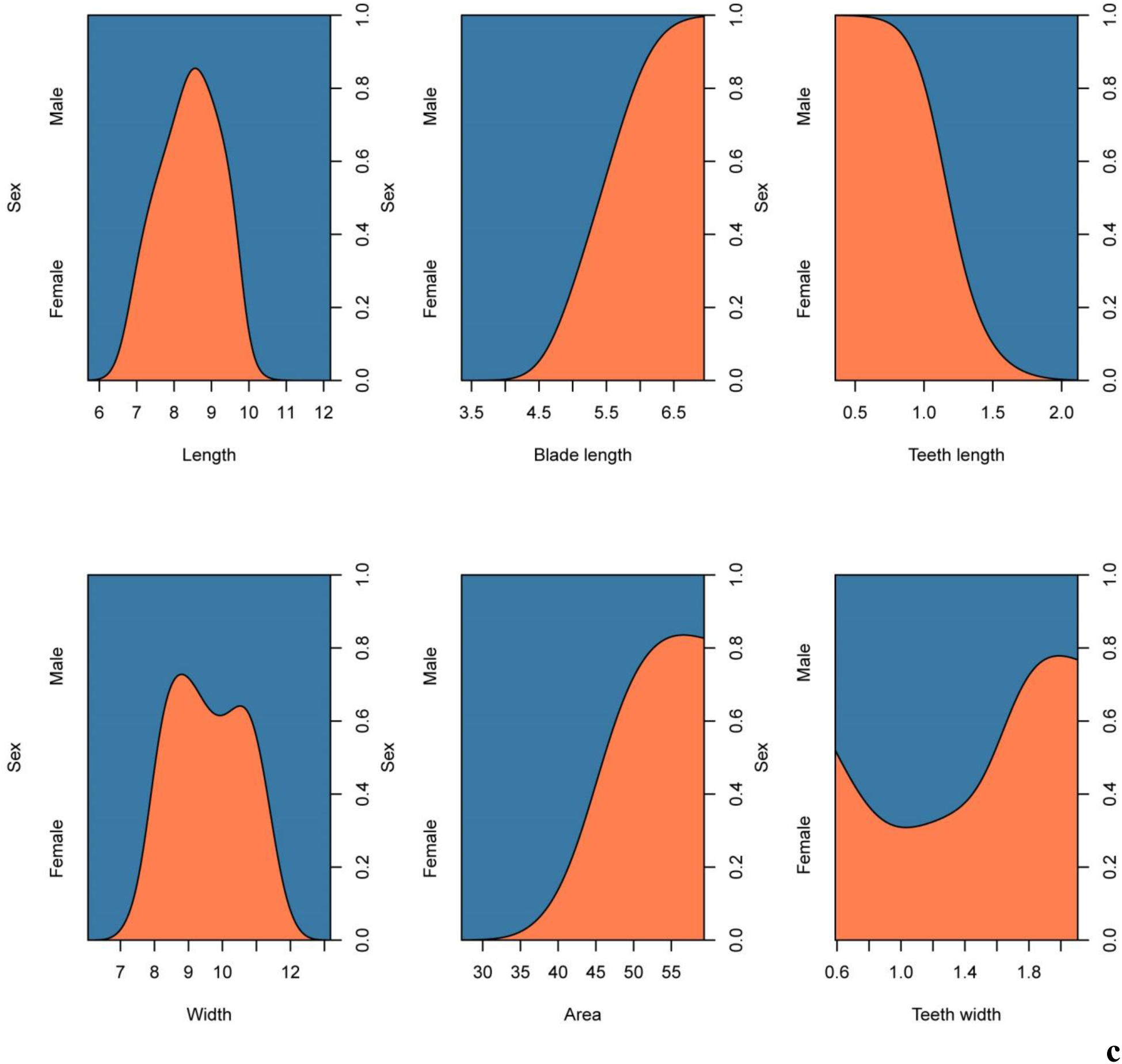
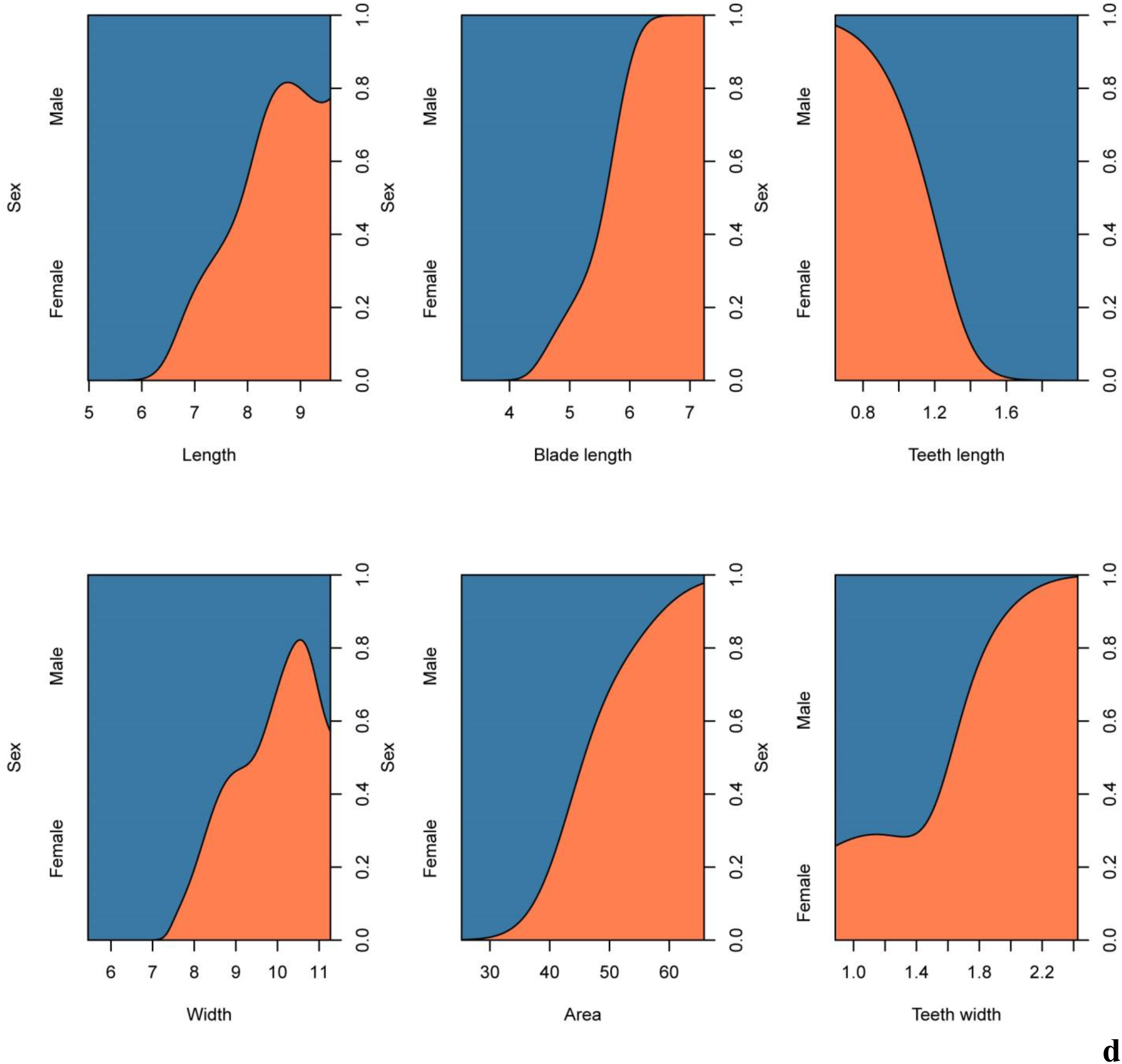
Conditional density plots of morphological indices of the first (a), second (b), third (c), fourth (d), fifth (e) dorsal scutes of adults *Acipenser ruthenus*. Number of fish treated = 37. Number of scutes treated = 185.

We found relationships in morphological parameters and indices between males and females of young sterlet (Fig. 13, 14). Increase (the scute area, the teeth width, the blade length, the scute length, the fill index, the Bl/L index) and decrease (the teeth length, the Tl/Tw index, the Tl/W index, the teeth number, the W/L index, the teeth number) of the morphological parameter and index size of the first five dorsal scute increase the sex identification probability.

**Fig. 13.**
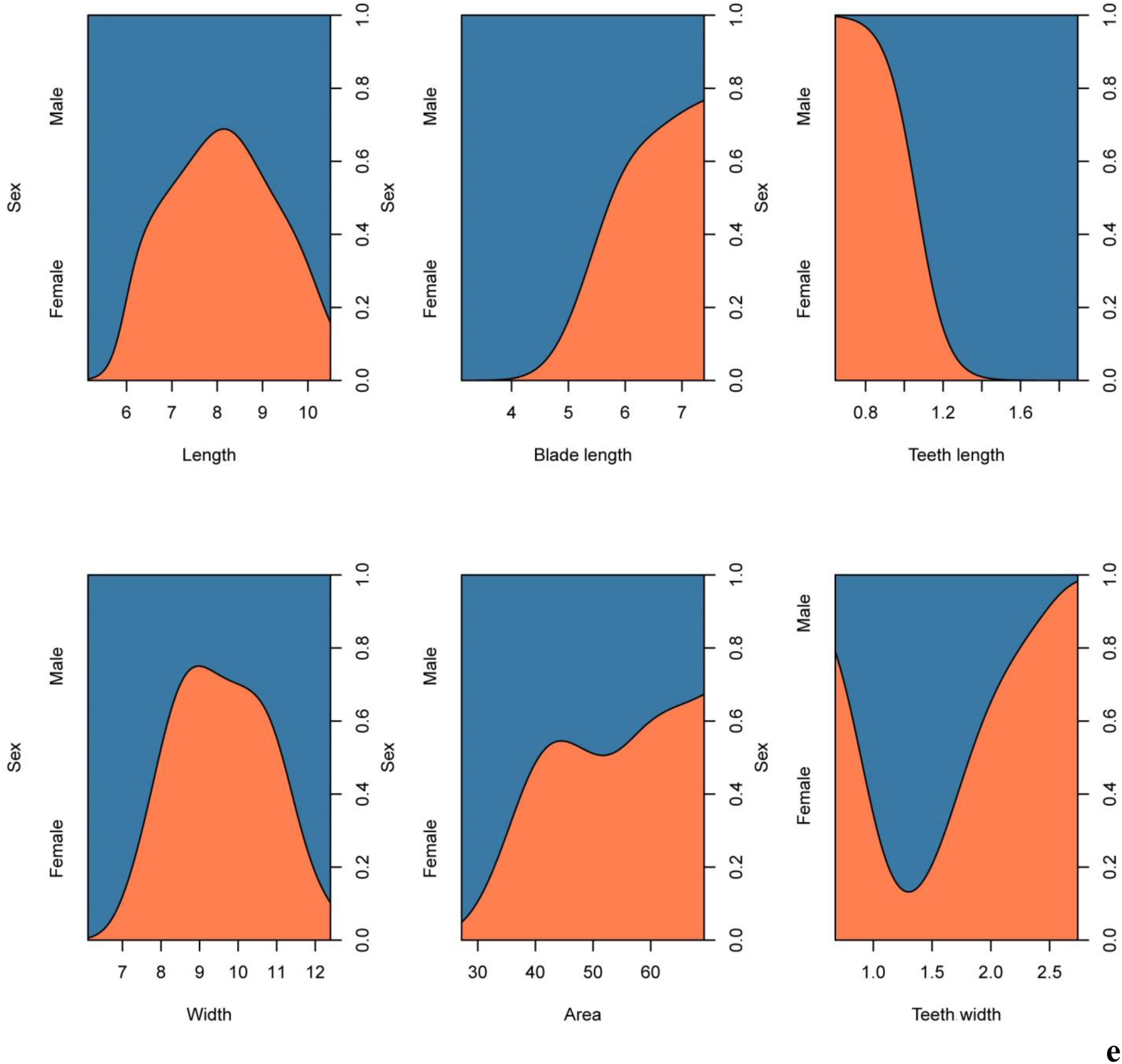
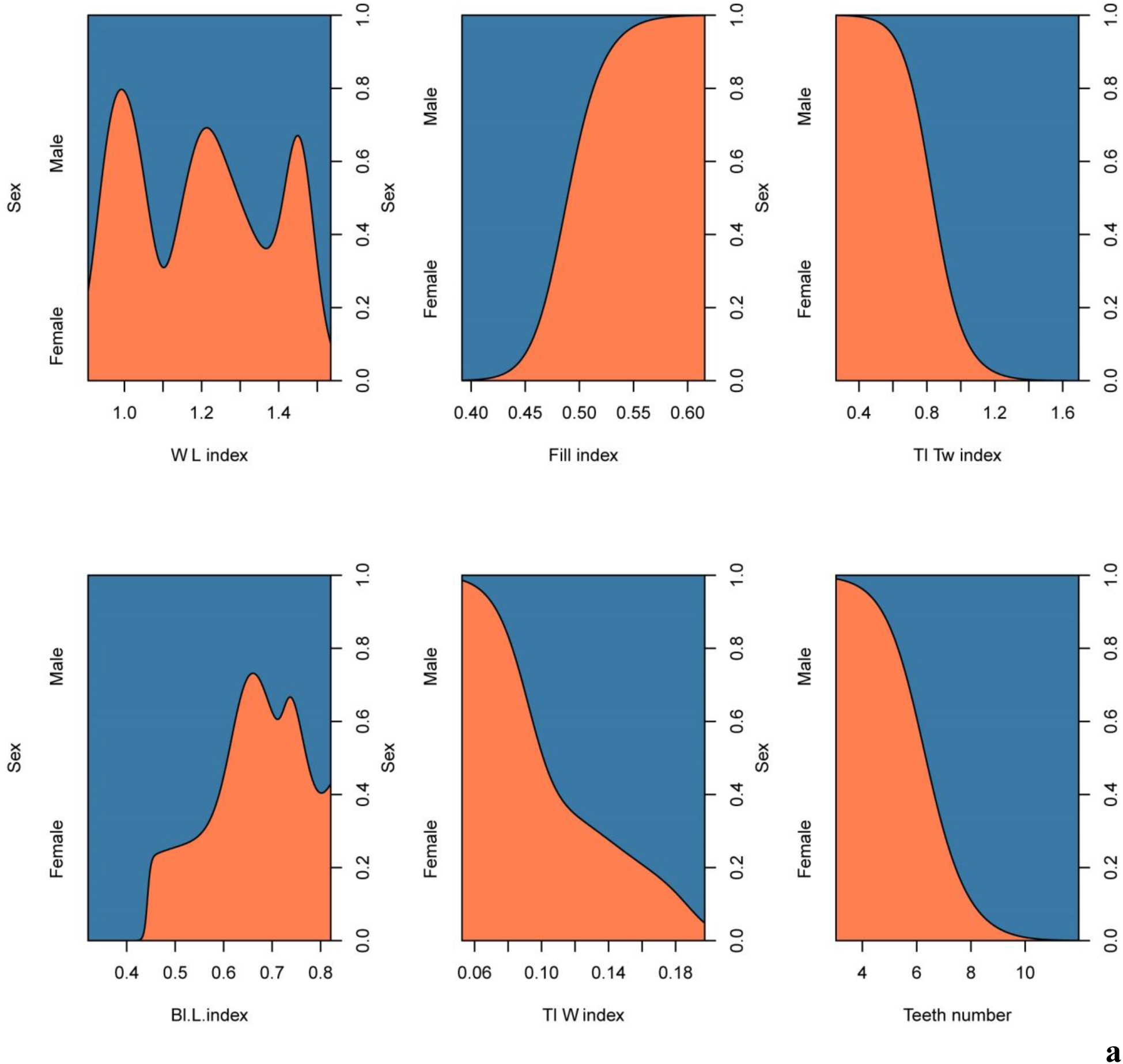
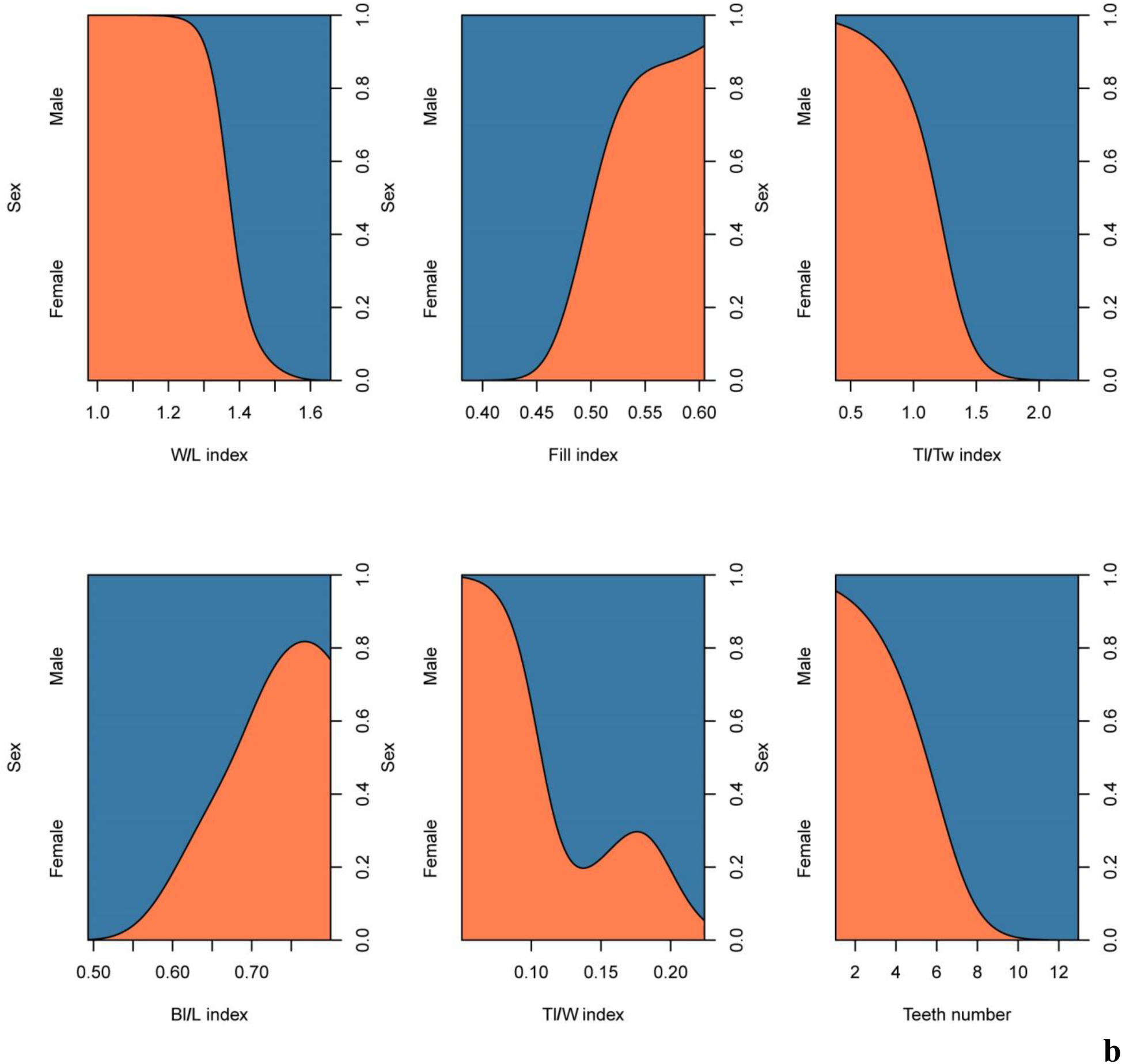
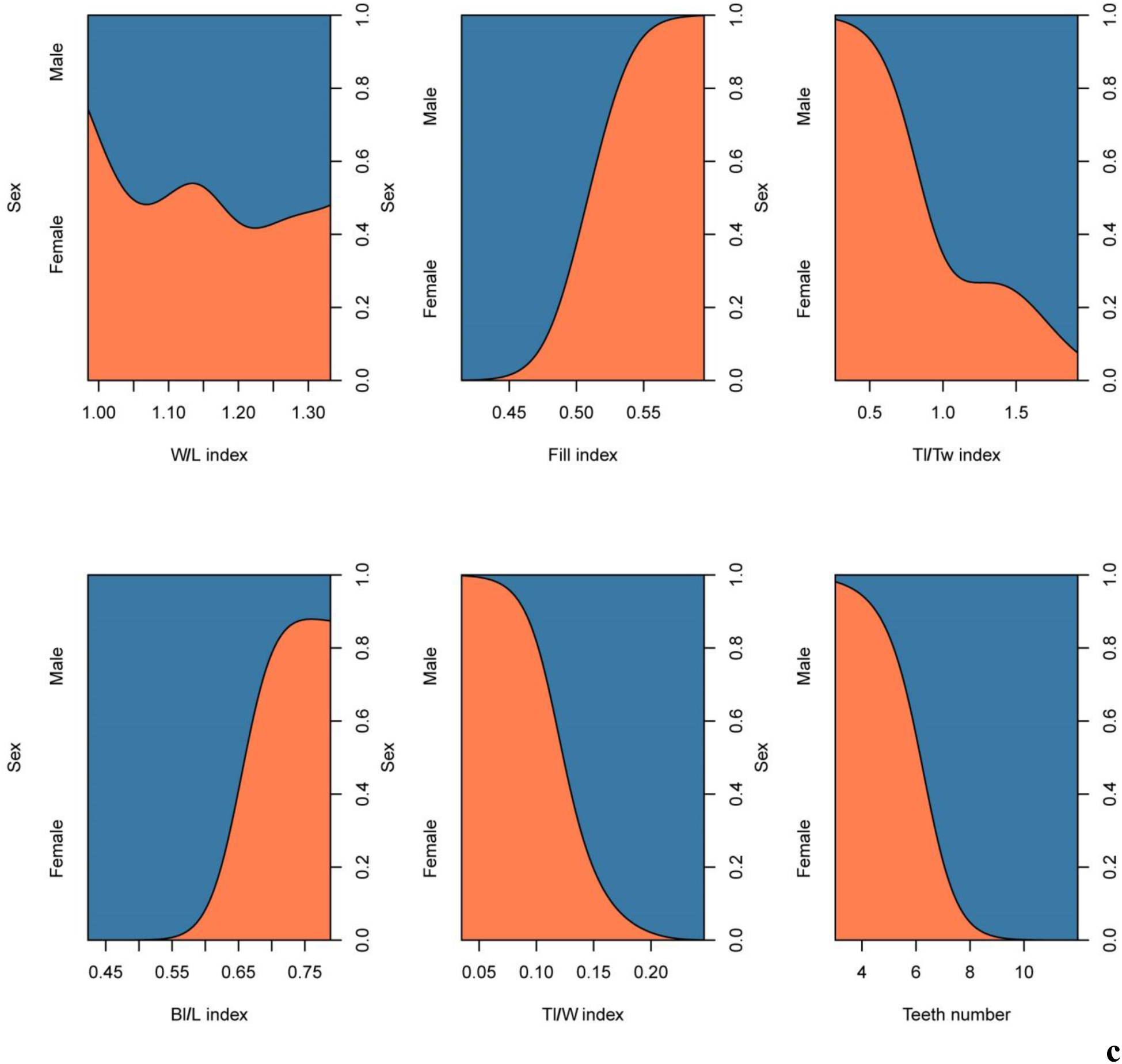
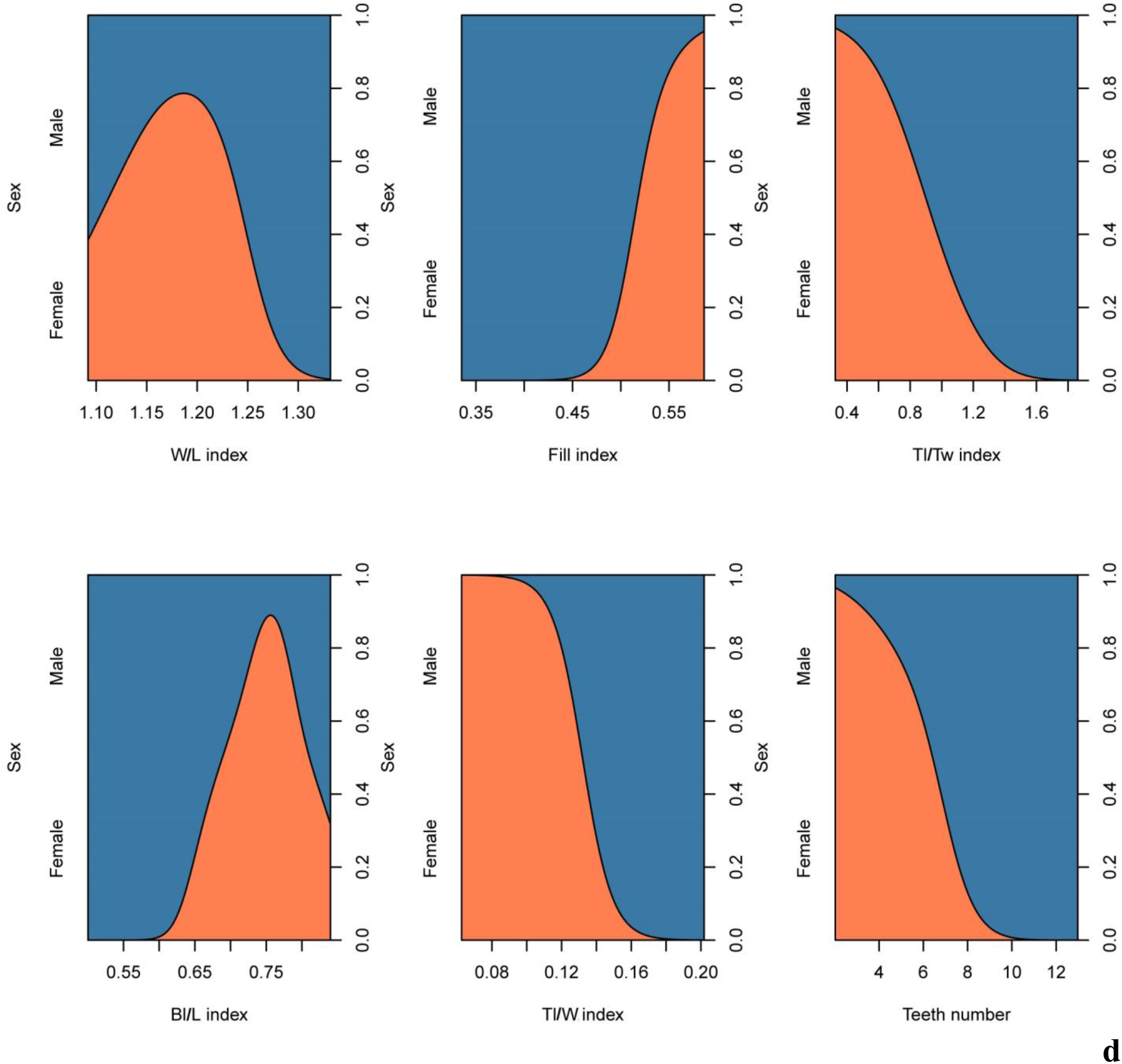
Conditional density plots of morphological parameters of the first (a), second (b), third (c), fourth (d), fifth (e) dorsal scutes of young *Acipenser ruthenus*. Number of fish treated = 20. Number of scutes treated = 100.

**Fig. 14.**
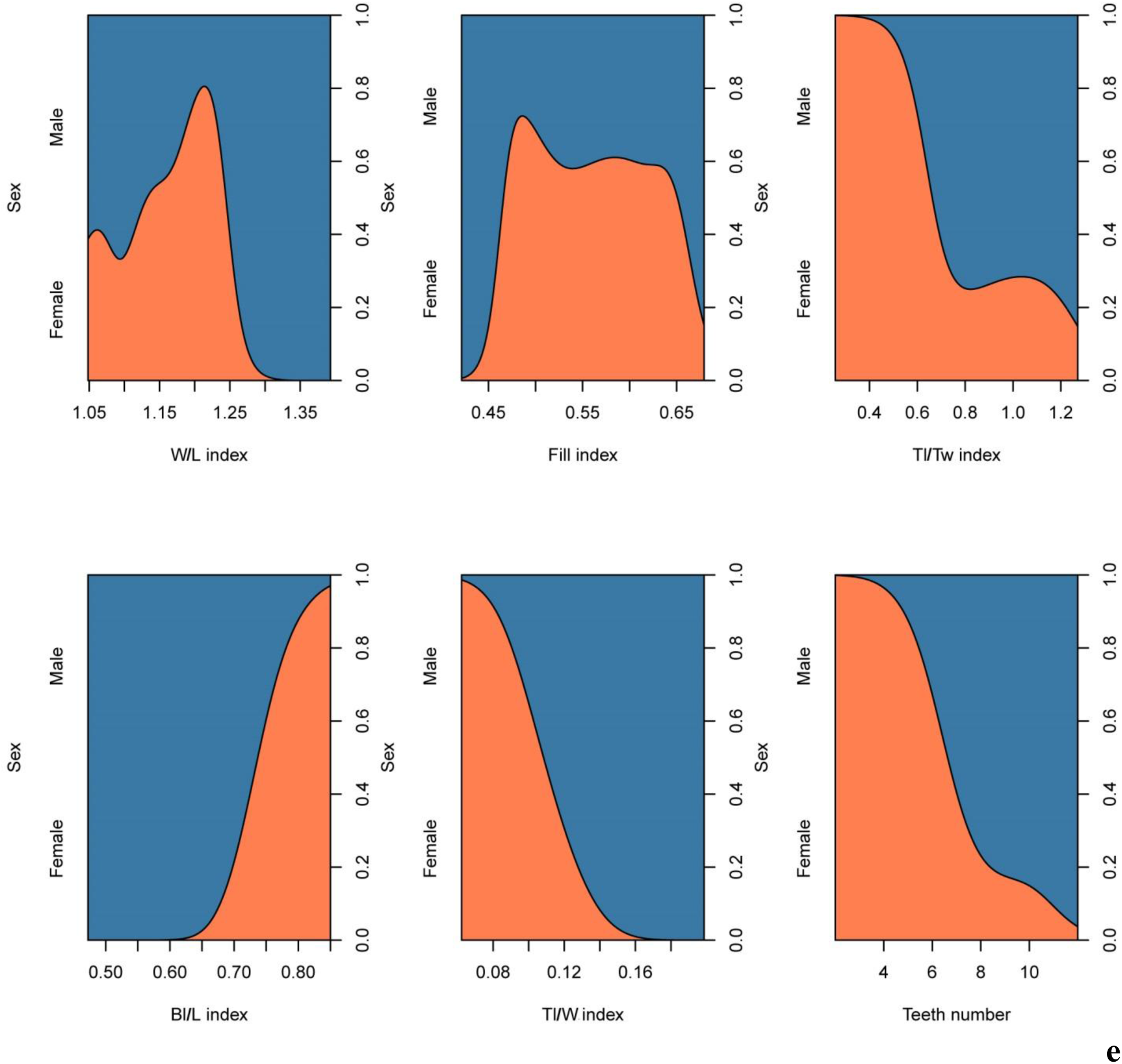
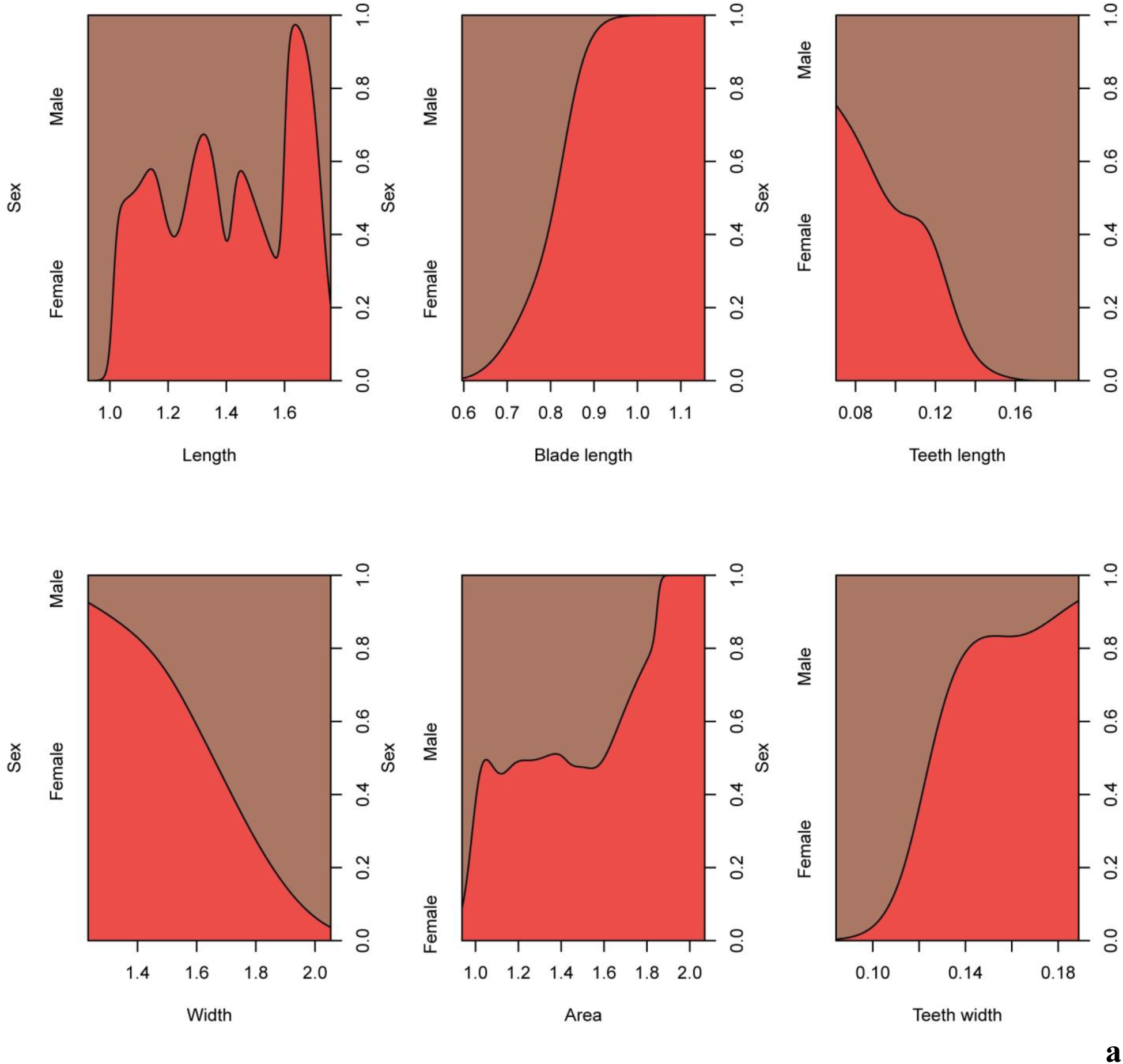
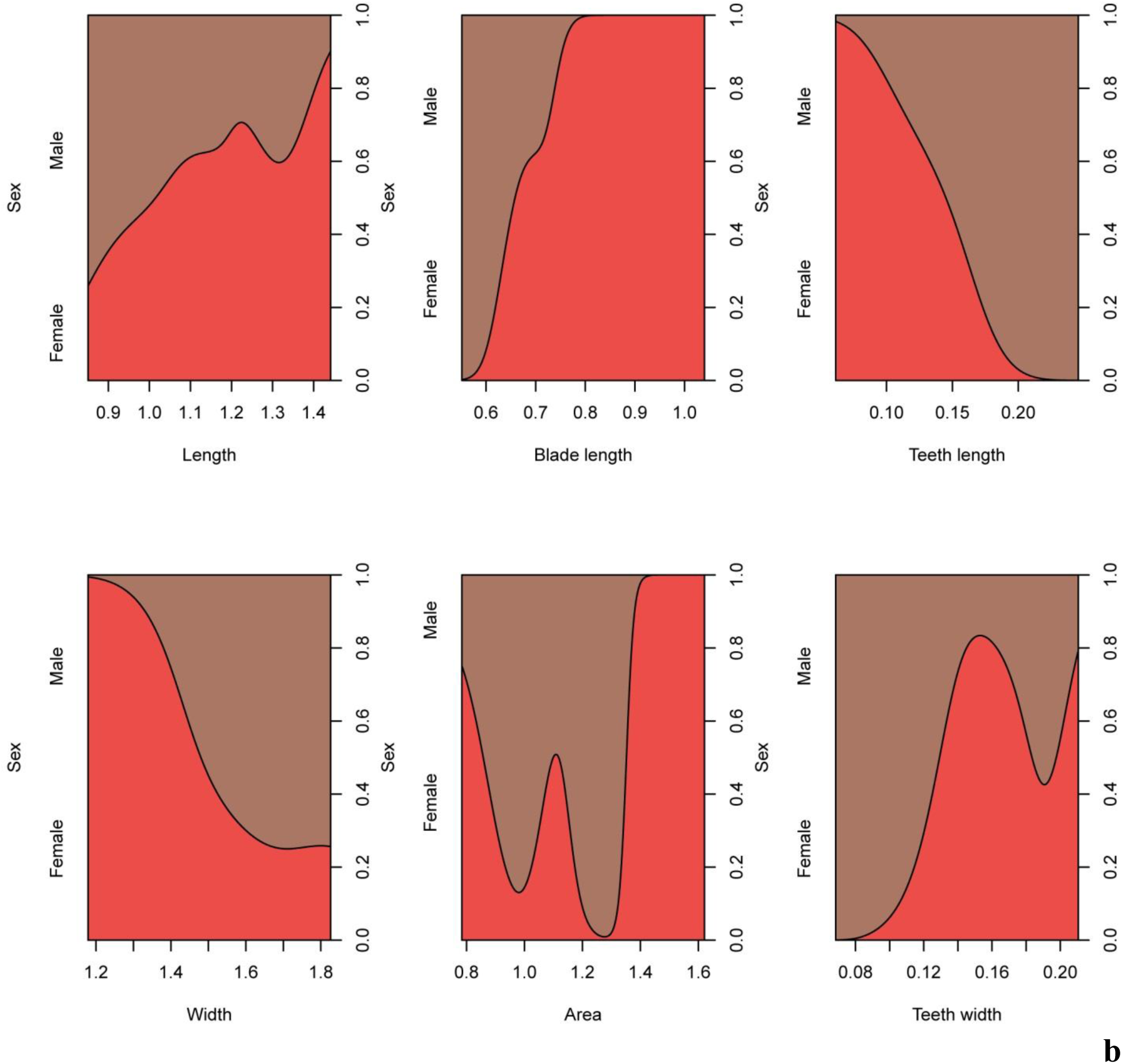
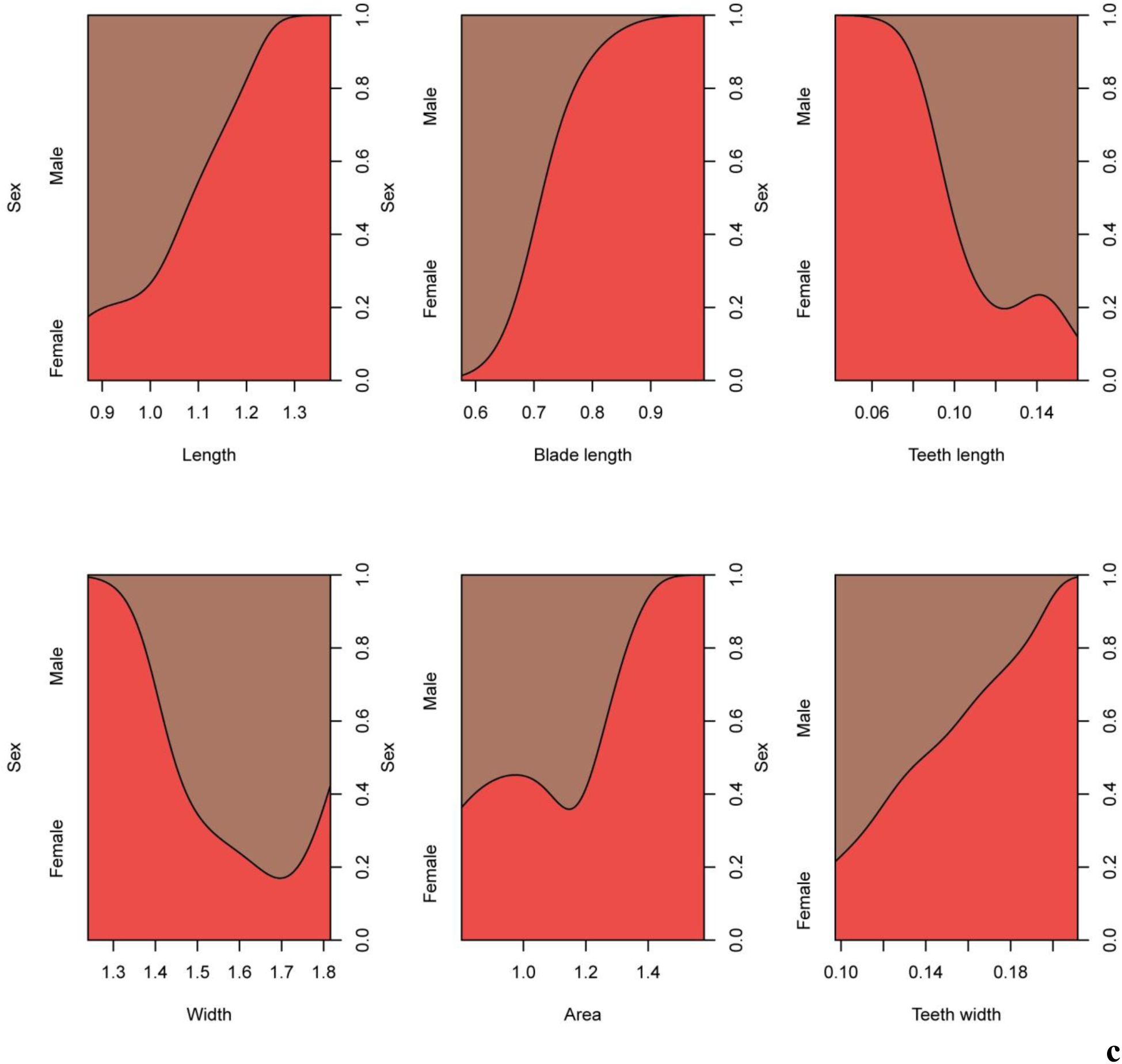
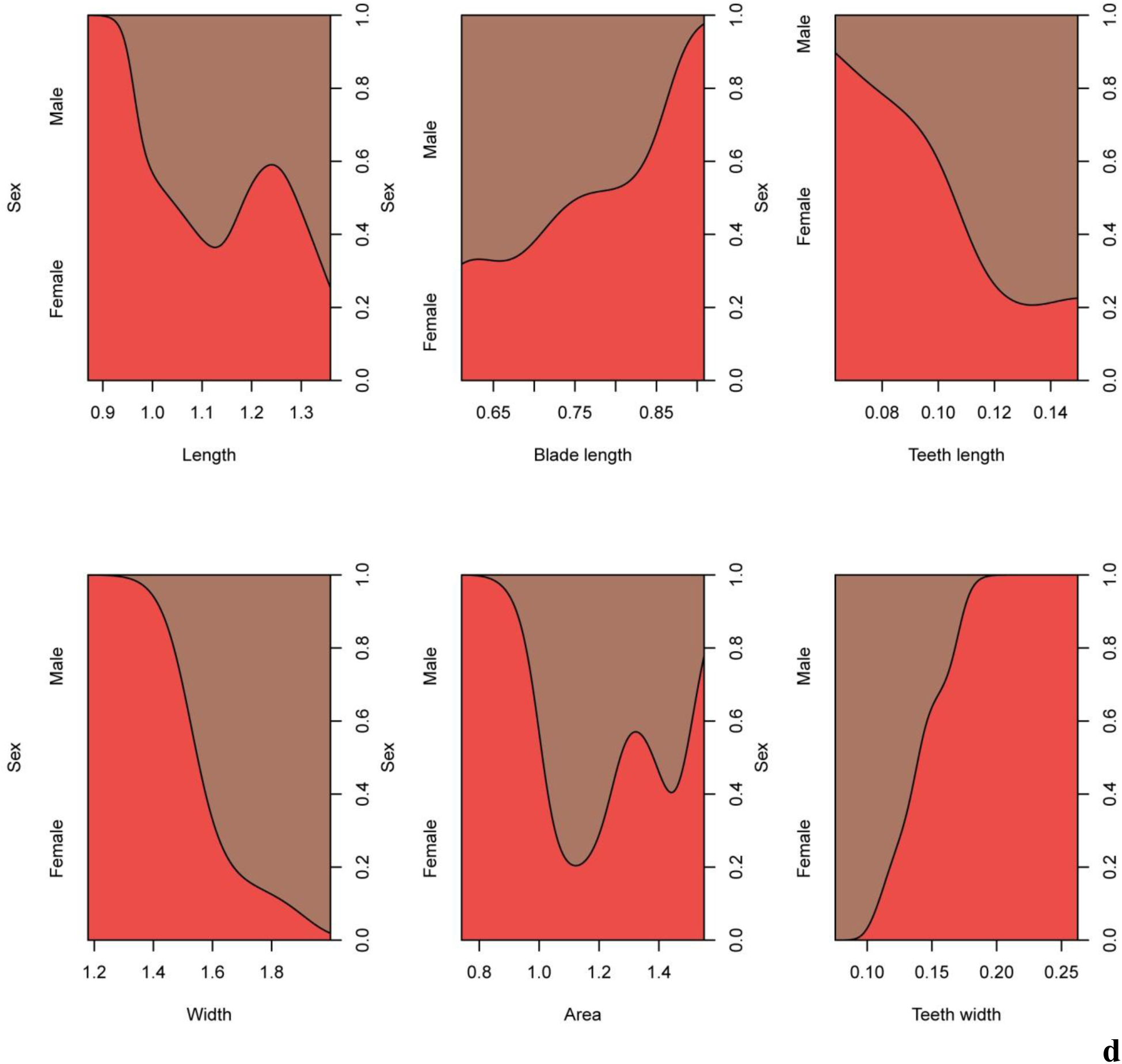
Conditional density plots of morphological indices of the first (a), second (b), third (c), fourth (d), fifth (e) dorsal scutes of young *Acipenser ruthenus*. Number of fish treated = 20. Number of scutes treated = 100.

We found relationships in morphological parameters and indices between males and females of juveniles sterlet (Fig. 15, 16). Increase (the blade length, the scute area, the teeth width, the scute length, the fill index, the Bl/L index) and decrease (the teeth length, the scute with, the Tl/Tw index, the teeth number, the W/L index, the Tl/Tw index, the Tl/W index) of the morphological parameter and index size of the first five dorsal scute increase the sex identification probability.

**Fig. 15.**
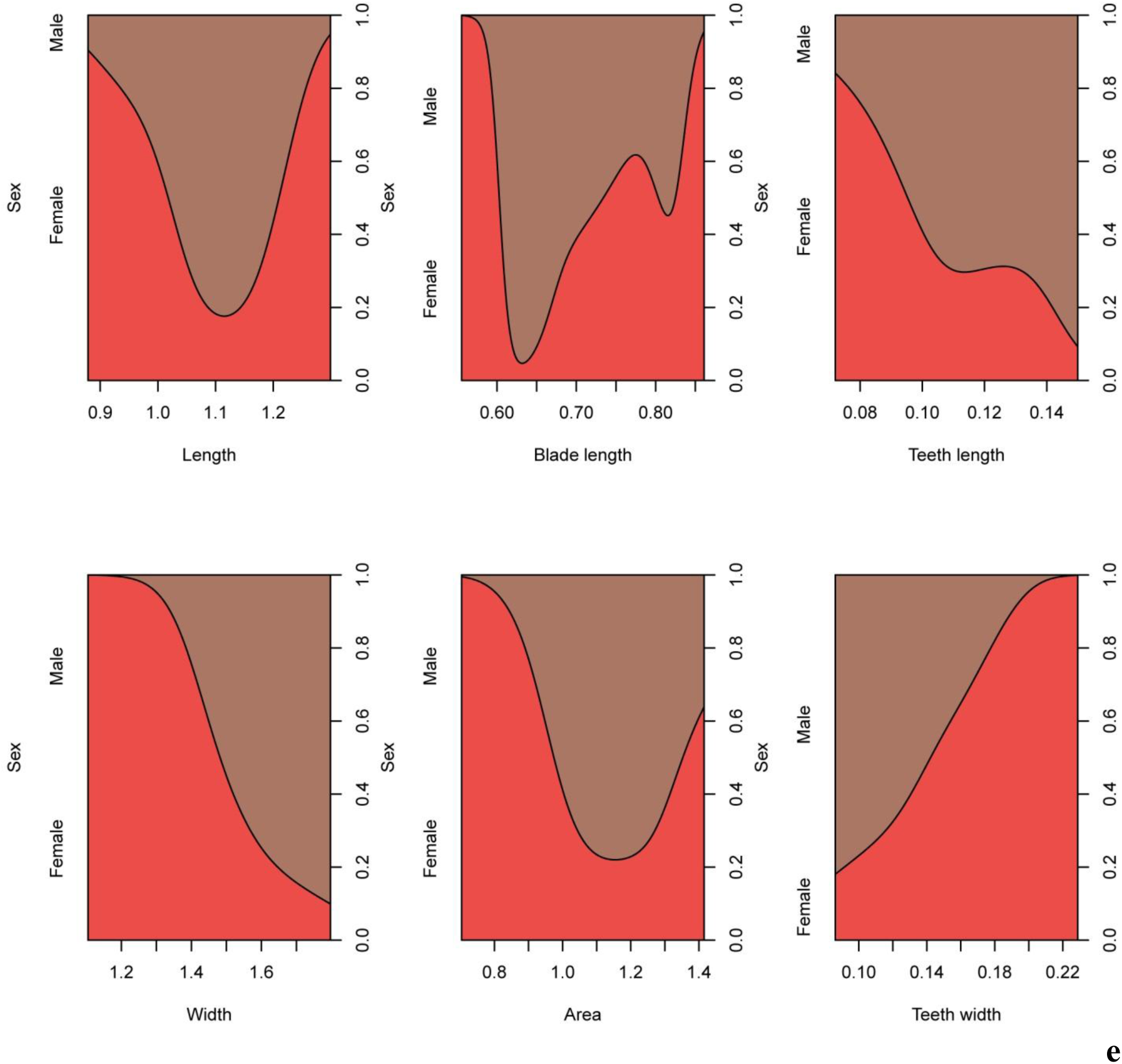
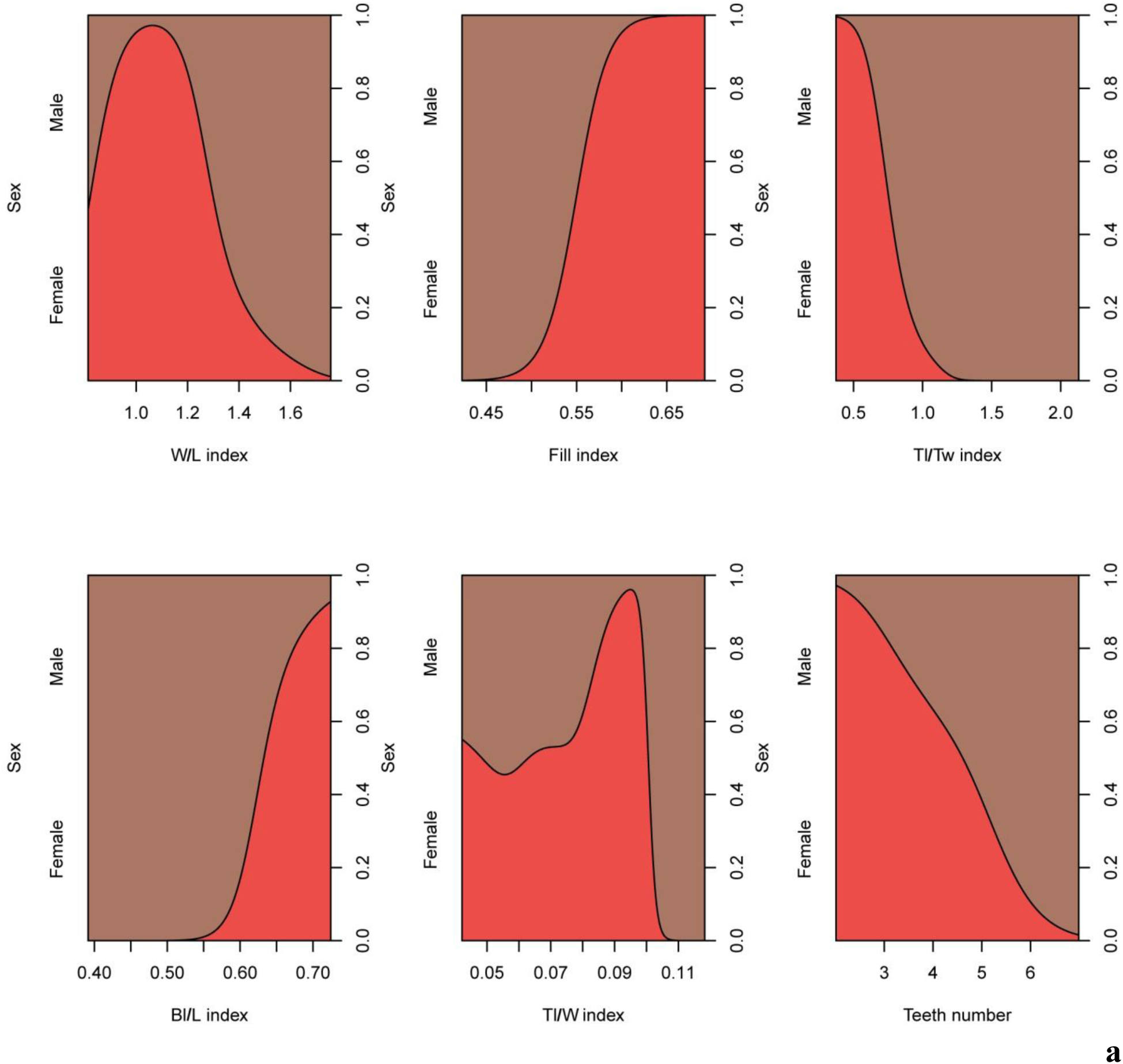
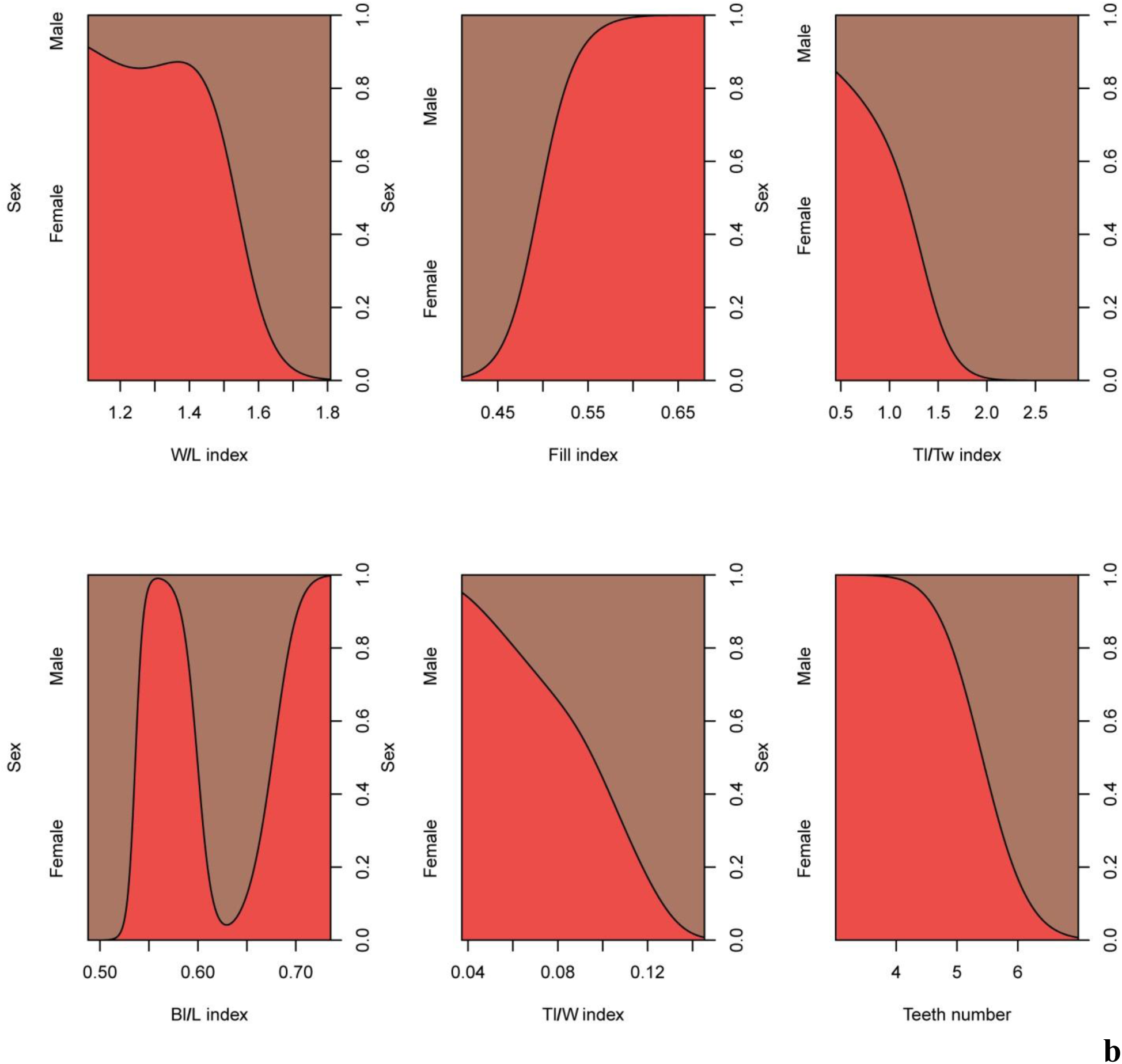
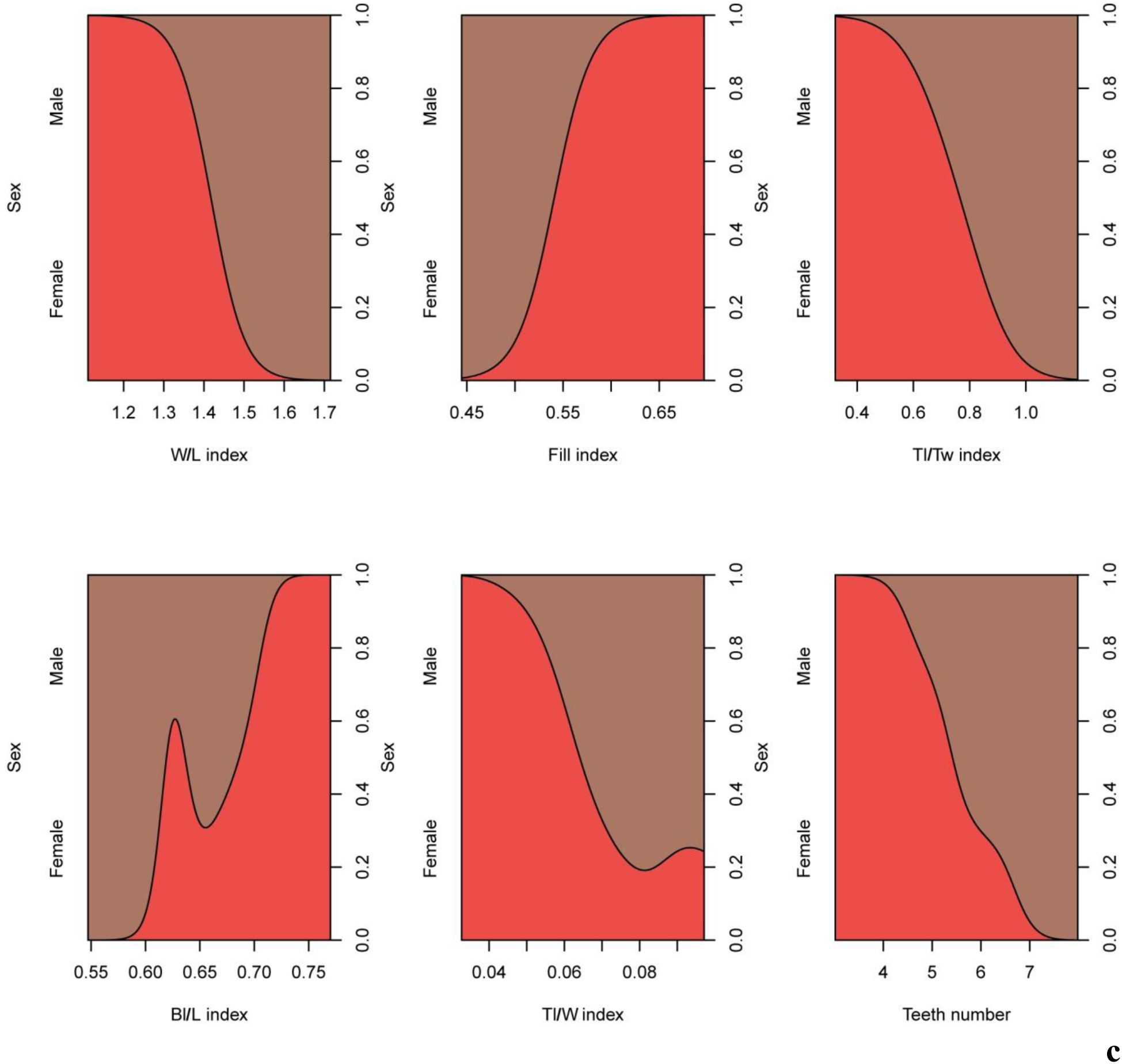
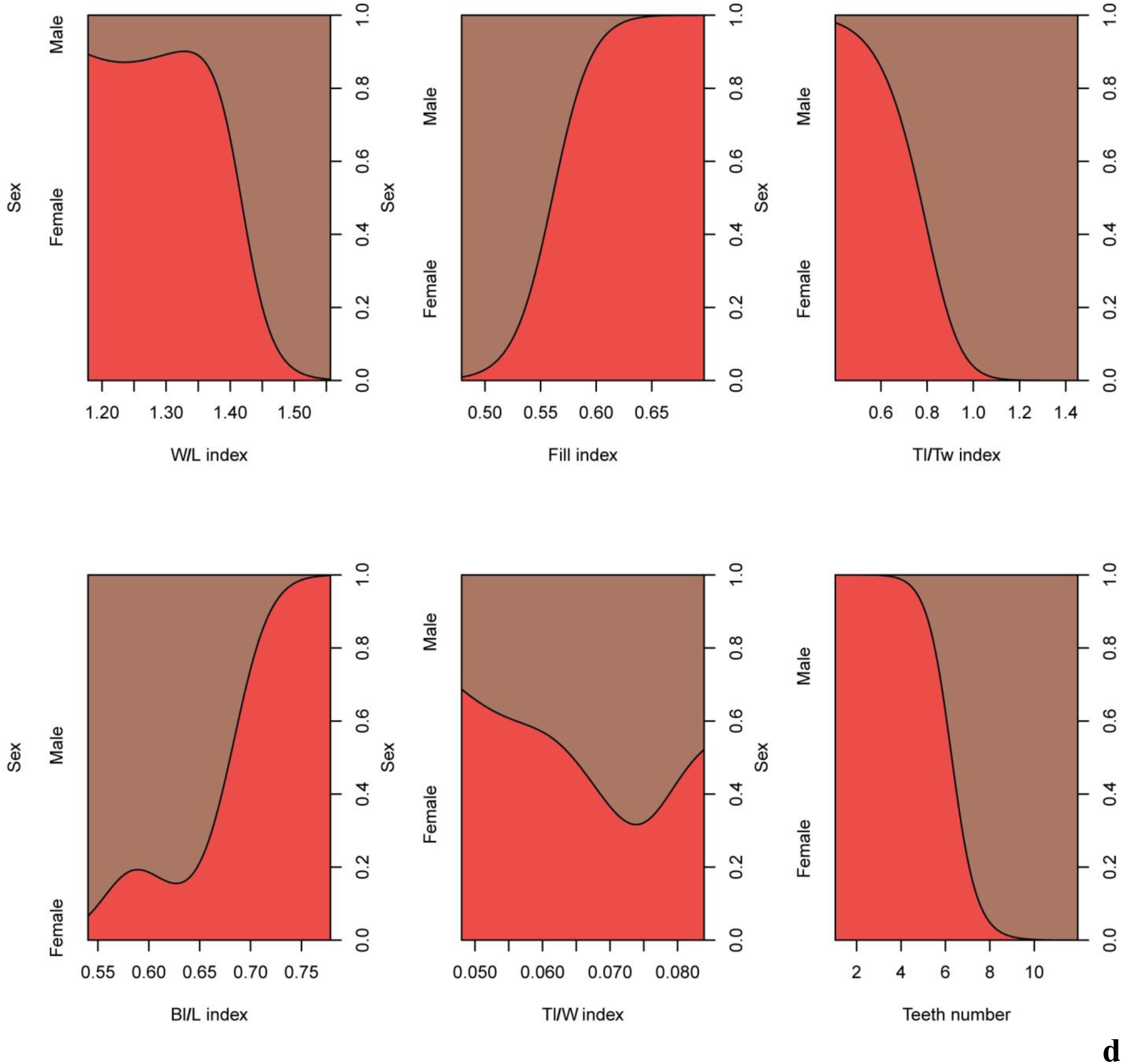
Conditional density plots of morphological parameters of the first (a), second (b), third (c), fourth (d), fifth (e) dorsal scutes of juveniles *Acipenser ruthenus*. Number of fish treated = 20. Number of scutes treated = 100.

**Fig. 16.**
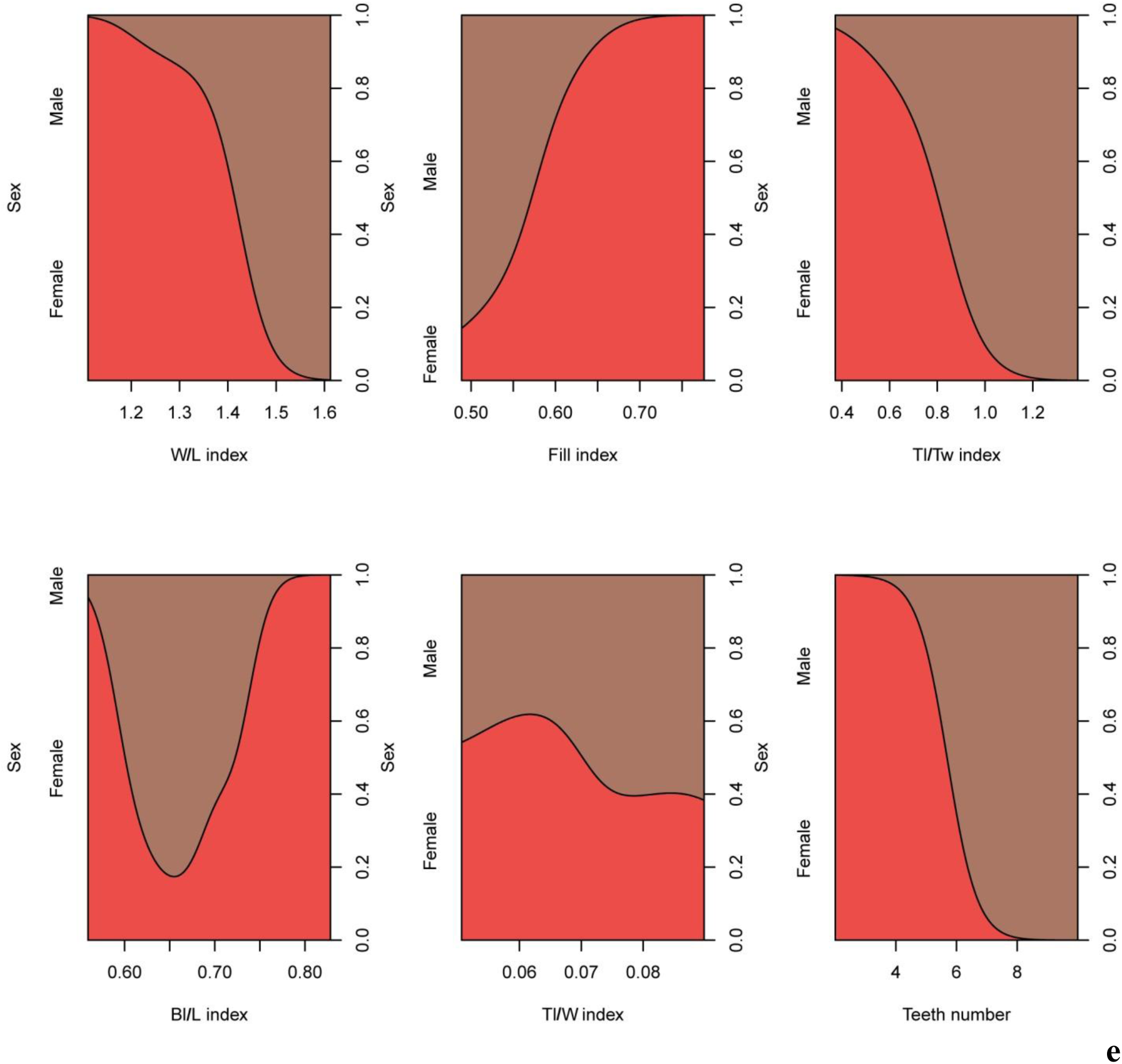
Conditional density plots of morphological indices of the first (a), second (b), third (c), fourth (d), fifth (e) dorsal scutes of juveniles *Acipenser ruthenus*. Number of fish treated = 20. Number of scutes treated = 100.

## DISCUSSION

We first found that sturgeon have external morphological differences depending on the sex, which was previously denied and considered impossible. We first found that sturgeon scutes (bone plates) have different forms (structure) depending on the sex. This dependence was preserved for sturgeons of all ages - from the juveniles (3-months-old) up to adults (2-years-old more). Therefore, sex determination is possible in the sturgeon early life periods without using of expensive equipment.

We found that differences between males and females of the sterlet dorsal scutes were up to 105.4 % for adults, 200.0 % for young, and 151.5 % for juveniles. Significant statistical differences remained in adults (2-years-old, length – 61.2±2.3 cm), and also in young (1-year-old, 24.8±1.5 cm) and juveniles (3 months-old, 70.3±3.6 mm) – future mature males and females.

Earlier, the sturgeon scutes were studied by other researchers in the field of morphological structure (Warth et al., 2017; Leprévost et al., 2017), environmental influence (Altenritter et al., 2015) and species identification (Thieren et al., 2015, Wuertz et al., 2011), but without studying the sex.

Discovered differences allow us to assert that the sterlet sex can be determined at the age of 2-3 months (length – 7-10 cm). However, the fish age is a relative indicator, i.e. the fish growth depends on the water temperature. For example, sterlet juveniles can reach up to 7-10 cm at the age of 5-6 months in pond farms.

We found that the morphological parameters of adult sterlet had a statistically significant decrease as a result of distance from the head (depending on the scute number) (Fig. 4, 17). Morphological indices increased depending on the scute number or did not change (Fig. 4, 18). We also found the correlation and other relationships in morphological parameters and indices in adult sterlet (Fig. 7, 10, 11, 12, 17). Similar correlation relationships were found in the young and juveniles of sterlet (Fig. 8, 9, 10, 12, 13, 14, 15, 16, 17).

**Fig. 17.**
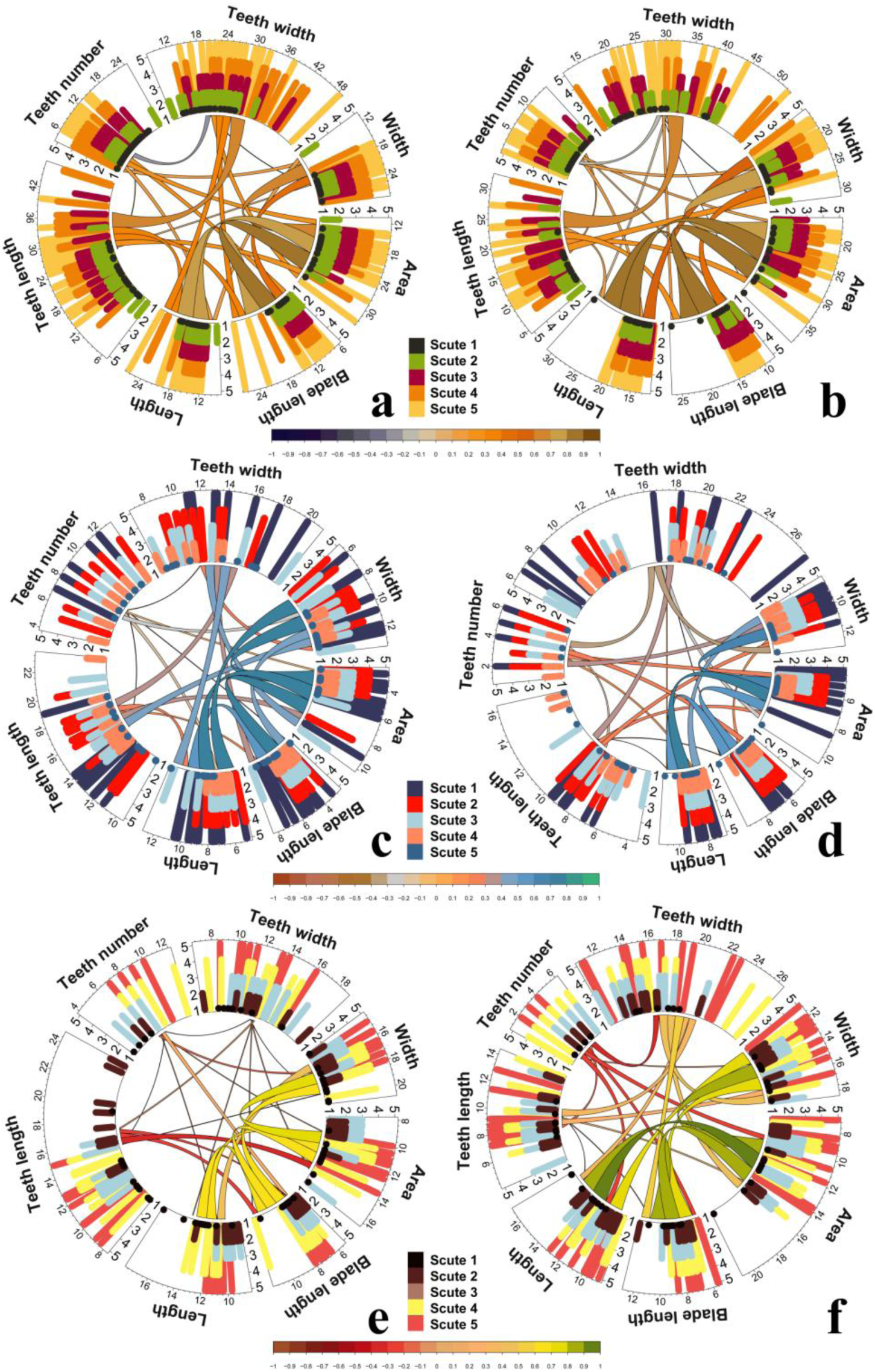
Circular representation of correlation relationship between all morphological parameters of *Acipenser ruthenus*. Outer circle shows differences between all morphological parameters (x-axis) from the first to the fifth dorsal scutes (y-axis) of adults (a, b), young (c, d) and juveniles (e, f) depending on sex (a, b, c, d, e, f) and scute number (y-axis). Inside circle shows correlation relationship between all morphological parameters. Line color and thickness represent the strength of correlation relationship. Number of fish treated = 37 (adults), 20 (young), 20 (juveniles). Number of scutes treated = 185 (adults), 100 (young), 100 (juveniles).

**Fig. 18.**
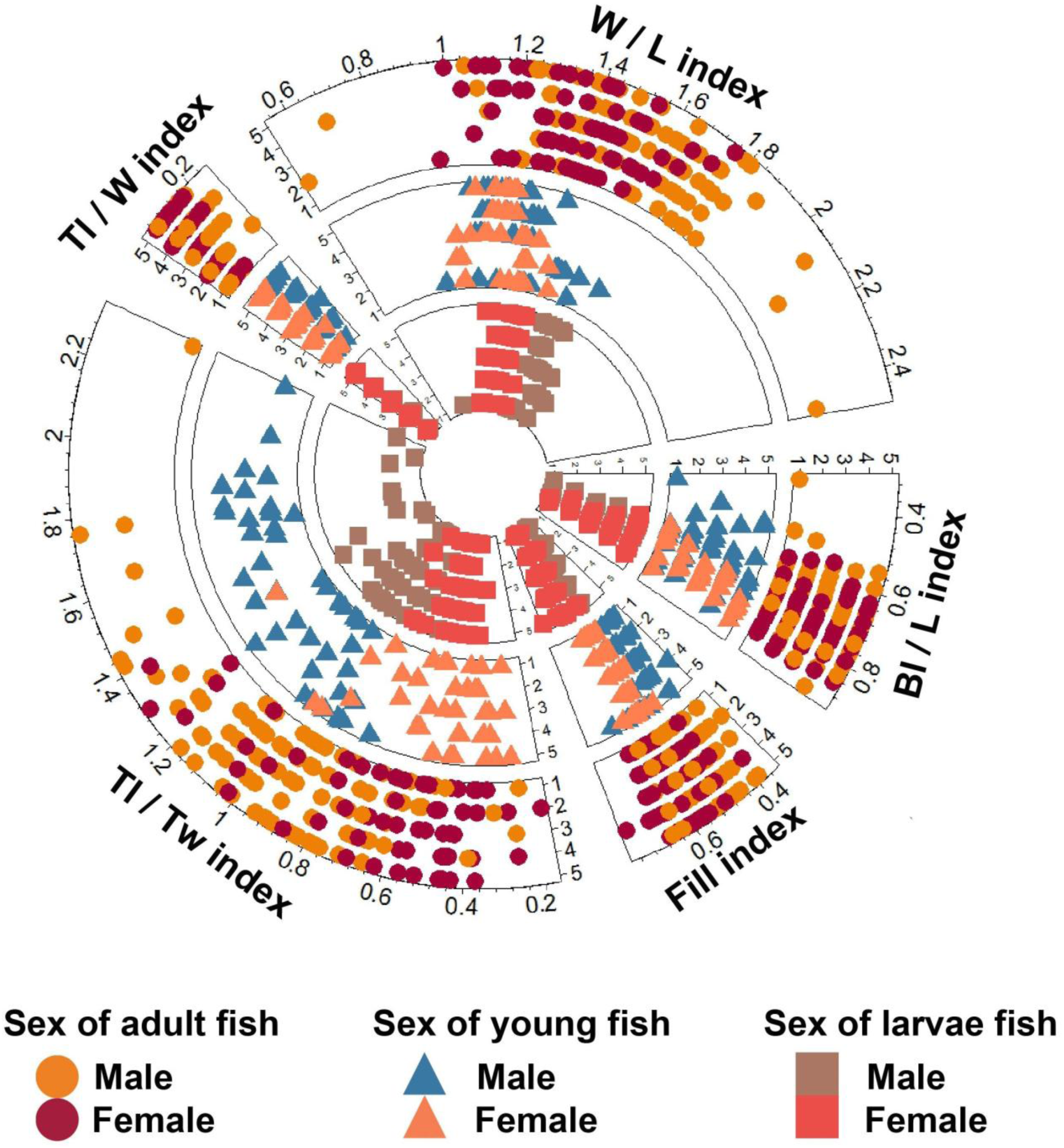
Circular represent differences between all morphological indices (x-axis) from the first to the fifth dorsal scutes of adults (outer circle), young (middle circle) and juveniles (inner circle) depending on sex and scute number (y-axis). Number of fish treated = 37 (adults), 20 (young), 20 (juveniles). Number of scutes treated = 185 (adults), 100 (young), 100 (juveniles).

Morphological parameters are not permanent parameters that depend on size and age of sterlet sturgeon. Therefore, important parameters for sex determination are the relative morphological parameters (indicators) which do not depend on the mass and age.

Two groups of indicators that characterized visual perceptions of scute forms were formed: 1. Group of morphological indices characterizing the dorsal scute shape: scute width / scute length (W/L) index, average blade length / scute length (Bl/L) index, fill index; 2. Group of morphological indices characterizing the teeth of dorsal scute: teeth length / scute width (Tl/W) index, teeth length / teeth width (Tl/Tw) index.

Next, we selected most significant morphological parameters and indices for sex determination, using the machine learning methods (random forrest method (Paluszynska, Biecek, 2017), boruta algorithm (Kursa, Rudnicki, 2010), neural networks (Fritsch, Guenther, 2016) (Fig 19, 20, 21). Selected parameters were tested for multicorrelation, and then a model was created that allows us to statistically significant determine sex using building trees method based on recursive partitioning (Milborrow, 2017) (Fig. 22, 23, 24 Tabl. 1).

**Fig. 19.**
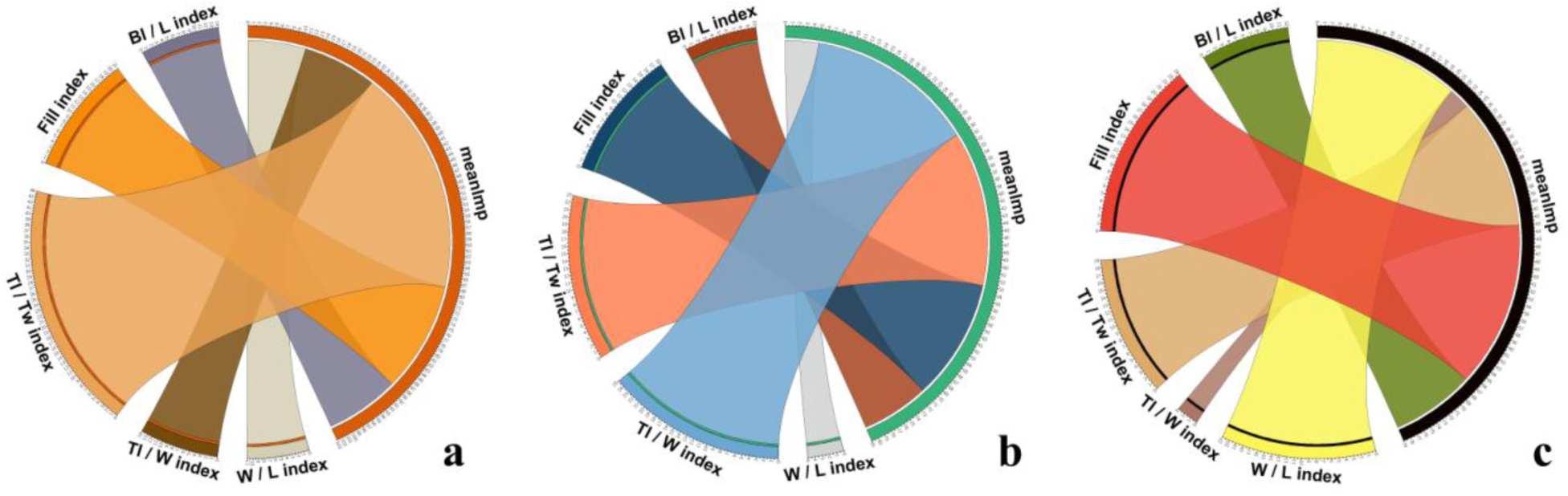
Selecting important morphological indices to create a sex determination model of *Acipenser ruthenus*. Circles (Krzywinski et al., 2009) (a-c) show the results of morphological indices assessment using the boruta algorithm for adults (a), young (b) and juveniles (c). Number of fish treated = 37 (adults), 20 (young), 20 (juveniles). Number of scutes treated = 370 (adults), 200 (young), 200 (juveniles).

**Fig. 20.**
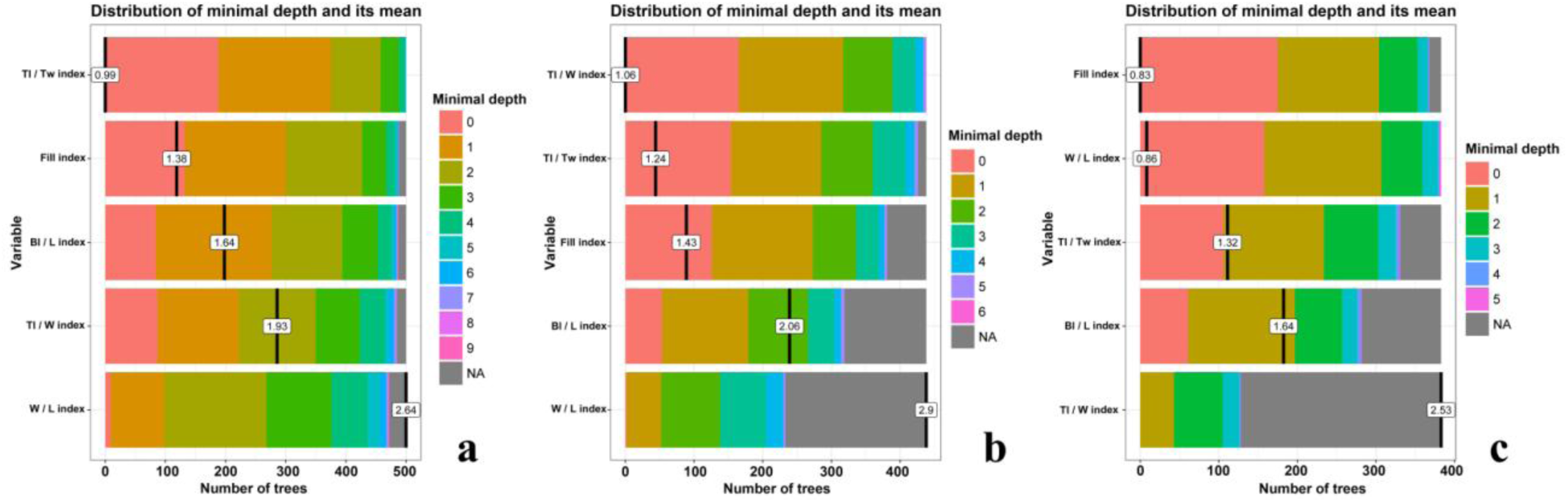
Selecting important morphological indices to create a sex determination model of *Acipenser ruthenus*. Graphs (a-c) show the results of morphological indices assessment using the random forrest method for adults (a), young (b) and juveniles (c). Number of fish treated = 37 (adults), 20 (young), 20 (juveniles). Number of scutes treated = 370 (adults), 200 (young), 200 (juveniles).

**Fig. 21.**
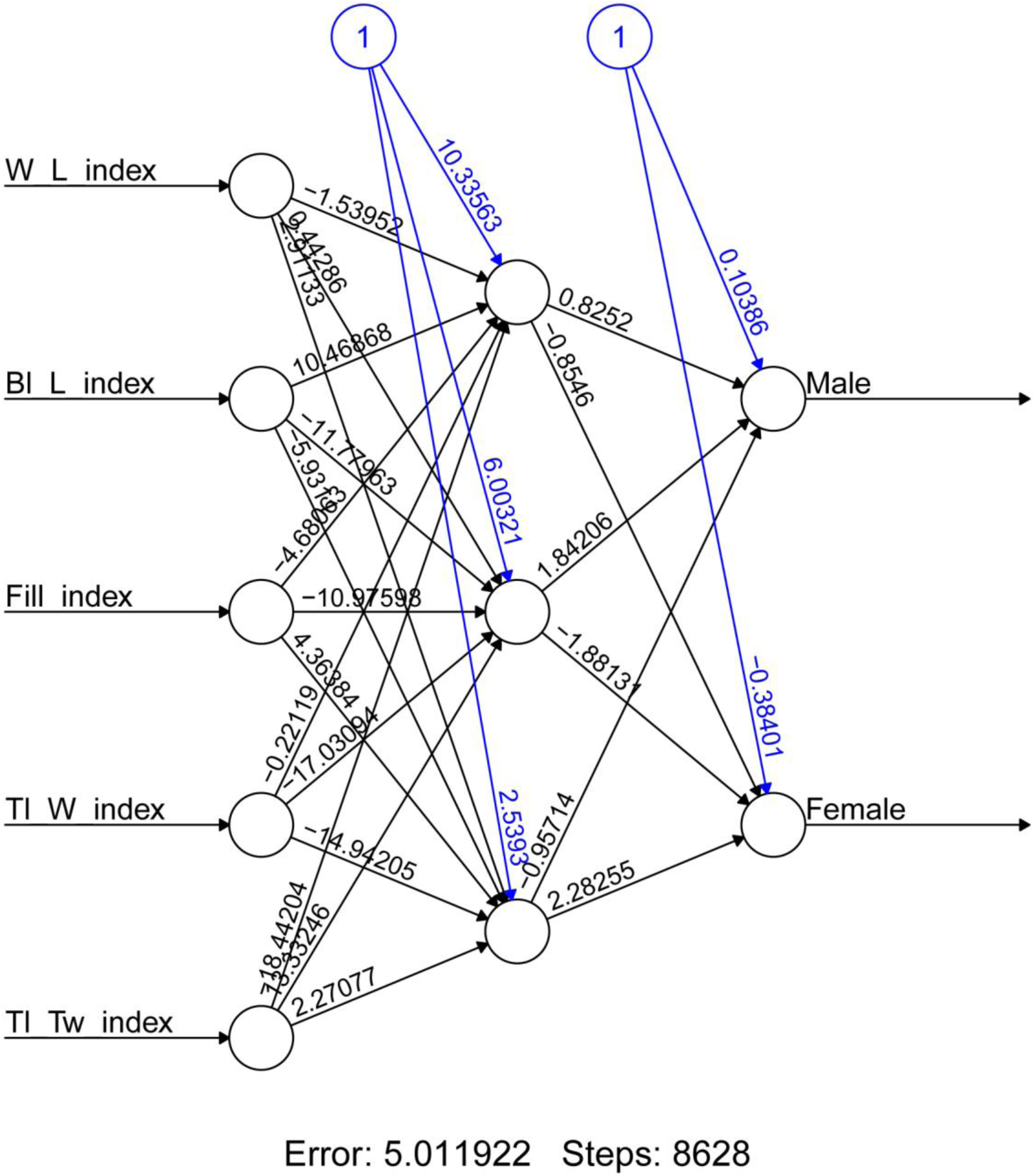
Example of trained neural network for sex identification of *Acipenser ruthenus*

**Fig. 22.**
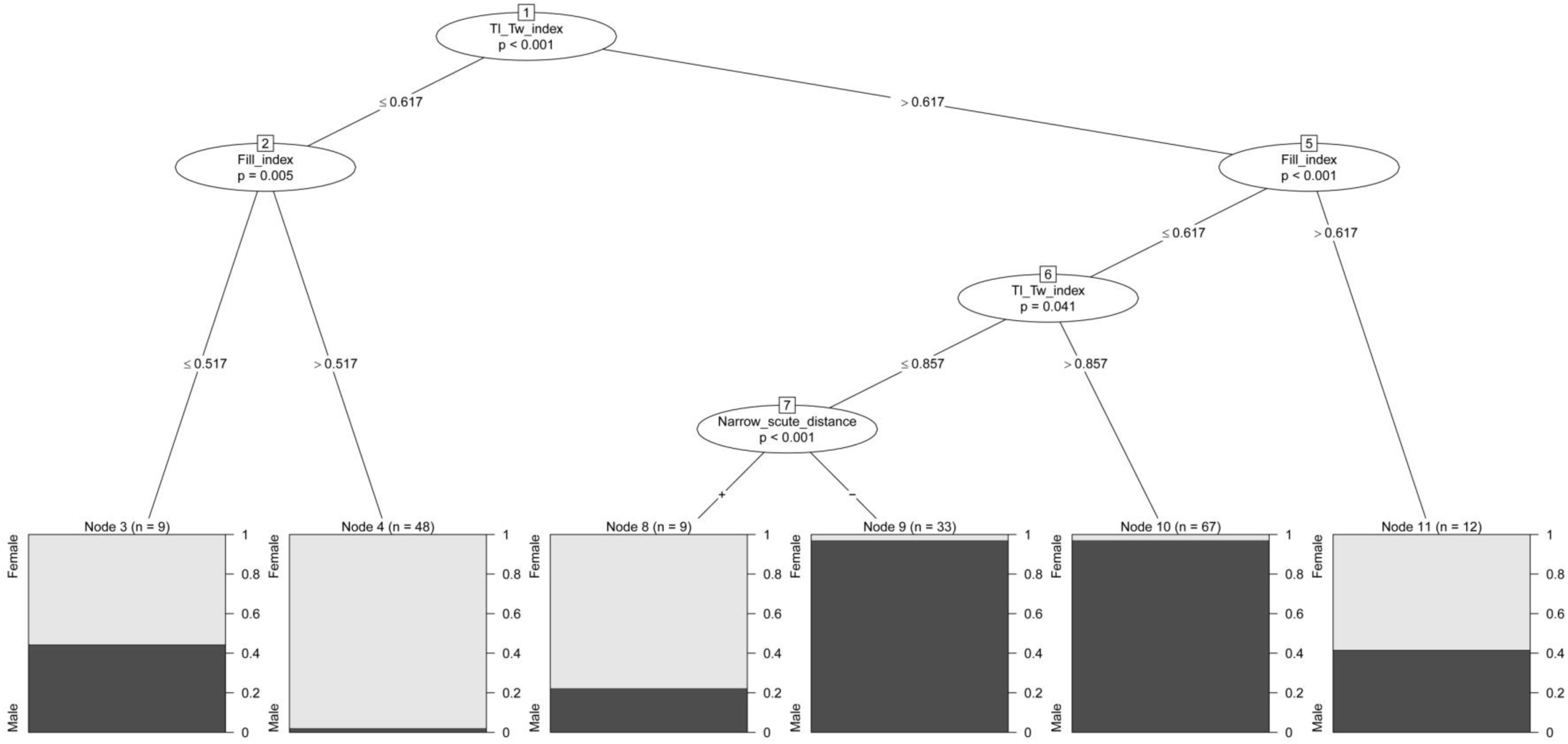
Decision tree model for sex identification of adults *Acipenser ruthenus*

**Fig. 23.**
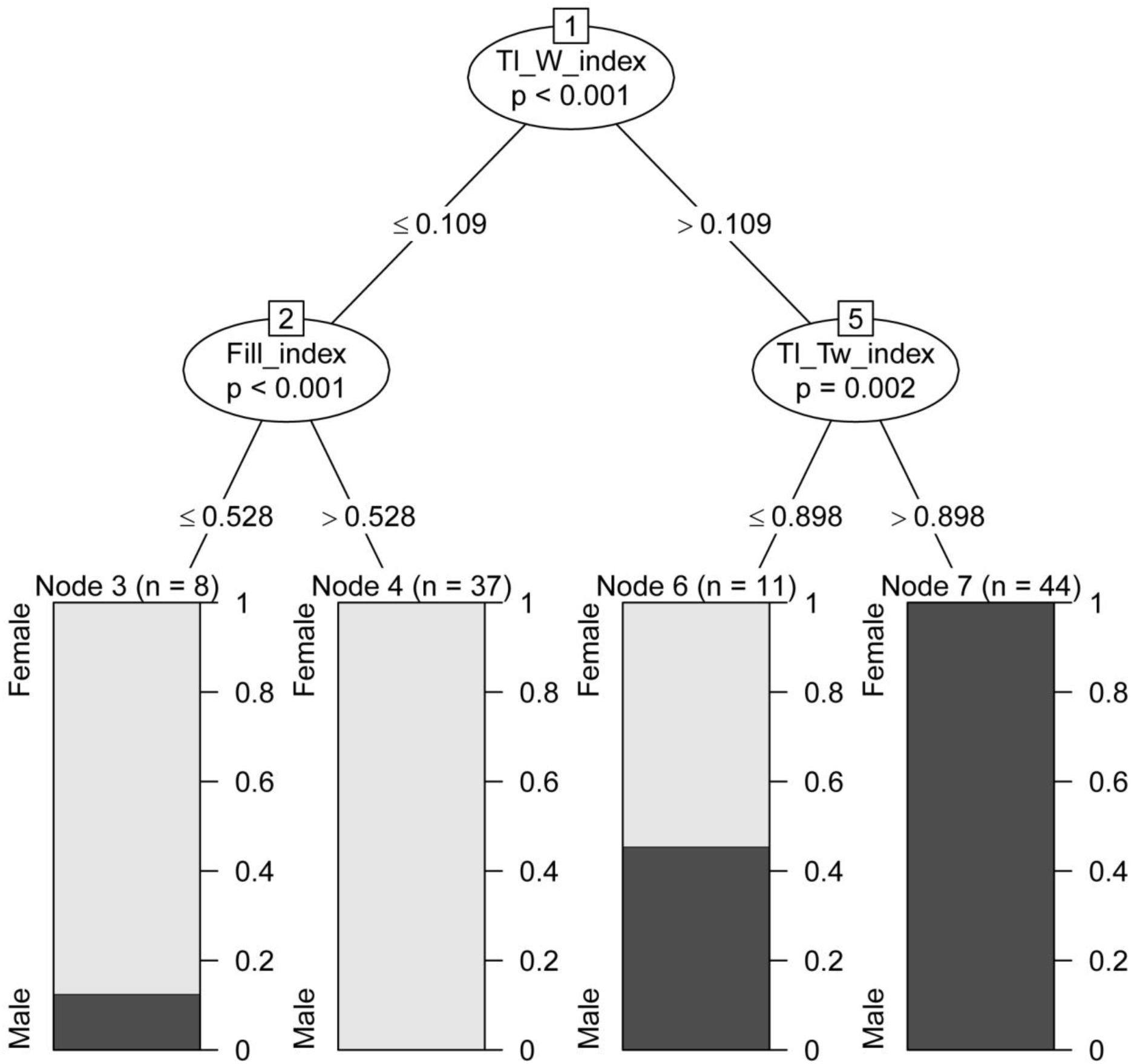
Decision tree model for sex identification of young *Acipenser ruthenus*

**Fig. 24.**
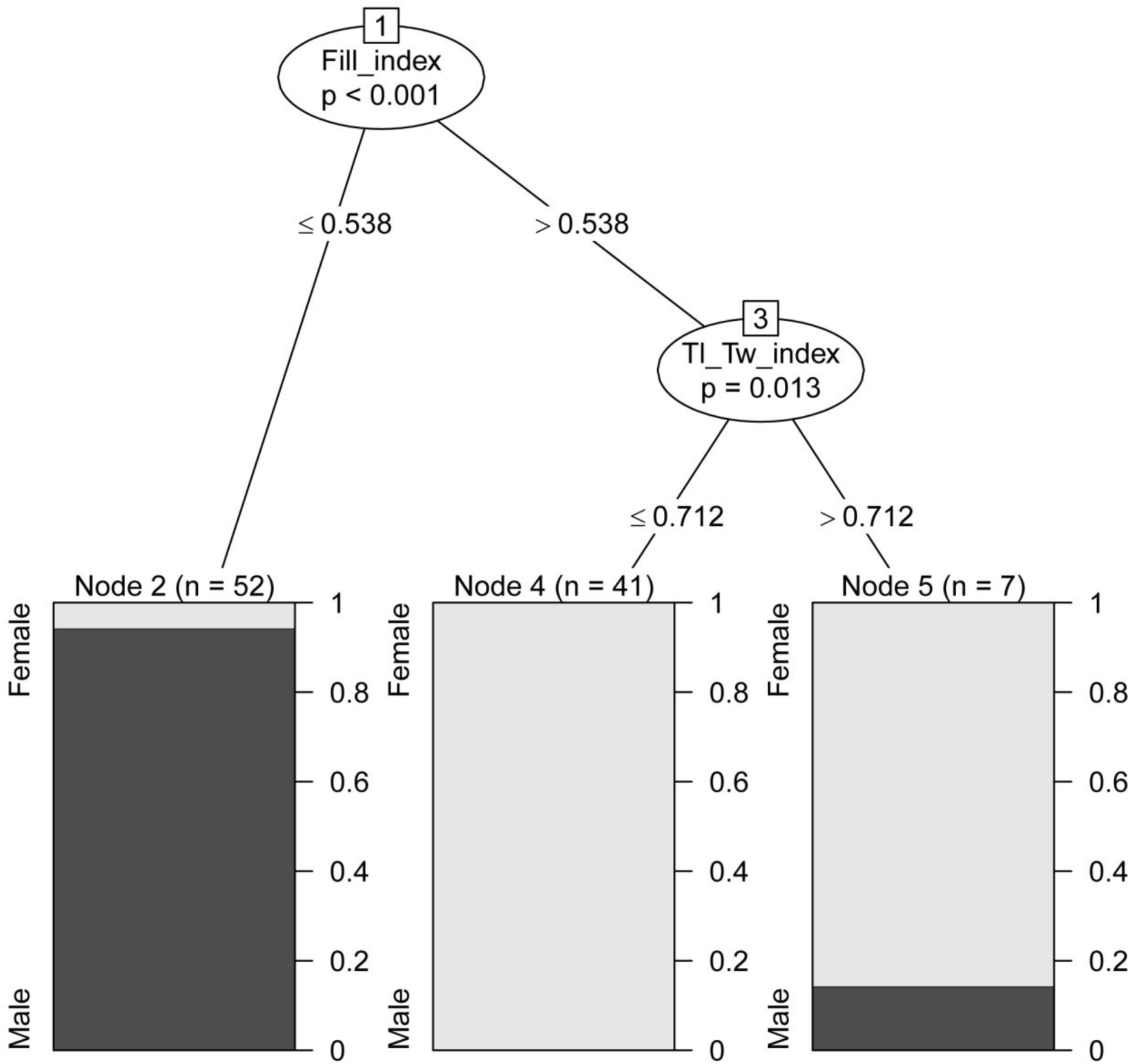
Decision tree model for sex identification of juveniles *Acipenser ruthenus*

**Table 1.** Cumulative logit model with the best regression (Akaike – criterion) for the adult -, young – juvenile dorsal scutes *of Acipenser ruthenus*

|  | Coefficients |  |  |
| --- | --- | --- | --- |
|  | Value | Std. Error | t value |
| W / L index | -1.959 | 0.6378 | -3.07 |
| Bl / L index | -7.972 | 1.487 | -5.36 |
| Fill index | -17.703 | 2.0689 | -8.56 |
| Teeth number | 0.048 | 0.0319 | 1.5 |
| Tl / W index | 9.583 | 3.4359 | 2.79 |
| Tl / Tw index | 2.531 | 0.4363 | 5.8 |

| Re-fitting to get Hessian |  |  |
| --- | --- | --- |
|  | 2.50% | 97.50% |
| W / L index | -3.20881 | -0.70869 |
| Bl / L index | -10.8865 | -5.05743 |
| Fill index | -21.7581 | -13.6483 |
| Teeth number | -0.01454 | 0.110466 |
| Tl / W index | 2.849207 | 16.31755 |
| Tl / Tw index | 1.67639 | 3.386471 |

The discrepancy between the empirical and predicted values of sex determination
|  |  | Prediction |  |
| --- | --- | --- | --- |
|  |  | Female | Male |
| Empirical values | Female | 133 | 36 |
|  | Male | 10 | 199 |
| % Error in females |  | 21.30178 | - |
| % Error in males |  | - | 4.784689 |

Next we found, that quantitative change in the selected morphological parameters and indices affect on the visual perception of scutes forms depending on the sex. We analyzed occurrence frequency of various scute forms and found significant differences between males and females of adult sterlet. Significant differences were spotted on the average from the first to the fifth dorsal scutes (Fig. 25, 26, 27, 28, Tabl. 2). Similar differences were found in young sterlet (Fig. 29, 30, 31, 32, Tabl. 3). Similar differences were also found in juveniles (Fig. 33, 34, 35, 36 Tabl. 4) but only when studying cleaned scutes of dead juveniles. In vivo, juvenile scutes of future males were significantly different from scutes of future females due to presence of small and sharp teeth on scute surface. The juvenile scutes of future males have also significant curvature at the top (Fig. 37).

**Fig. 25.**
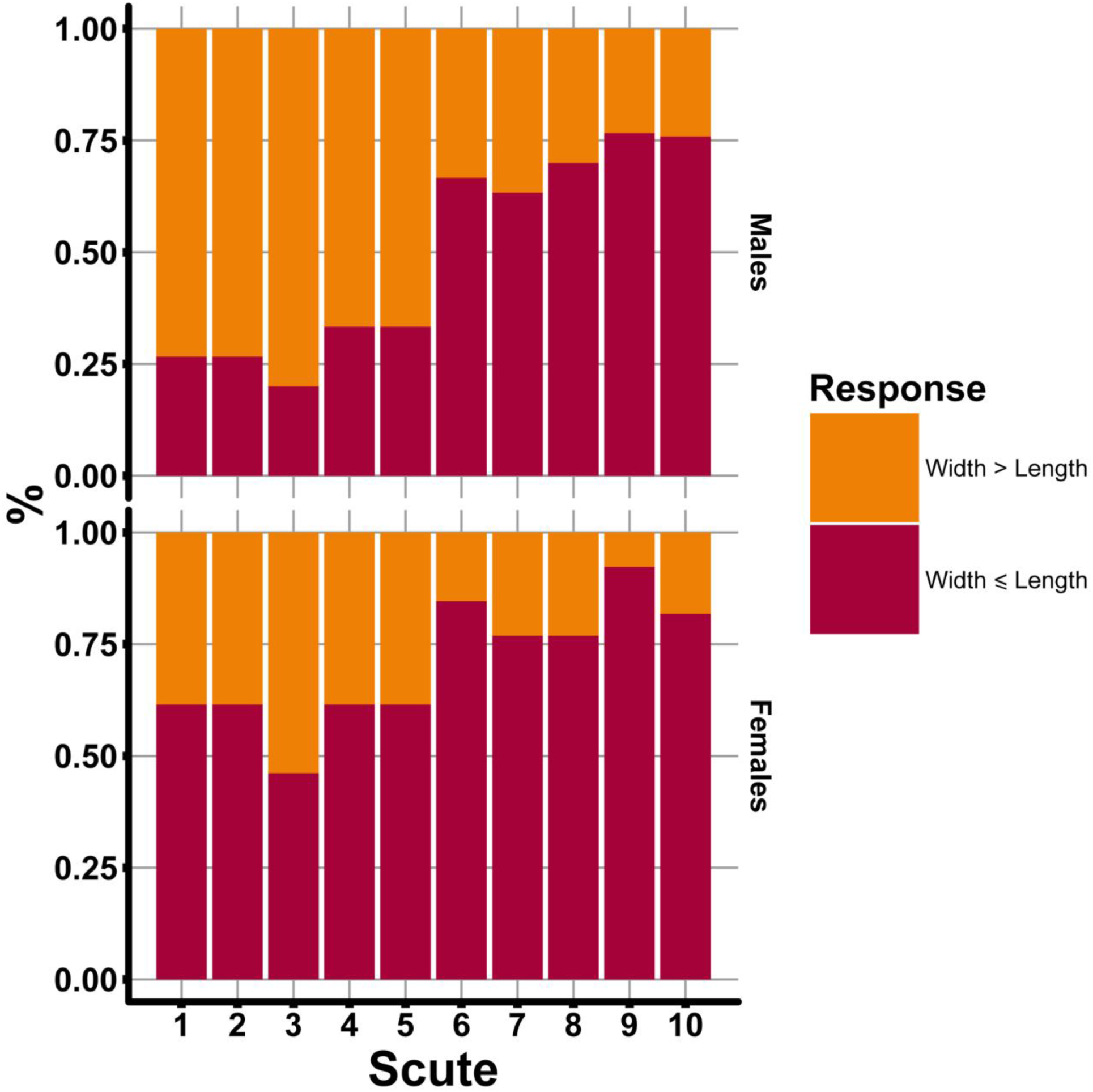
Plot show result of chi-square (χ^2^) test for visual perception of relationship between width (W) and length (L) of dorsal scutes of adults *Acipenser ruthenus*. Number of fish treated = 40-43.

**Fig. 26.**
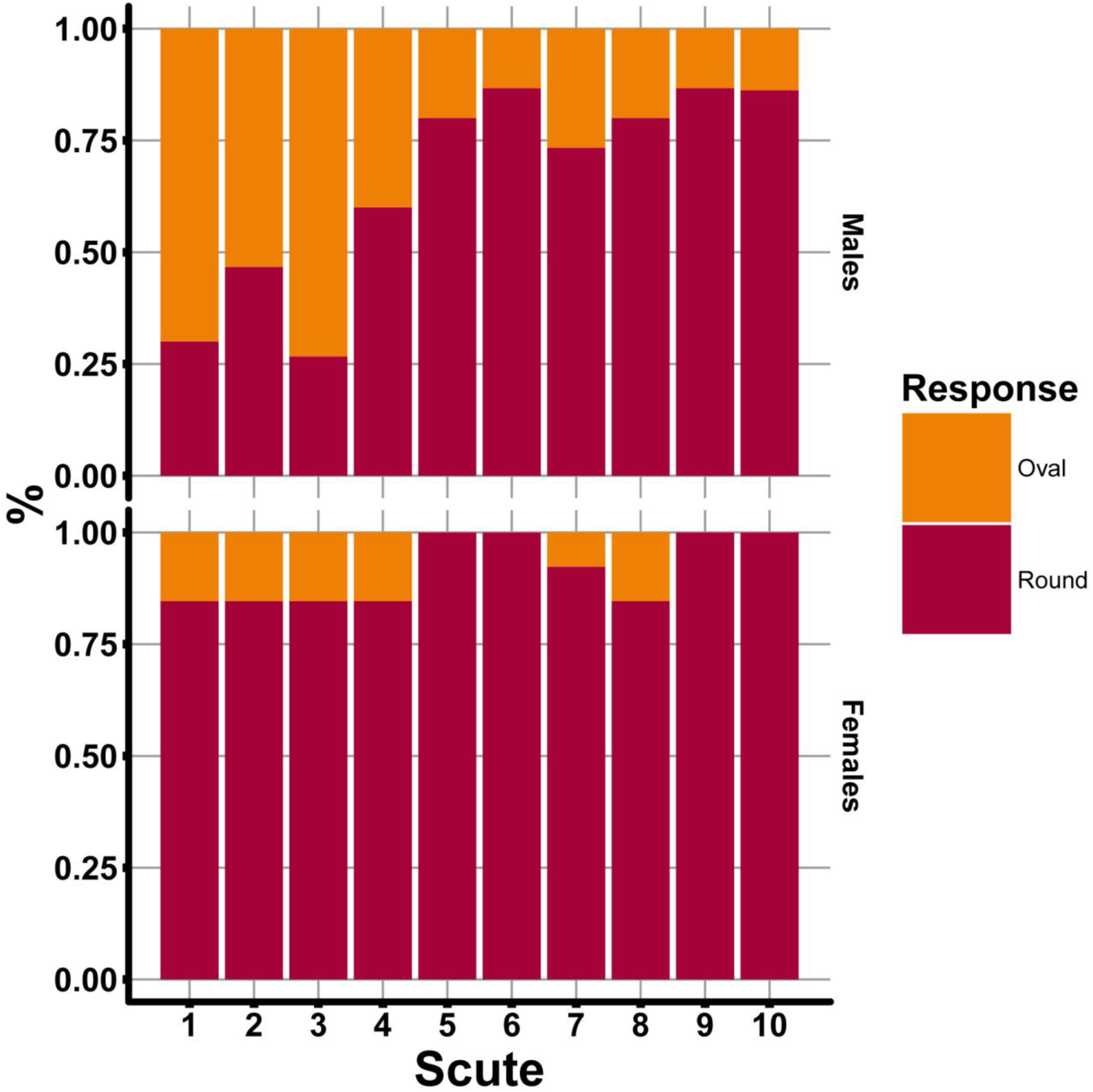
Plot show result of chi-square (χ^2^) test for visual perception of dorsal scute shape of adults *Acipenser ruthenus*. Number of fish treated = 40-43.

**Fig. 27.**
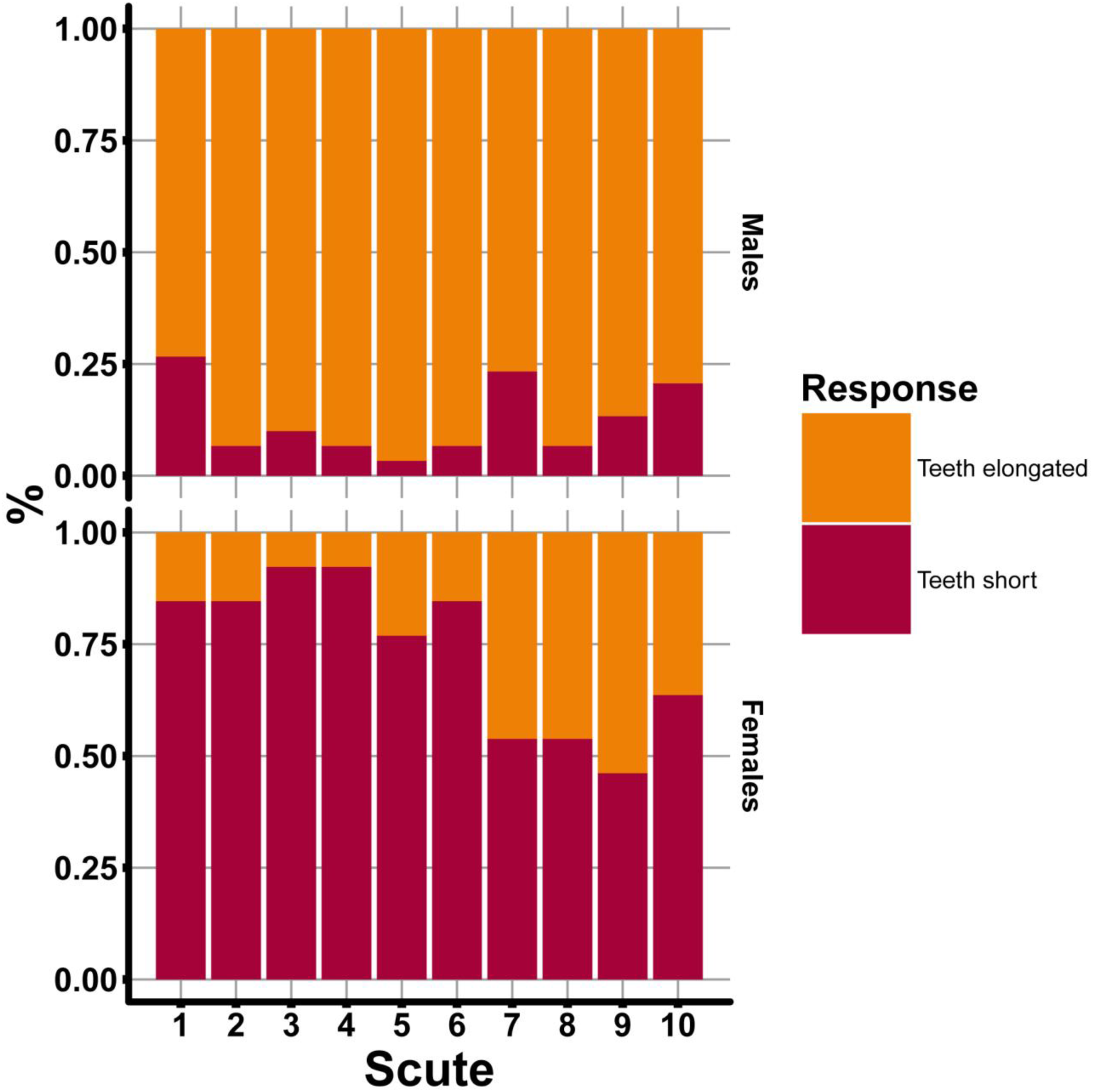
Plot show result of chi-square (χ^2^) test for visual perception of length of the dorsal scute teeth of adults *Acipenser ruthenus*. Number of fish treated = 40-43.

**Fig. 28.**
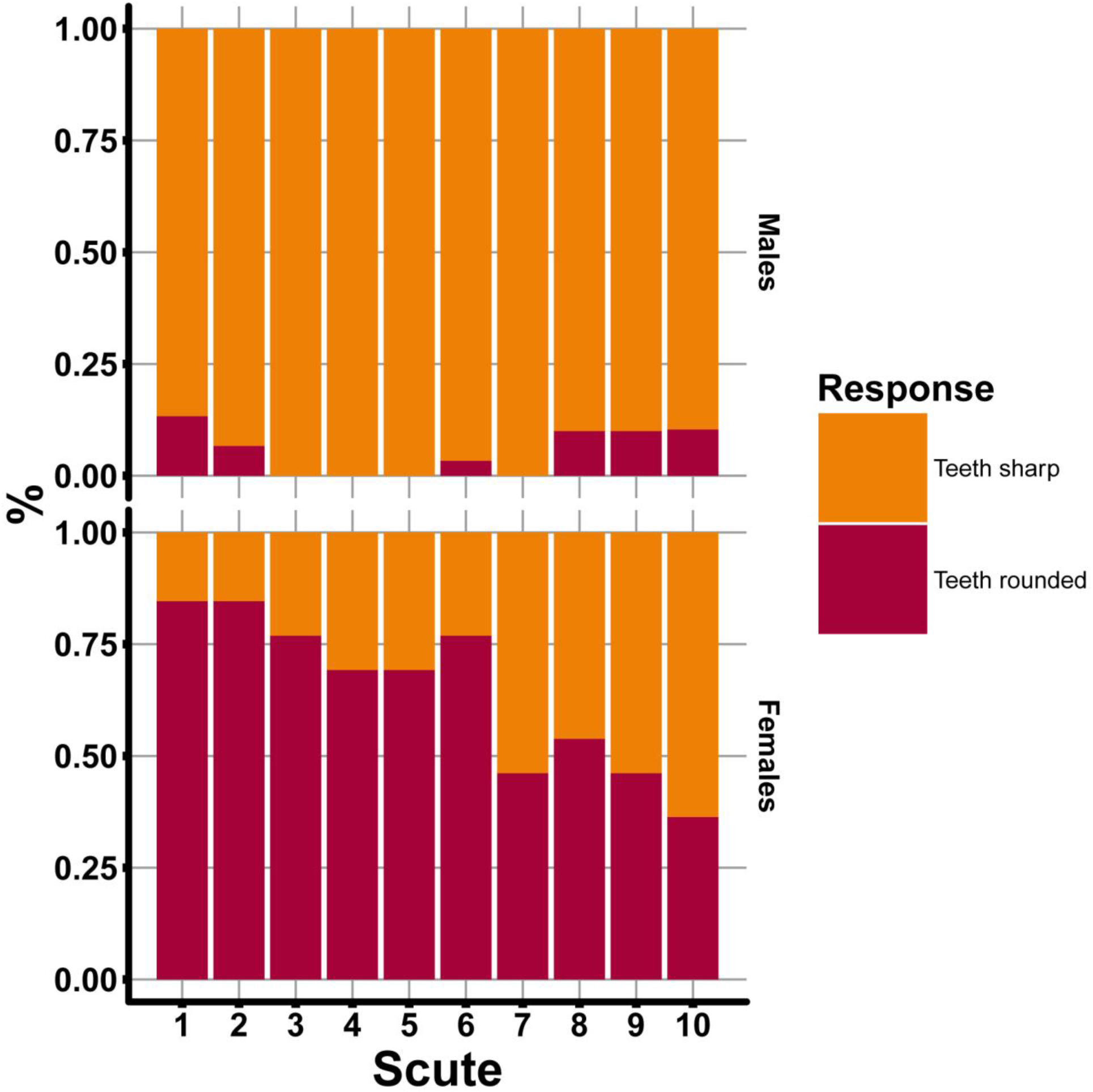
Plot show result of chi-square (χ^2^) test for visual perception of shape of the dorsal scute teeth of adults *Acipenser ruthenus*. Number of fish treated = 40-43.

**Fig. 29.**
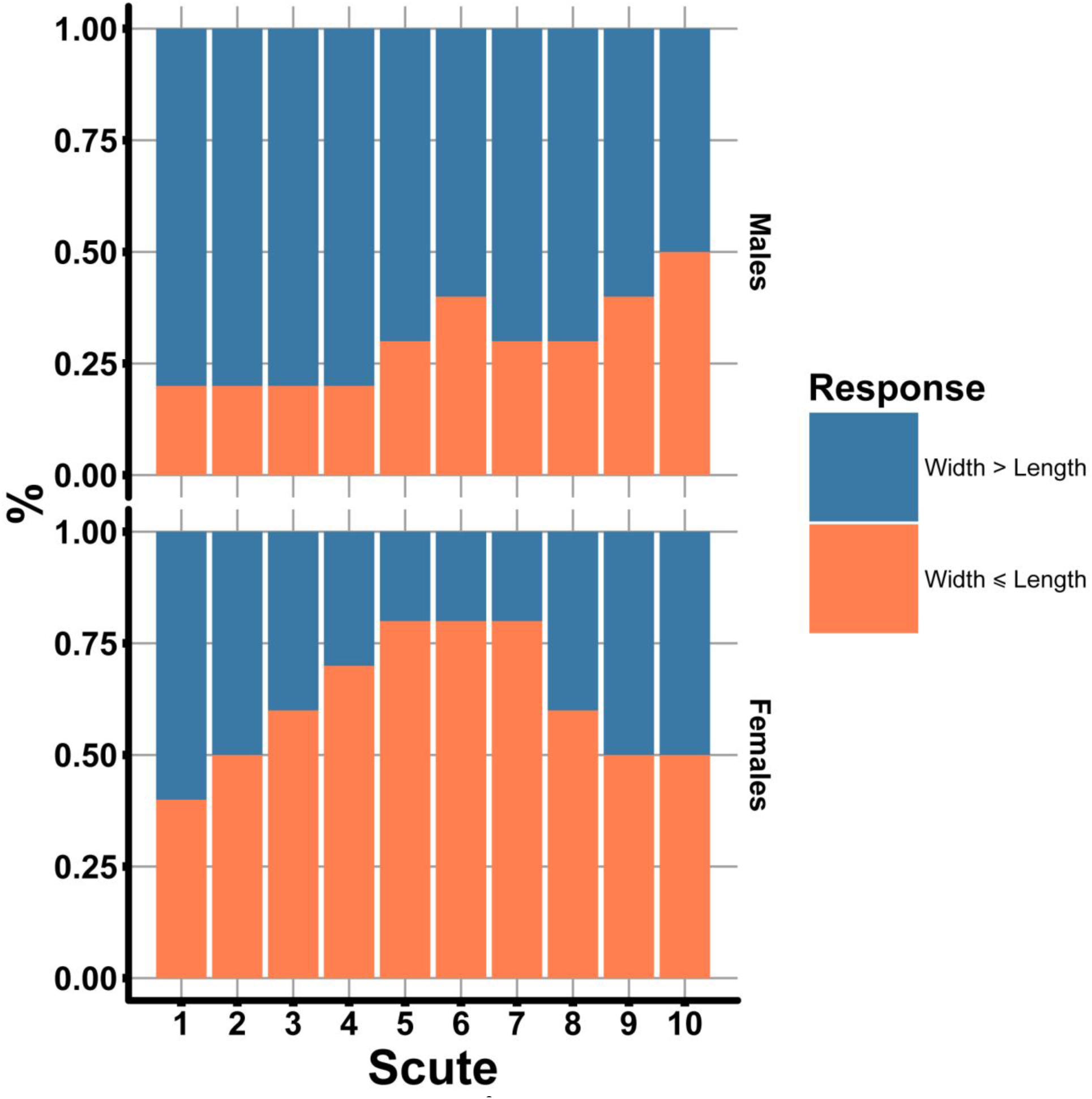
Plot show result of chi-square (χ^2^) test for visual perception of relationship between width (W) and length (L) of dorsal scutes of young *Acipenser ruthenus* (Table S3). Number of fish treated = 20.

**Fig. 30.**
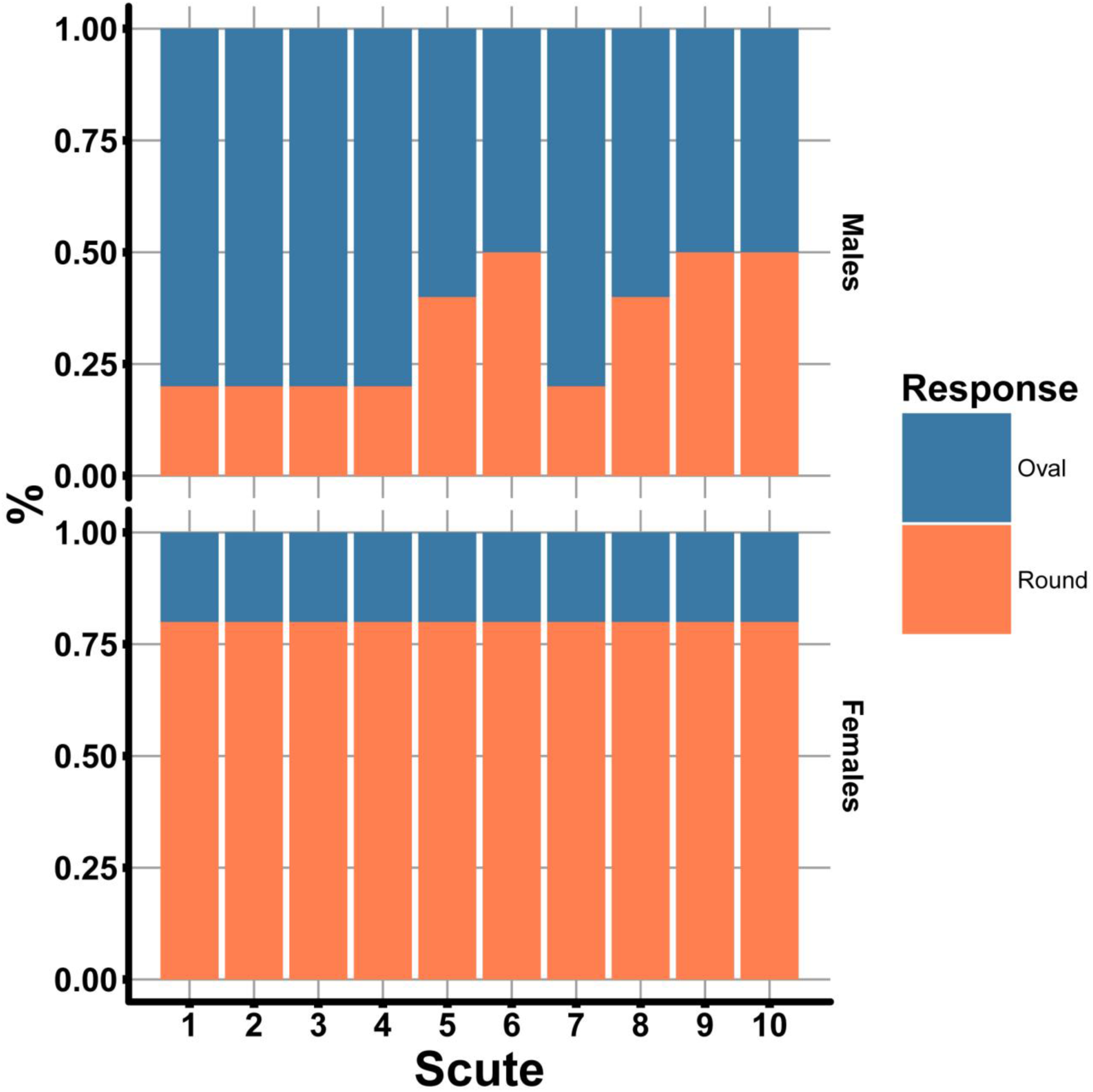
Plot show result of chi-square (χ^2^) test for visual perception of dorsal scute shape of young *Acipenser ruthenus*. Number of fish treated = 20.

**Fig. 31.**
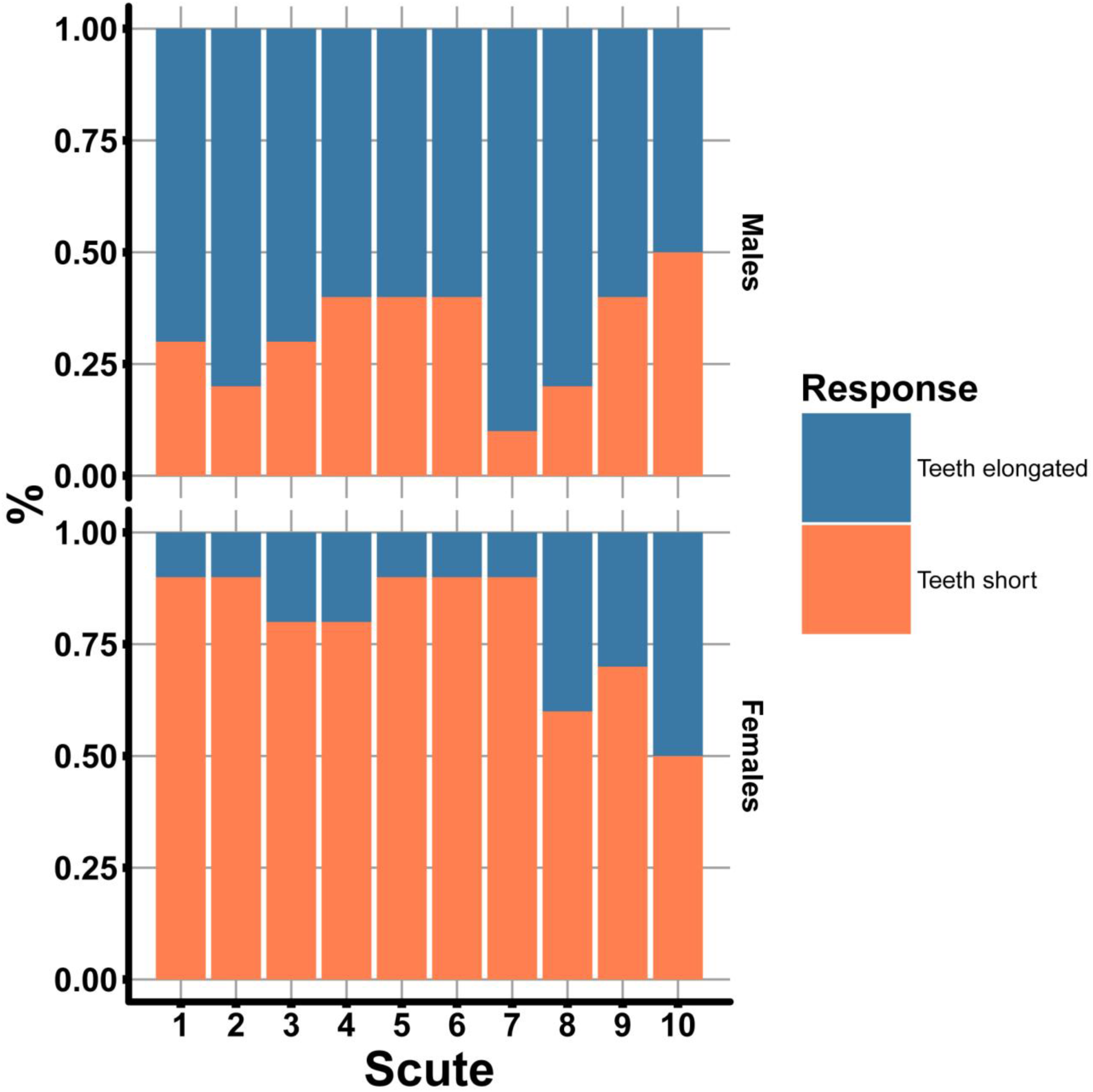
Plot show result of chi-square (χ^2^) test for visual perception of length of the dorsal scute teeth of young *Acipenser ruthenus*. Number of fish treated = 20.

**Fig. 32.**
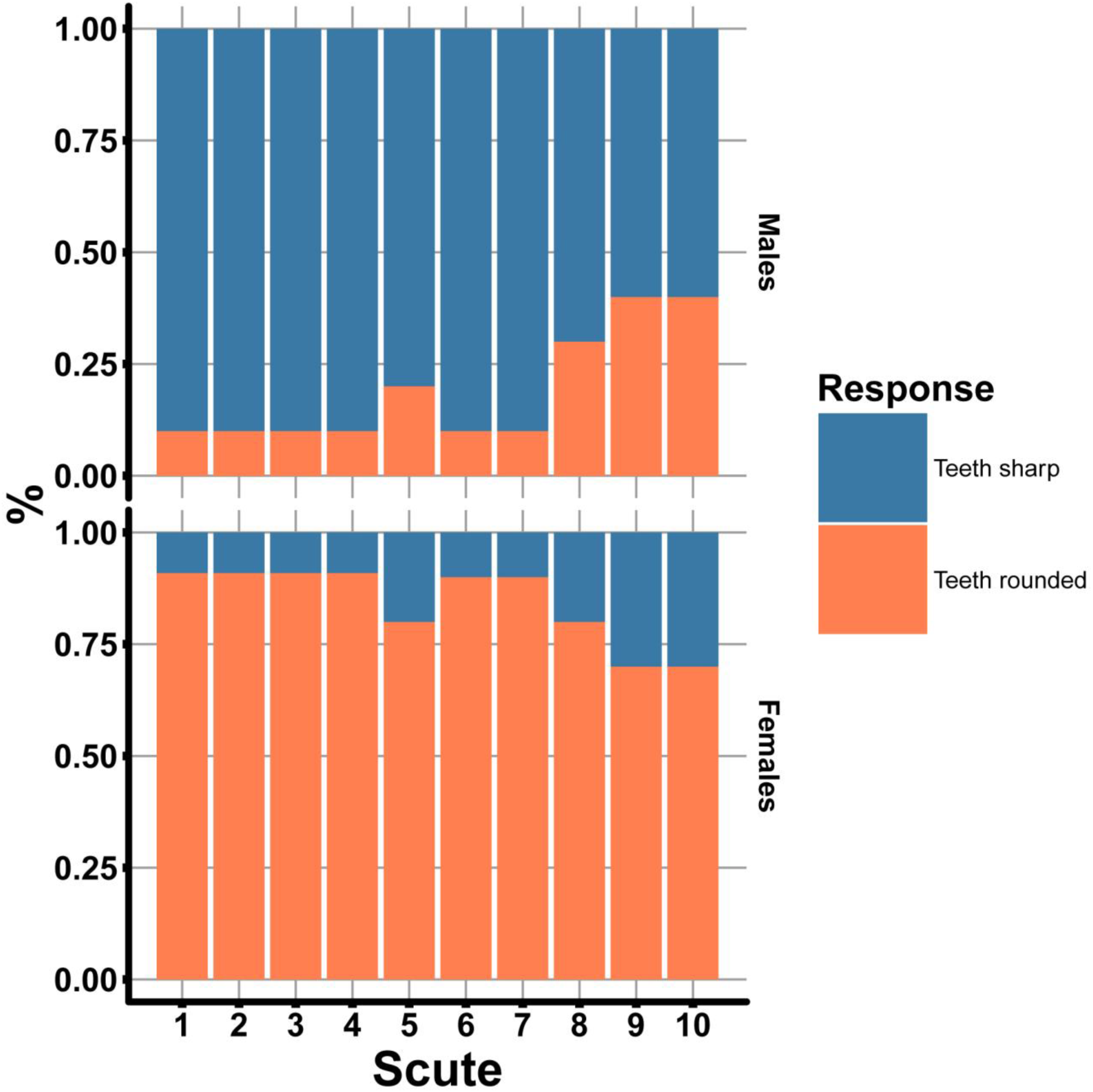
Plot show result of chi-square (χ^2^) test for visual perception of shape of the dorsal scute teeth of adults *Acipenser ruthenus*. Number of fish treated = 20.

**Fig. 33.**
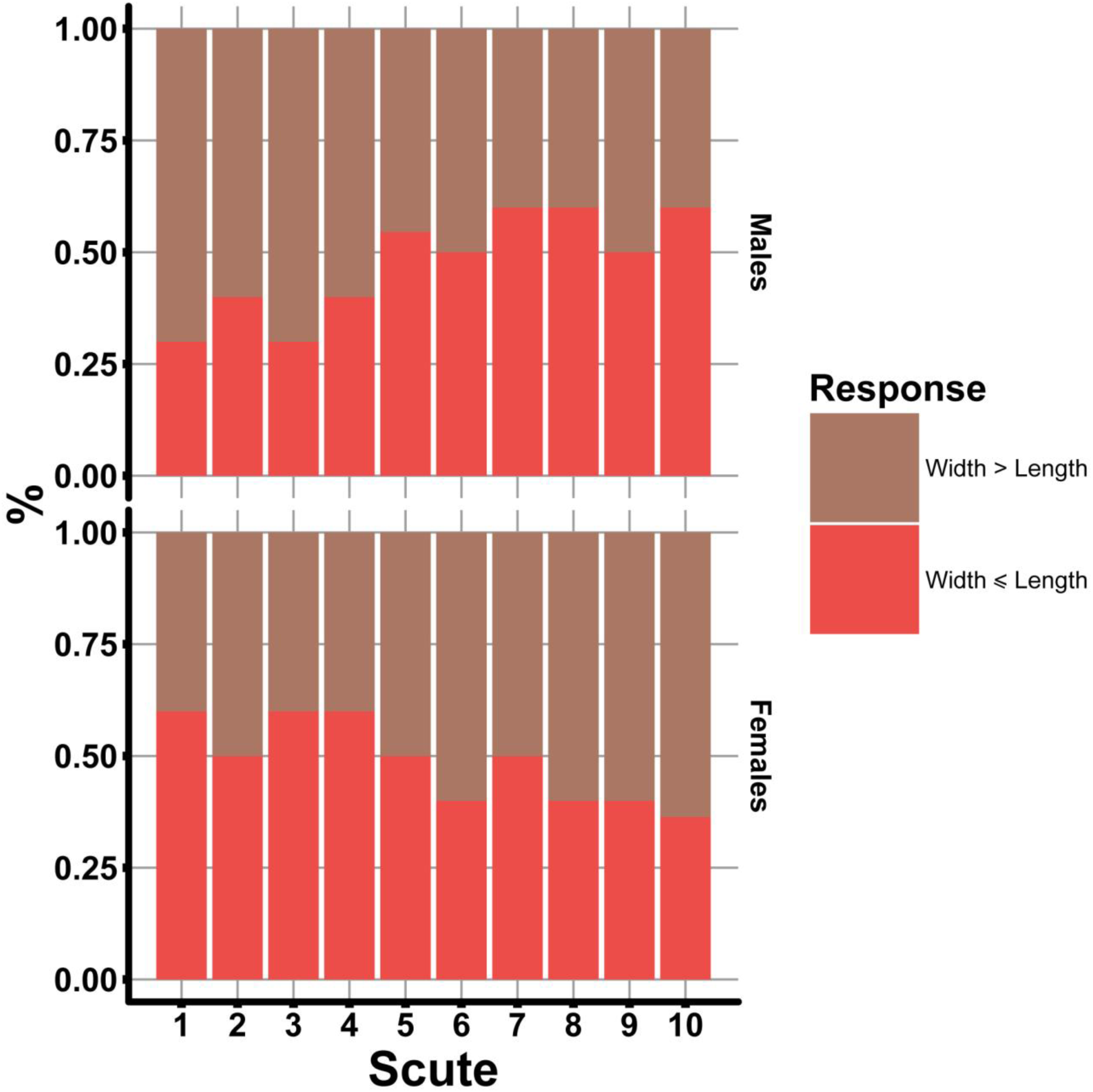
Plot show result of chi-square (χ^2^) test for visual perception of relationship between width (W) and length (L) of dorsal scutes of juveniles *Acipenser ruthenus*. Number of fish treated = 20.

**Fig. 34.**
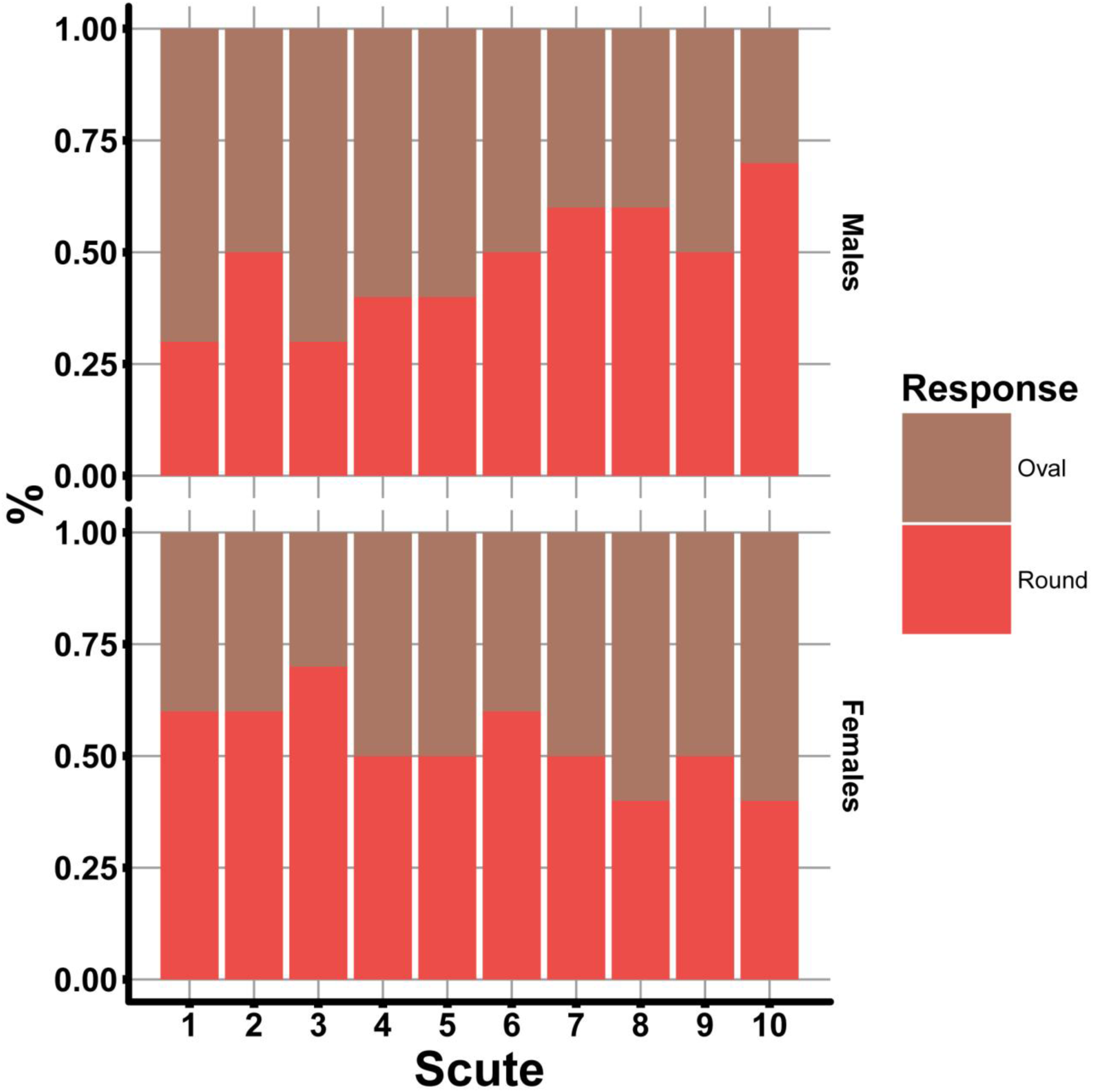
Plot show result of chi-square (χ^2^) test for visual perception of dorsal scute shape of juveniles *Acipenser ruthenus*. Number of fish treated = 20.

**Fig. 35.**
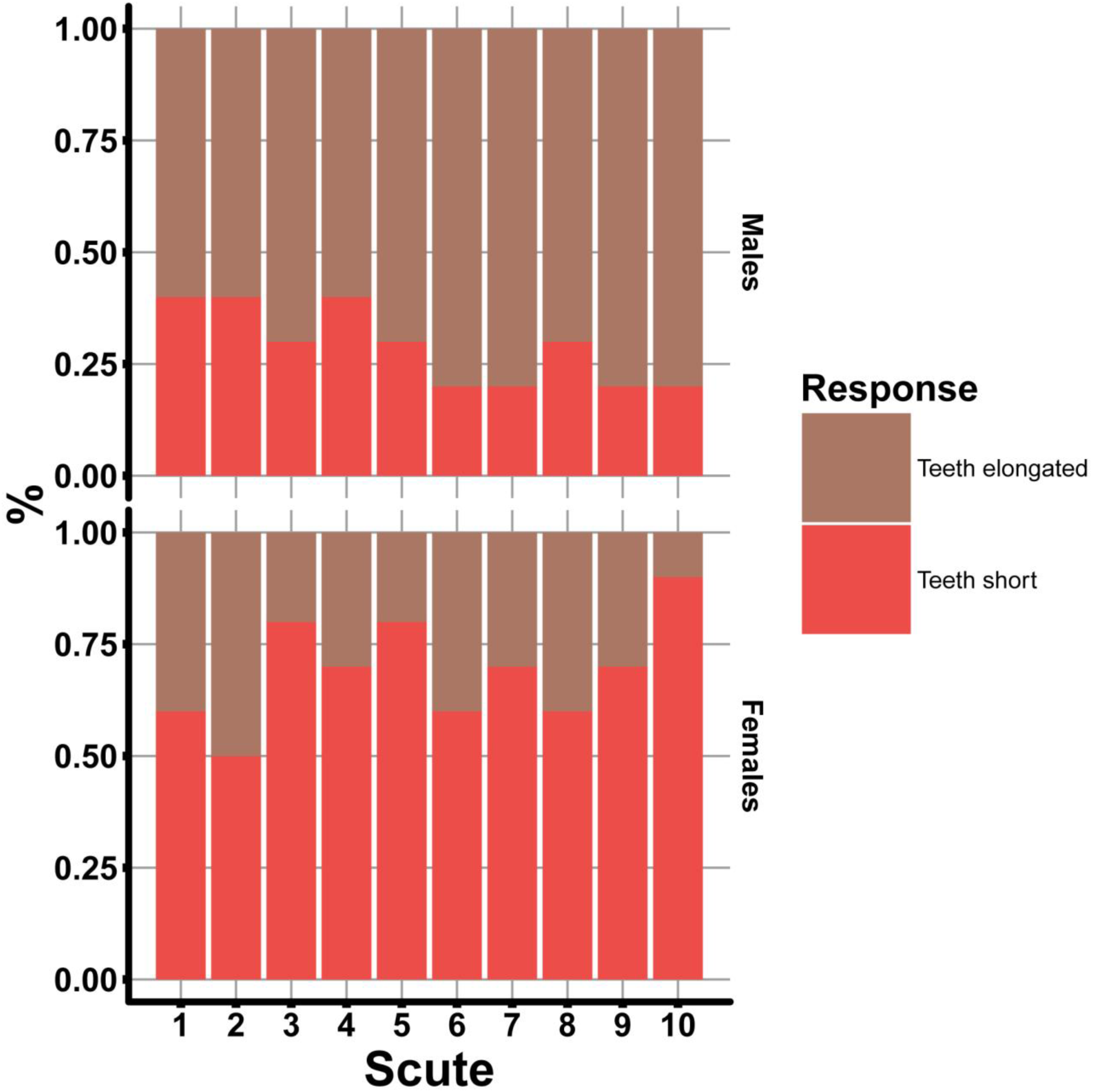
Plot show result of chi-square (χ^2^) test for visual perception of length of the dorsal scute teeth of juveniles *Acipenser ruthenus*. Number of fish treated = 20.

**Fig. 36.**
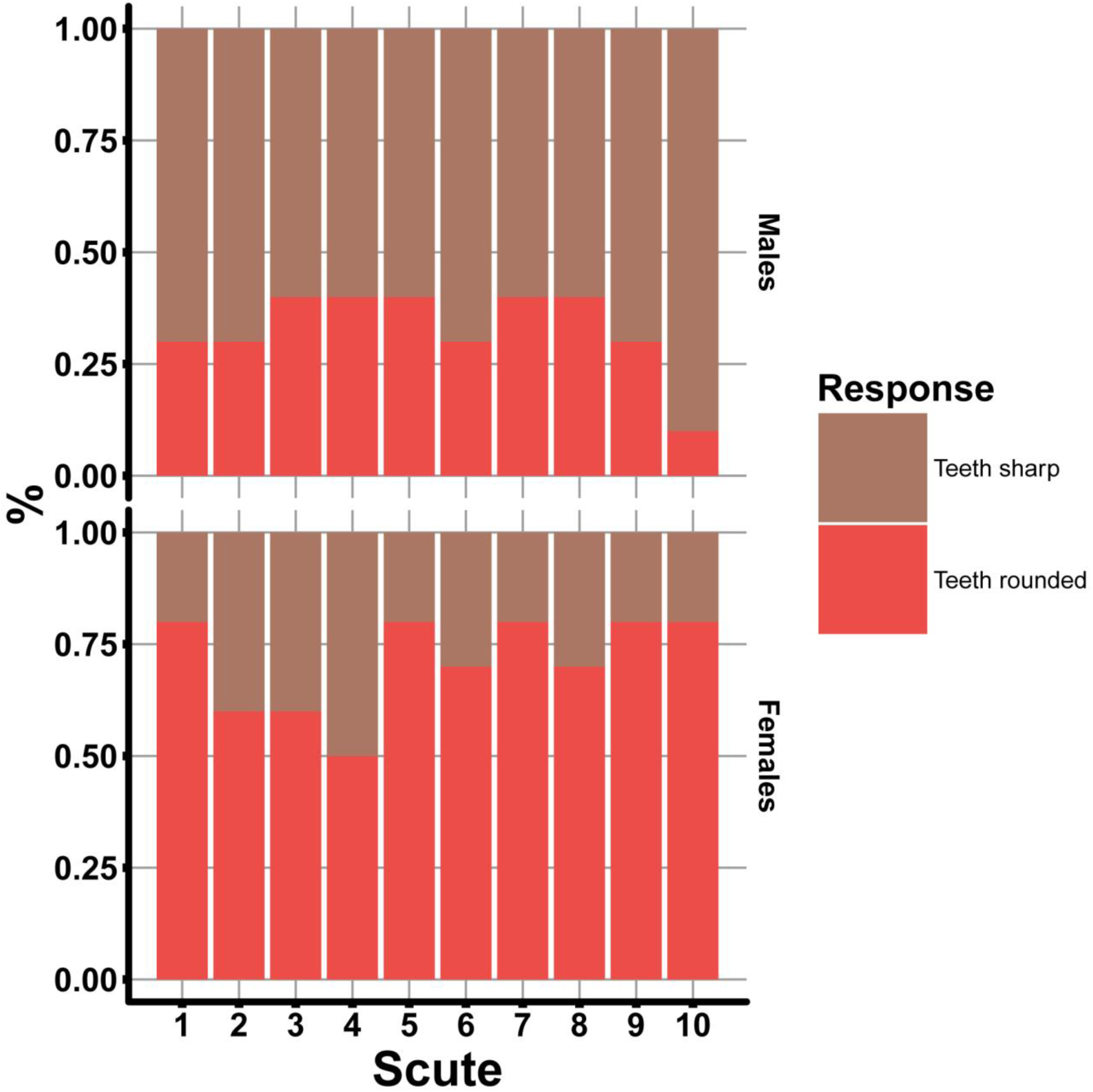
Plot show result of chi-square (χ^2^) test for visual perception of shape of the dorsal scute teeth of adults *Acipenser ruthenus*. Number of fish treated = 20.

**Fig. 37.**
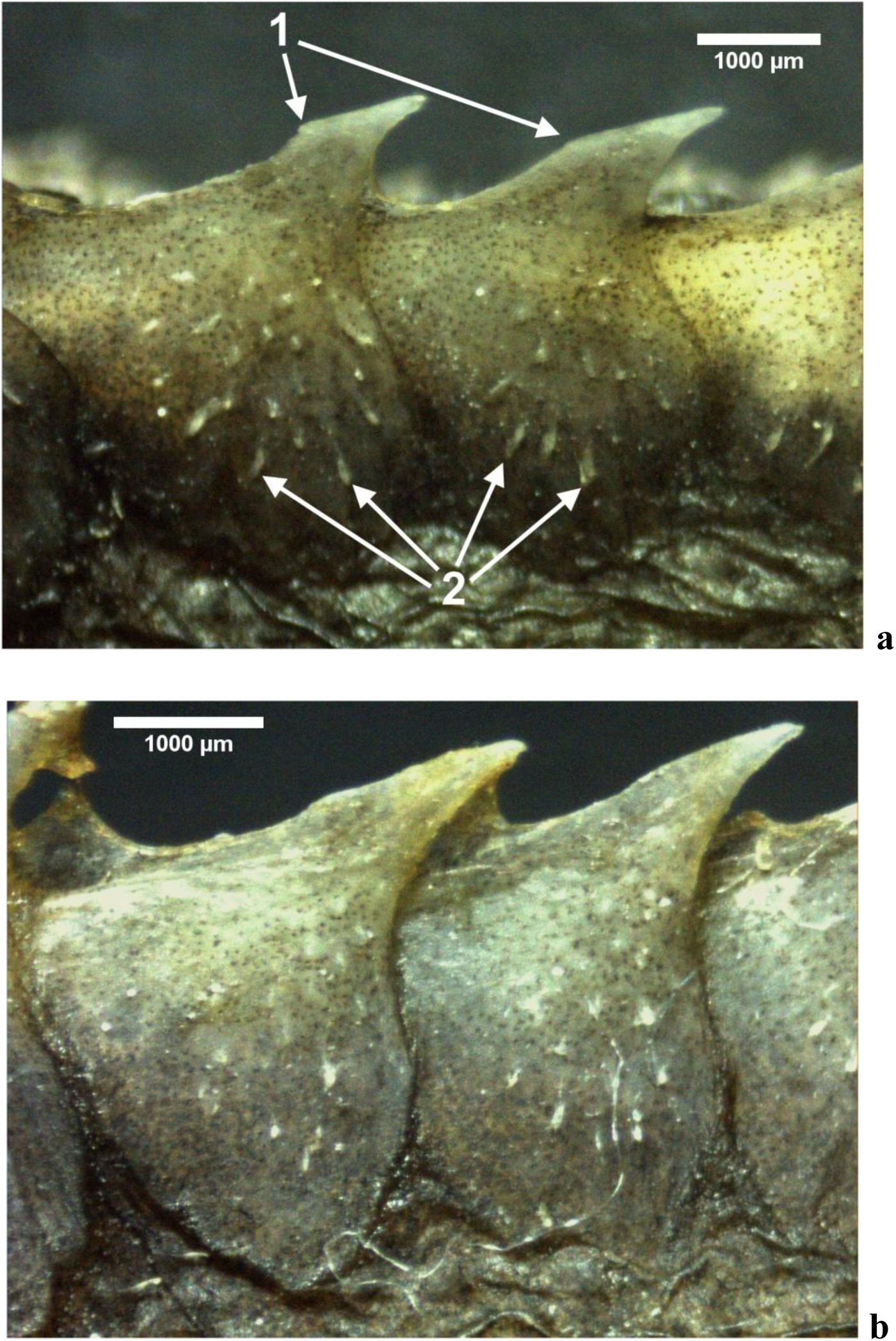
Juvenile scutes of future males (a) have significantly different than scutes of future females (b). The arrows show small and sharp teeth on the scute surface (1) and the curvature at the top (2).

**Table 2.**
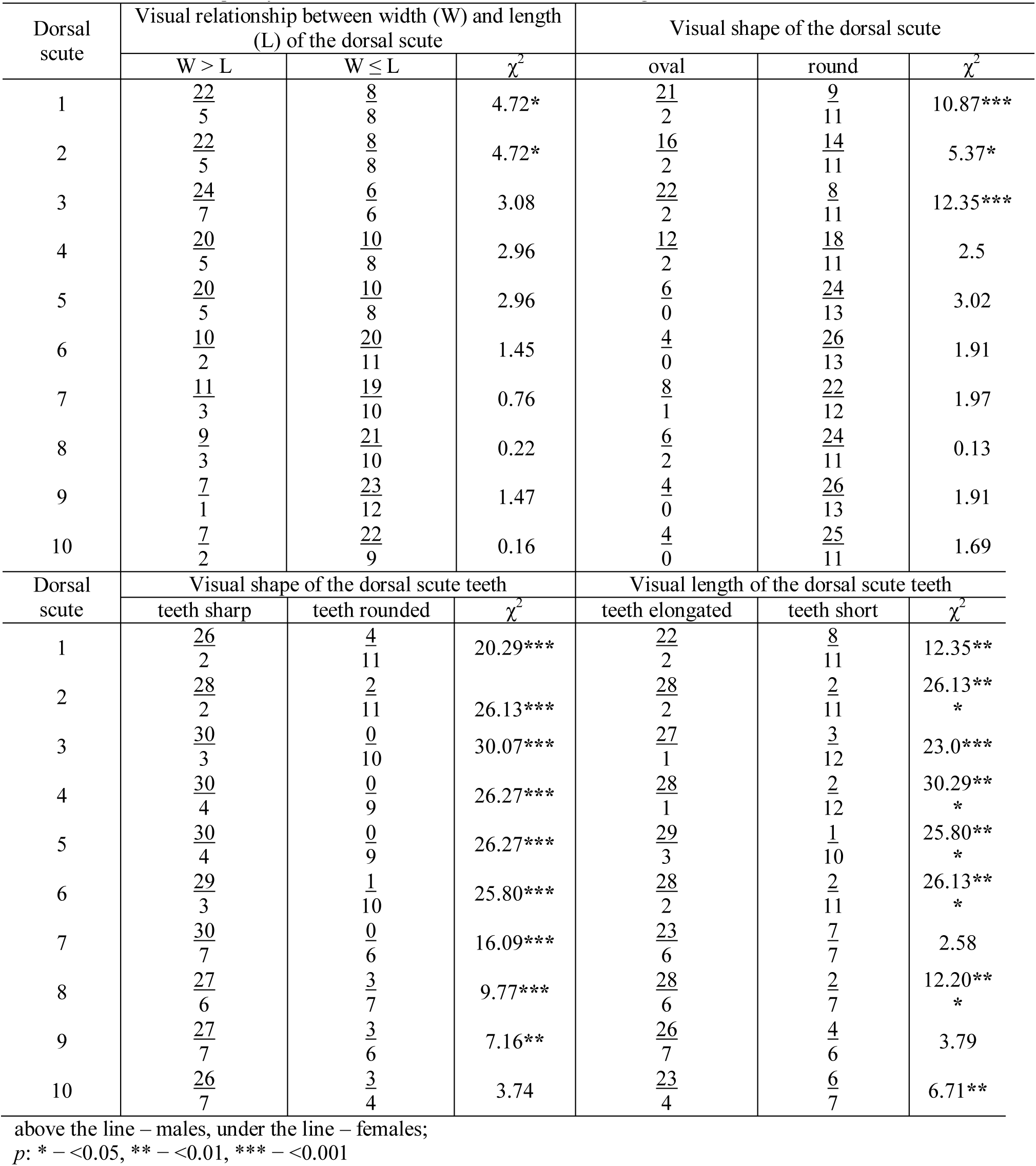
Occurrence frequency of various scute forms of adult sterlet sturgeons

**Table 3.** Occurrence frequency of various scute forms of young sterlet sturgeons

| Dorsal scute | Visual relationship between width (W) and length (L) of the dorsal scute |  |  | Visual shape of the dorsal scute |  |  |
| --- | --- | --- | --- | --- | --- | --- |
| | W > L | W ≤ L | $\chi^2$ | oval | round | $\chi^2$ |
| 1 | $\frac{8}{6}$ | $\frac{2}{4}$ | 0.95 | $\frac{8}{2}$ | $\frac{2}{8}$ | 7.20** |
| 2 | $\frac{8}{5}$ | $\frac{2}{5}$ | 1.98 | $\frac{8}{2}$ | $\frac{2}{8}$ | 7.20** |
| 3 | $\frac{8}{4}$ | $\frac{2}{6}$ | 3.33 | $\frac{8}{2}$ | $\frac{2}{8}$ | 7.20** |
| 4 | $\frac{8}{3}$ | $\frac{2}{7}$ | 5.05* | $\frac{8}{2}$ | $\frac{2}{8}$ | 7.20** |
| 5 | $\frac{7}{2}$ | $\frac{3}{8}$ | 5.05* | $\frac{6}{2}$ | $\frac{4}{8}$ | 3.33 |
| 6 | $\frac{6}{2}$ | $\frac{4}{8}$ | 3.33 | $\frac{5}{2}$ | $\frac{5}{8}$ | 1.98 |
| 7 | $\frac{7}{2}$ | $\frac{3}{8}$ | 5.05* | $\frac{8}{2}$ | $\frac{2}{8}$ | 7.20** |
| 8 | $\frac{7}{4}$ | $\frac{3}{6}$ | 1.82 | $\frac{6}{2}$ | $\frac{4}{8}$ | 3.33 |
| 9 | $\frac{6}{5}$ | $\frac{4}{5}$ | 0.65 | $\frac{5}{2}$ | $\frac{5}{8}$ | 1.98 |
| 10 | $\frac{5}{5}$ | $\frac{5}{5}$ | 0.00 | $\frac{5}{2}$ | $\frac{5}{8}$ | 1.98 |
| Dorsal scute | Visual shape of the dorsal scute teeth |  |  | Visual length of the dorsal scute teeth |  |  |
| | teeth sharp | teeth rounded | $\chi^2$ | teeth elongated | teeth short | $\chi^2$ |
| 1 | $\frac{9}{1}$ | $\frac{1}{9}$ | 12.80*** | $\frac{7}{1}$ | $\frac{3}{9}$ | 7.50** |
| 2 | $\frac{9}{1}$ | $\frac{1}{9}$ | 12.80*** | $\frac{8}{1}$ | $\frac{2}{9}$ | 9.90** |
| 3 | $\frac{9}{1}$ | $\frac{1}{9}$ | 12.80*** | $\frac{7}{2}$ | $\frac{3}{8}$ | 5.05* |
| 4 | $\frac{9}{1}$ | $\frac{1}{9}$ | 12.80*** | $\frac{6}{2}$ | $\frac{4}{8}$ | 3.33 |
| 5 | $\frac{8}{2}$ | $\frac{2}{8}$ | 7.20** | $\frac{6}{1}$ | $\frac{4}{9}$ | 5.49* |
| 6 | $\frac{9}{2}$ | $\frac{1}{9}$ | 10.83*** | $\frac{9}{2}$ | $\frac{1}{8}$ | 9.90** |
| 7 | $\frac{9}{1}$ | $\frac{1}{9}$ | 12.80*** | $\frac{9}{1}$ | $\frac{1}{9}$ | 12.80*** |
| 8 | $\frac{7}{2}$ | $\frac{3}{8}$ | 5.05* | $\frac{8}{4}$ | $\frac{2}{6}$ | 3.33 |
| 9 | $\frac{6}{3}$ | $\frac{4}{7}$ | 1.82 | $\frac{6}{3}$ | $\frac{4}{7}$ | 1.82 |
| 10 | $\frac{6}{3}$ | $\frac{4}{7}$ | 1.82 | $\frac{6}{3}$ | $\frac{4}{7}$ | 1.82 |

above the line – males, under the line – females;
p: \* – <0.05, \*\* – <0.01, \*\*\* – <0.001.

**Table 4.**
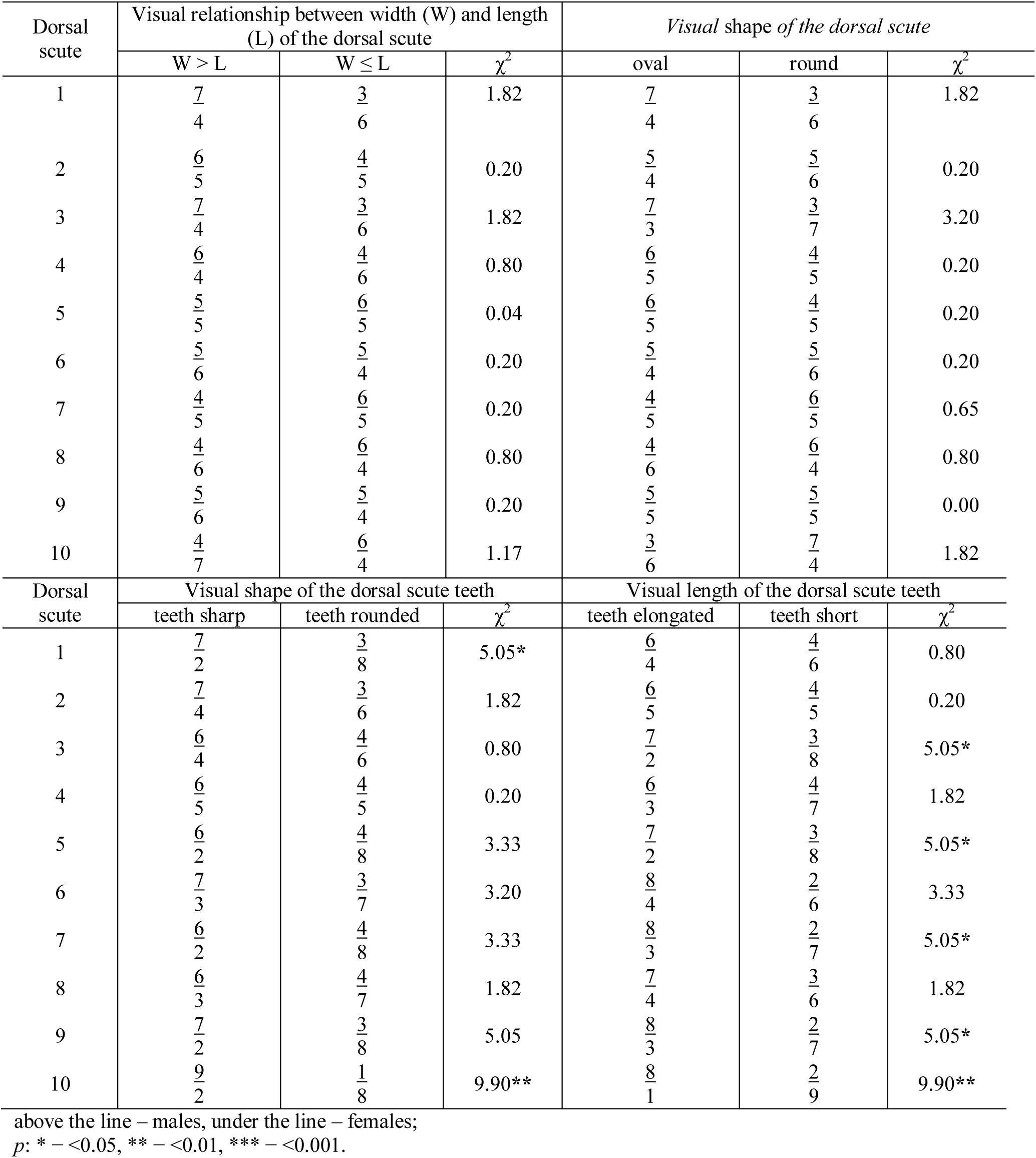
Occurrence frequency of various scute forms of juvenile sterlet sturgeons

Thus, we found, that male scutes had a broader and oblate (oval) shape; conditional lines drawn through the scute endpoints formed an isosceles triangle; the scute blade were separated from the scute at a less acute angle; the scutes teeth were more pointed and elongated relative to the scute width; the scutes teeth number was more than in females; the scutes surface had many acute small teeth. Female scutes had a more rounded shape, conditional lines drawn through the scute endpoints formed an equilateral triangle, the scute blade separated from the scute at a more obtuse angle, the scutes teeth were more rounded and small lengthened relative to the scute width, the scutes surface had more smooth (Fig. 38, 39, 40, 41, 42, 43).

**Fig. 38.**
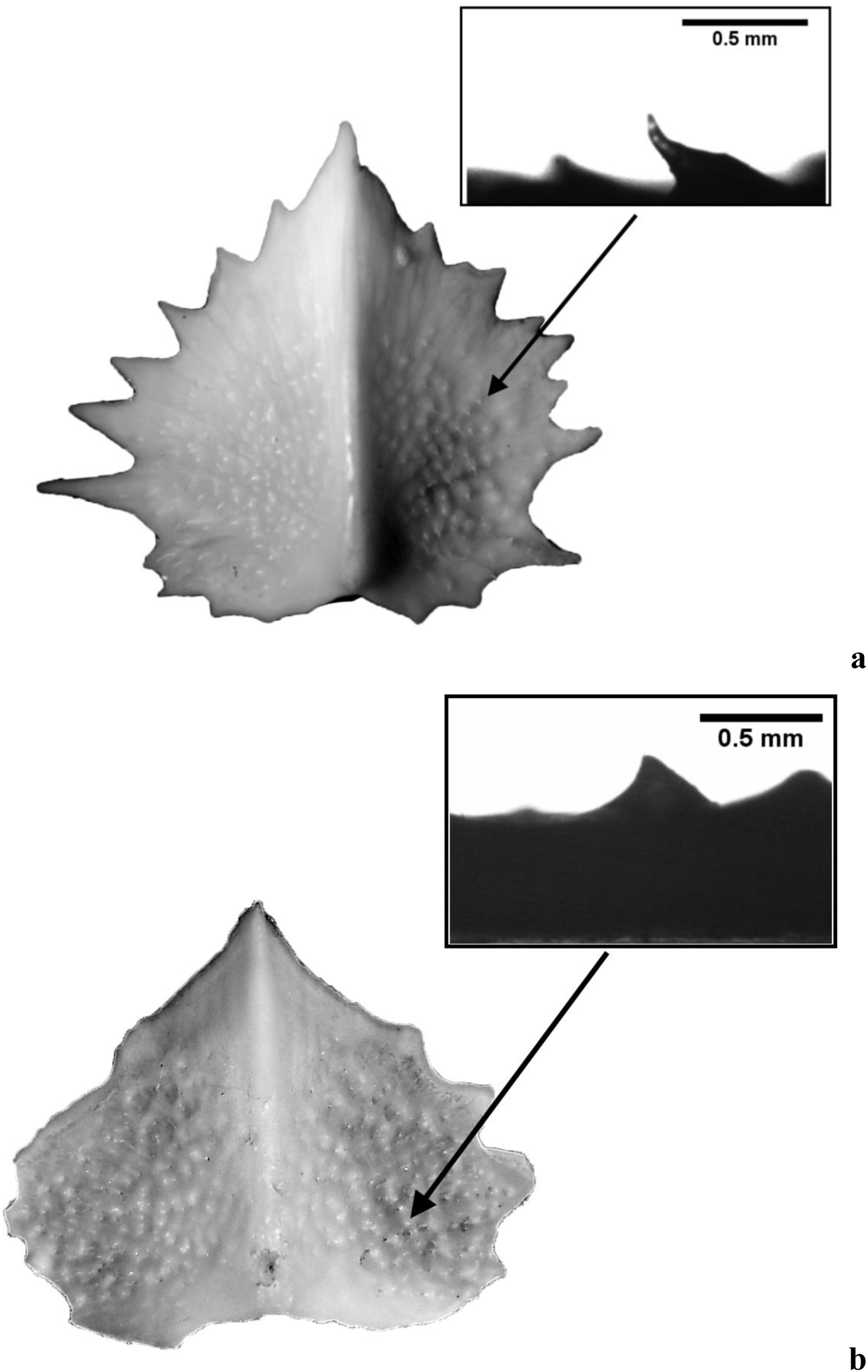
Typical surface of male (a) and female (b) dorsal scute of adult *Acipenser ruthenus*

**Fig. 39.**
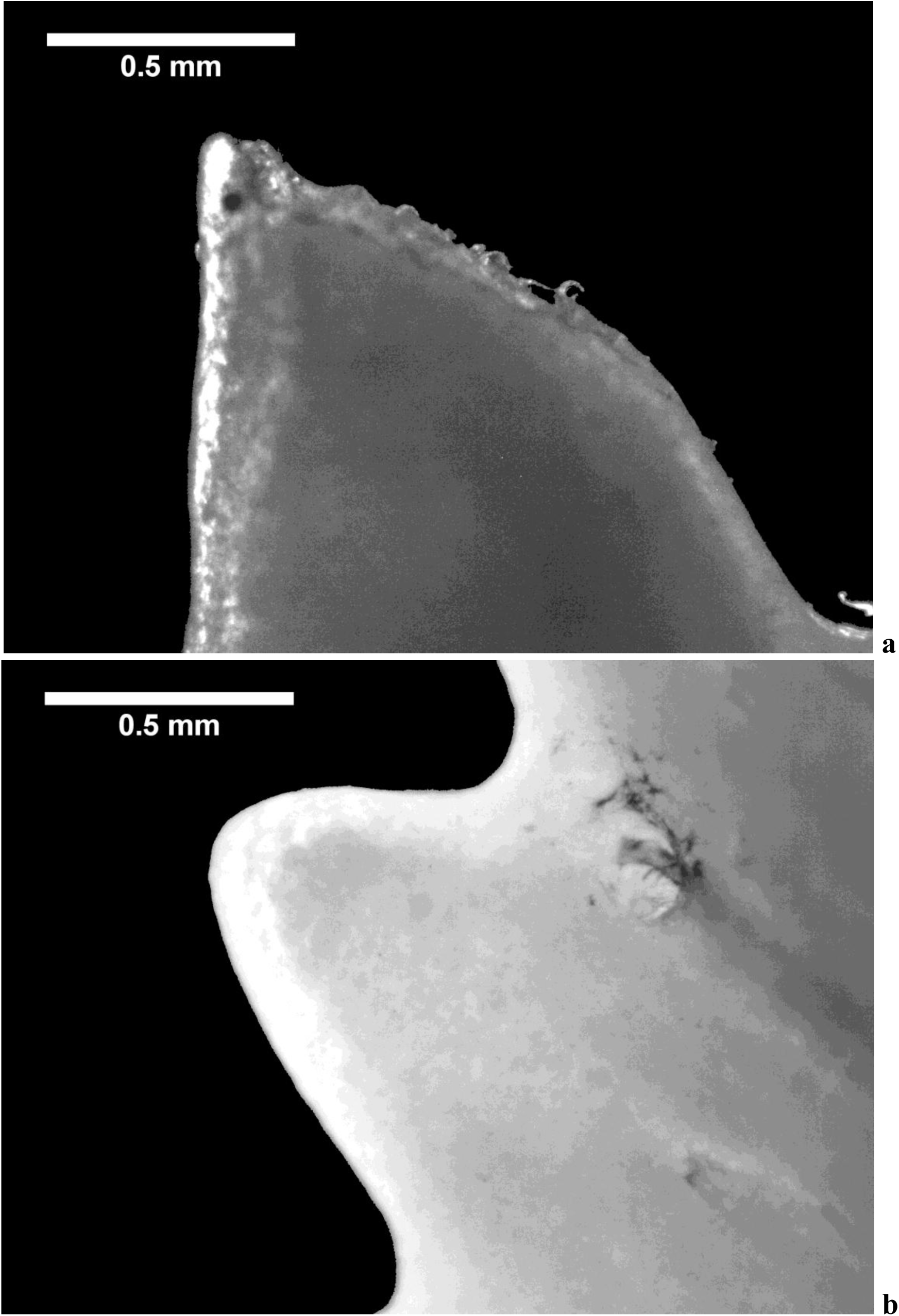
Typical edge of male (a) and female (b) scute teeth of adult *Acipenser ruthenus*

**Fig. 40.**
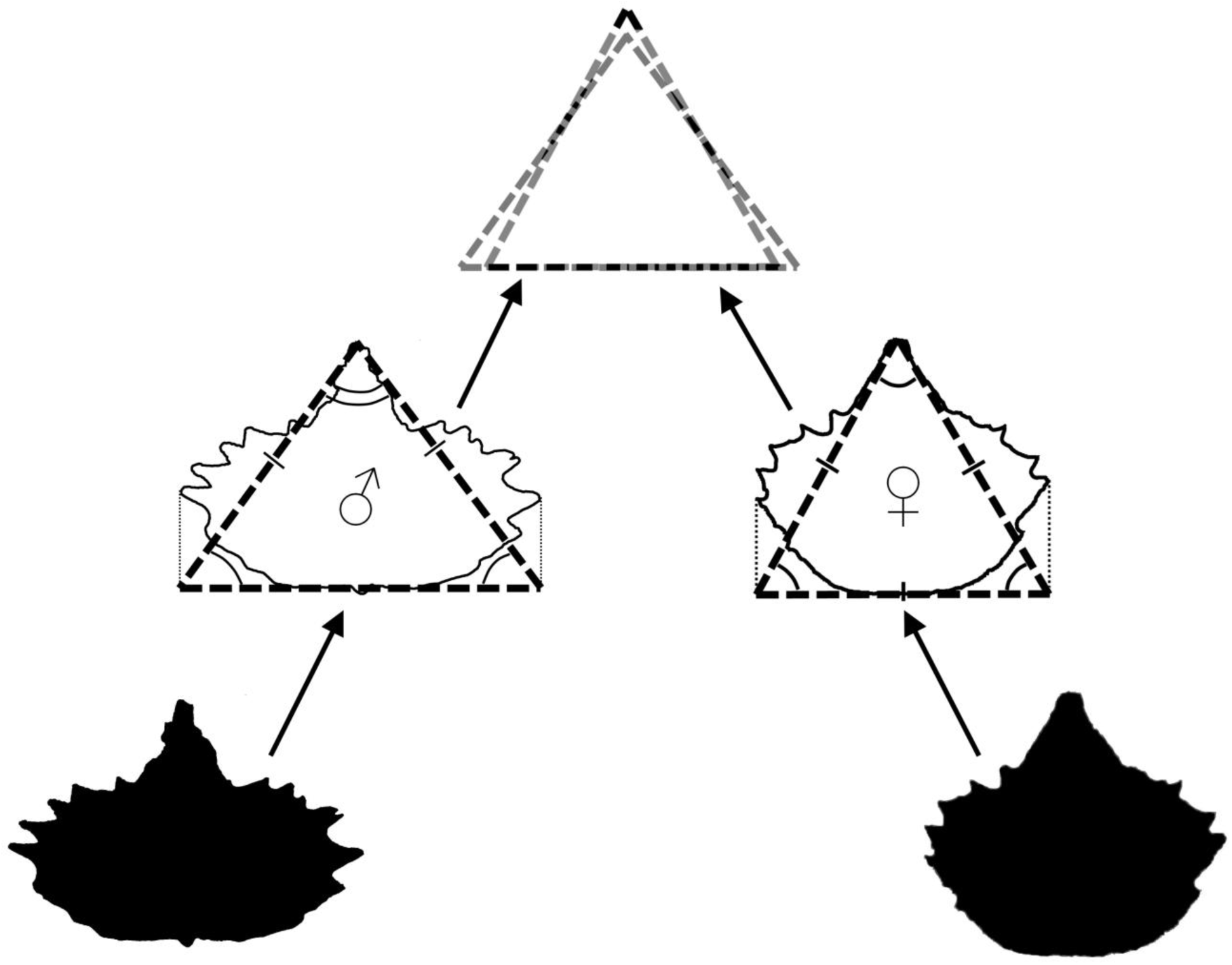
Typical male scutes of *Acipenser ruthenus* had a broader and oblate (oval) shape; conditional lines drawn through the scute endpoints formed an isosceles triangle. Female scutes of *Acipenser ruthenus* had a more rounded shape, conditional lines drawn through the scute endpoints formed an equilateral triangle.

**Fig. 41.**
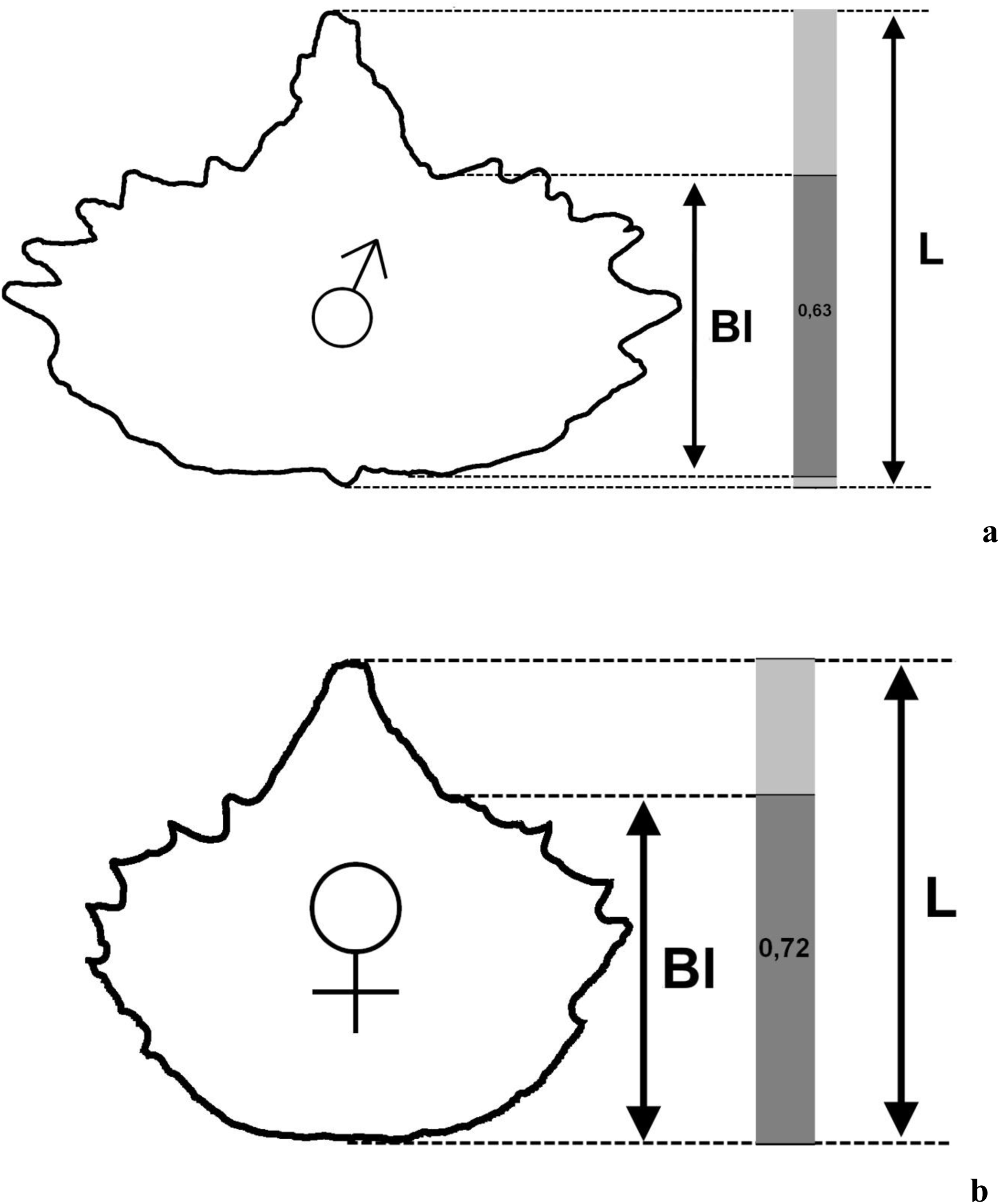
Impact of changing the blade index on visual perception of different dorsal scute form between male (a) and female (b) of *Acipenser ruthenus*

**Fig. 42.**
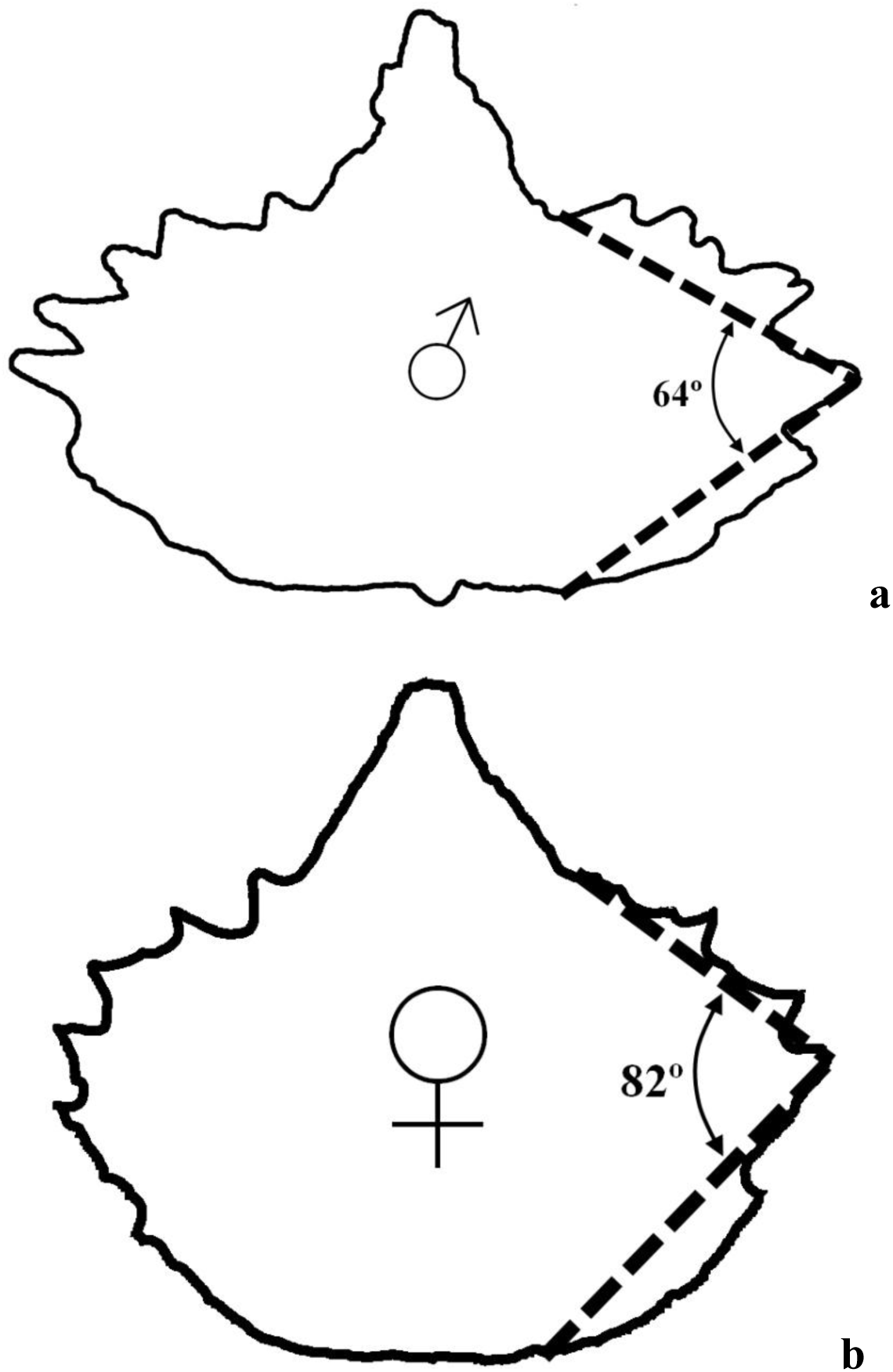
Changing the blade angle of dorsal scute between male (a) and female (b) of *Acipenser ruthenus*

**Fig. 43.**
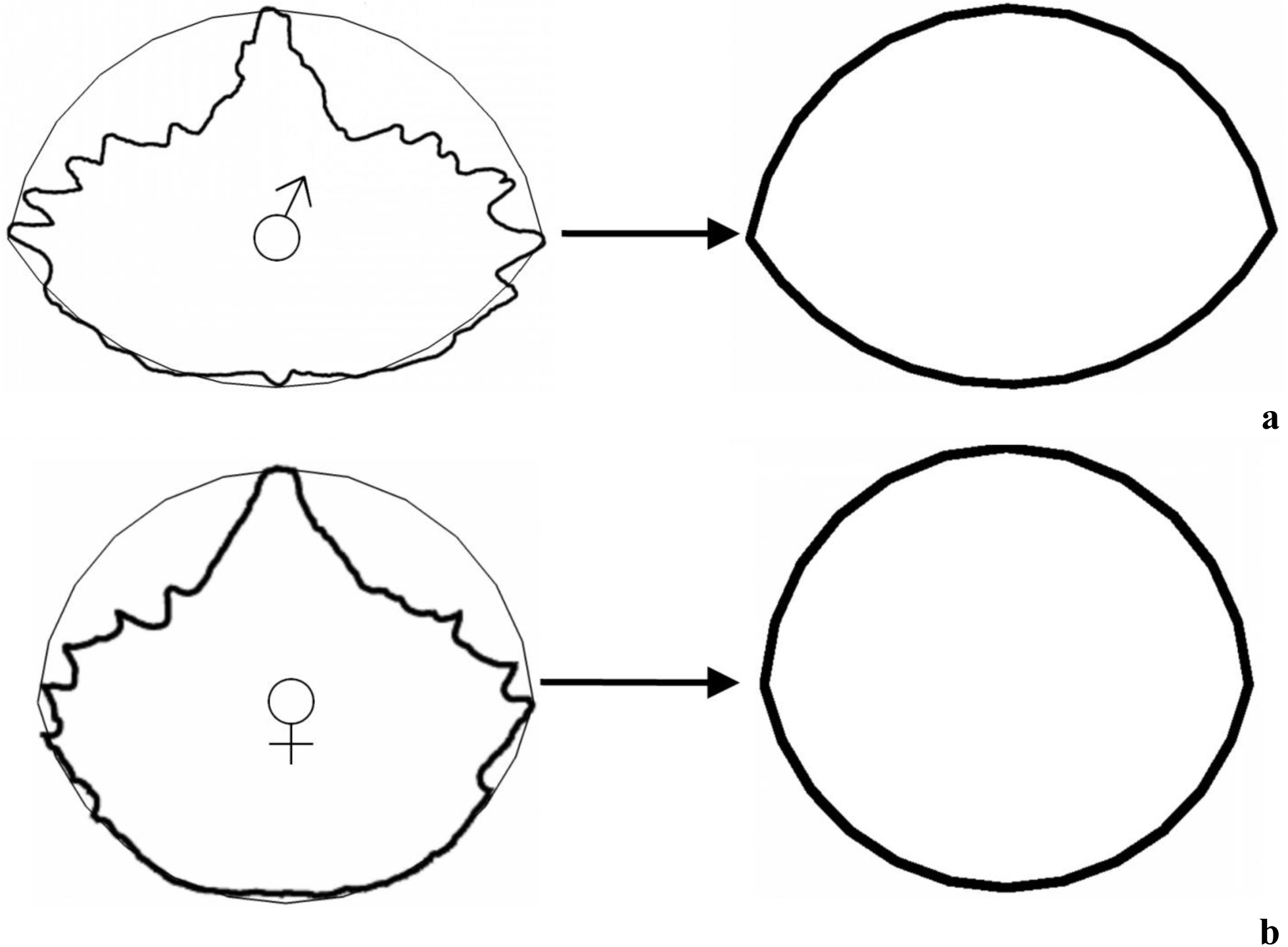
Impact of changing the fill index on visual perception of dorsal scute form between male (a) and female (b) of *Acipenser ruthenus*

We first found that the distance between the dorsal scutes in adult sterlet depended on sex. This dependency was manifested only in adults. The spacing between the first five dorsal scutes was wider in females than in males. (Fig. 44, 45).

**Fig. 44.**
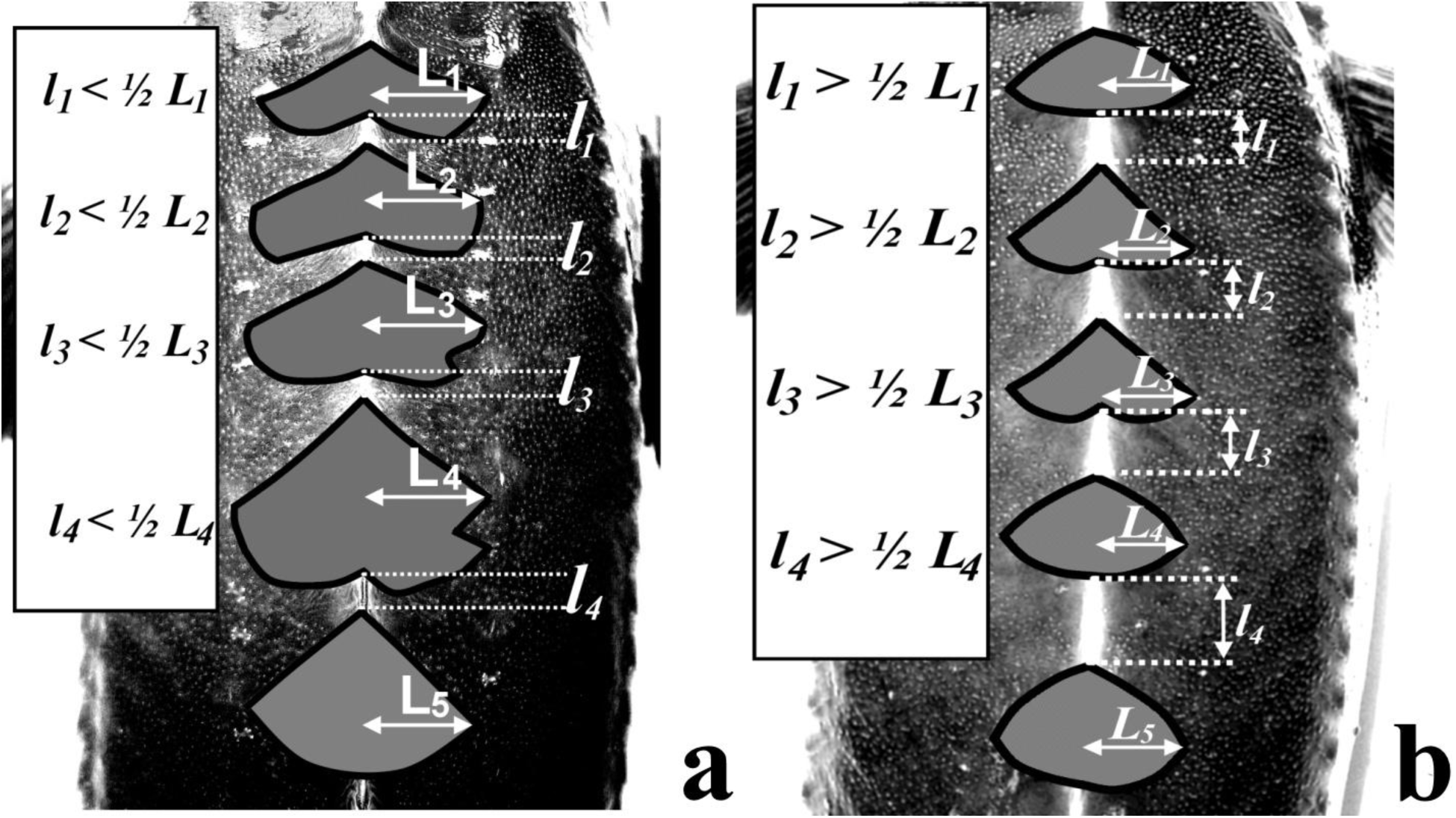
Example of narrow scute distance. Criterion for this parameter is the average distance (*l*) between the first to fifth dorsal scutes, which should be less than ½ the average blade length (*L*) of dorsal scute (a)). This criterion is determined only in adult fishes.

**Fig. 45.**
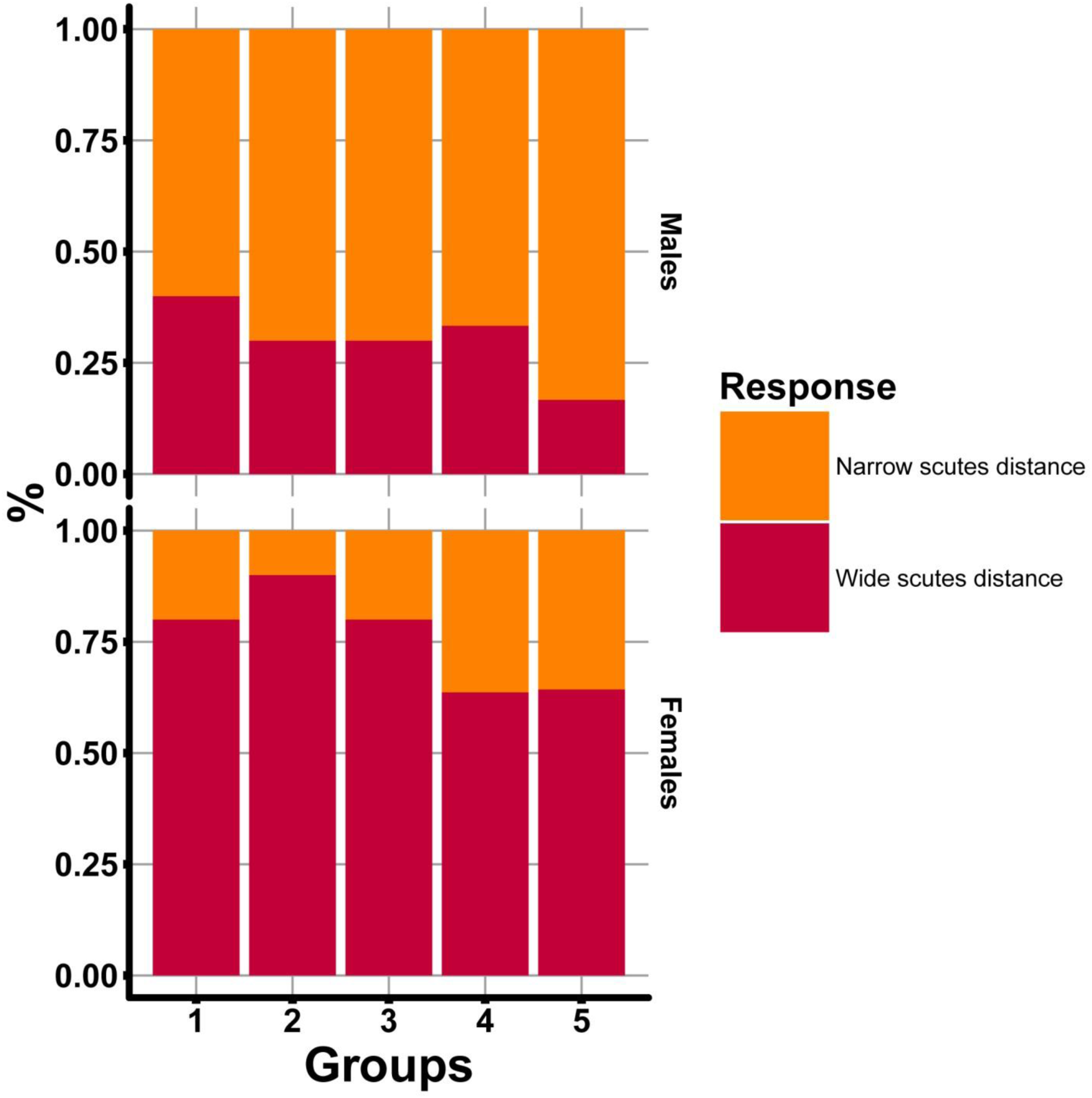
Plot show result of chi-square (χ^2^) test for visual perception of narrow scute distance between the first to fifth dorsal scutes of adults *Acipenser ruthenus*. Number of fish treated = 20.

We developed the sex determination method for sterlet using a binary matrix and binary discriminant analysis with DA algorithm based on the selection results of quantitative parameters using machine learning method (neural networks (Fritsch, Guenther, 2016), random forrest method (Paluszynska, Biecek, 2017), boruta algorithm (Kursa, Rudnicki, 2010)) and chi-square test. The results obtained with the help of tree derivation method on the basis of recursive decomposition were used as the criterion of transition from 1 to 0 in the binary matrix (Fig. 46, 47, 48). The error of model was 7 % for adults, 4 % for young and 4 % for juveniles sterlet.

**Fig. 46.**
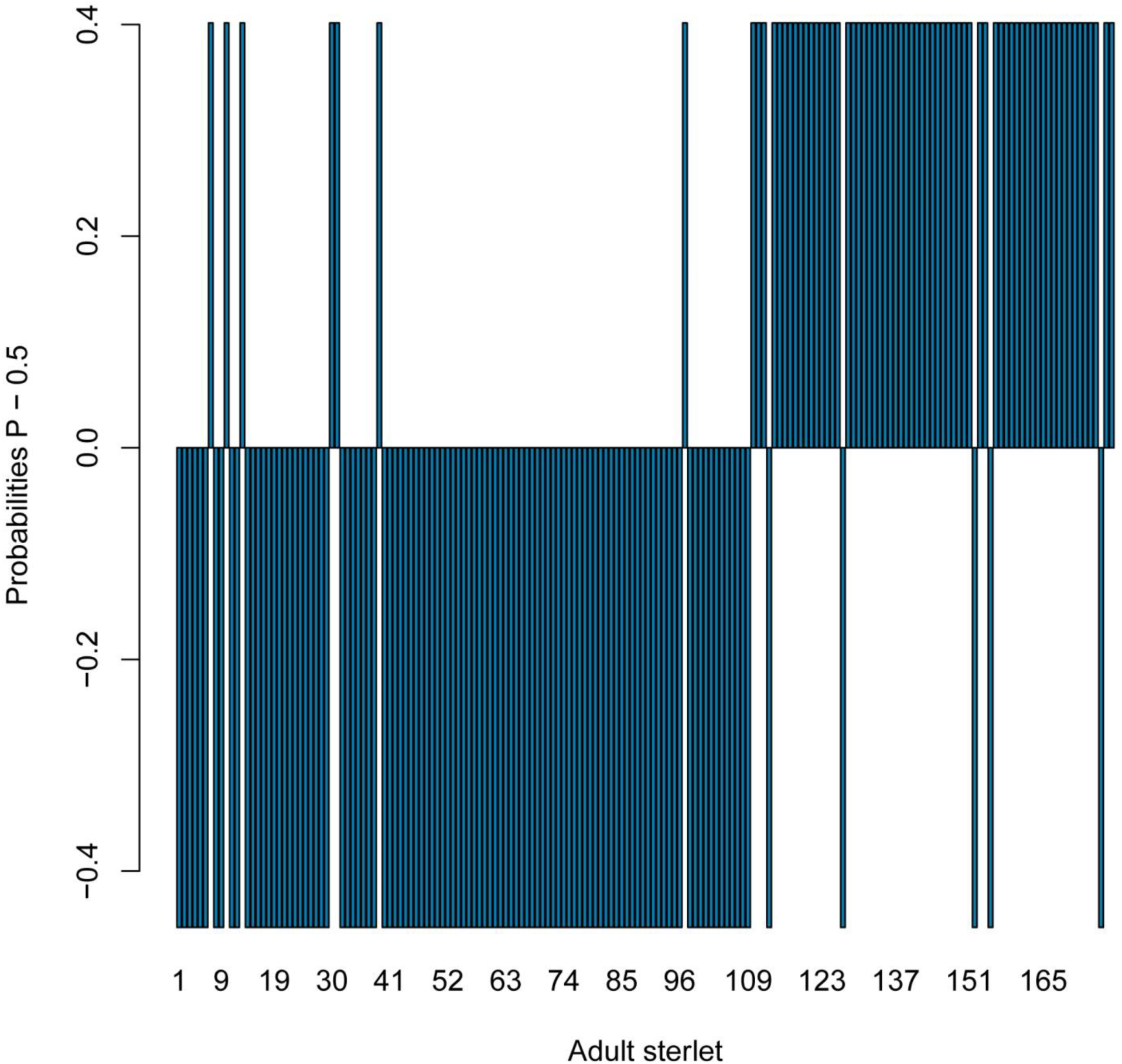
Estimating sex predicted probabilities from logistic regression for adults *Acipenser ruthenus* using decision tree model. Error of model – 7 %.

**Fig. 47.**
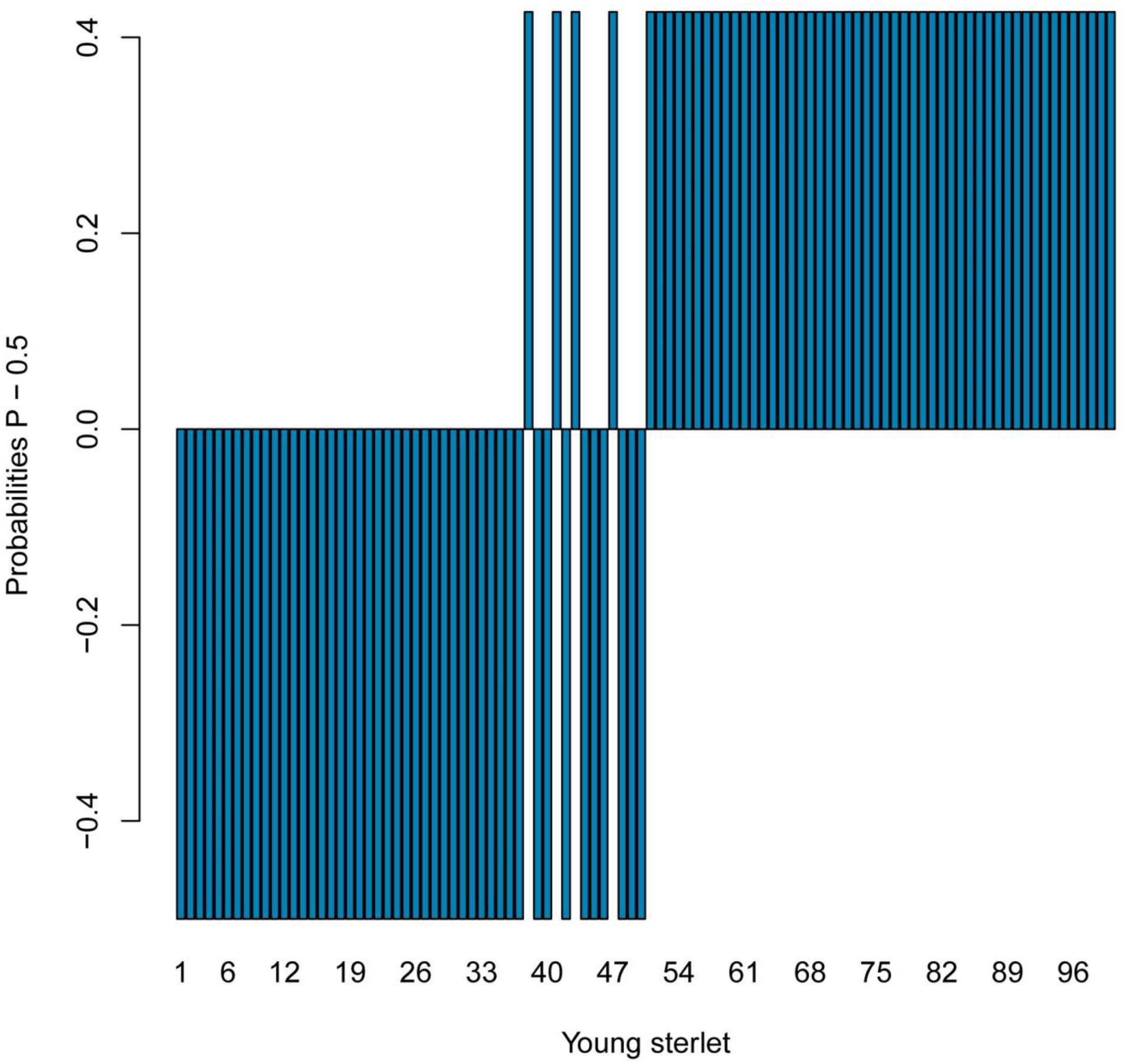
Estimating sex predicted probabilities from logistic regression for young *Acipenser ruthenus* using decision tree model. Error of model – 4 %.

**Fig. 48.**
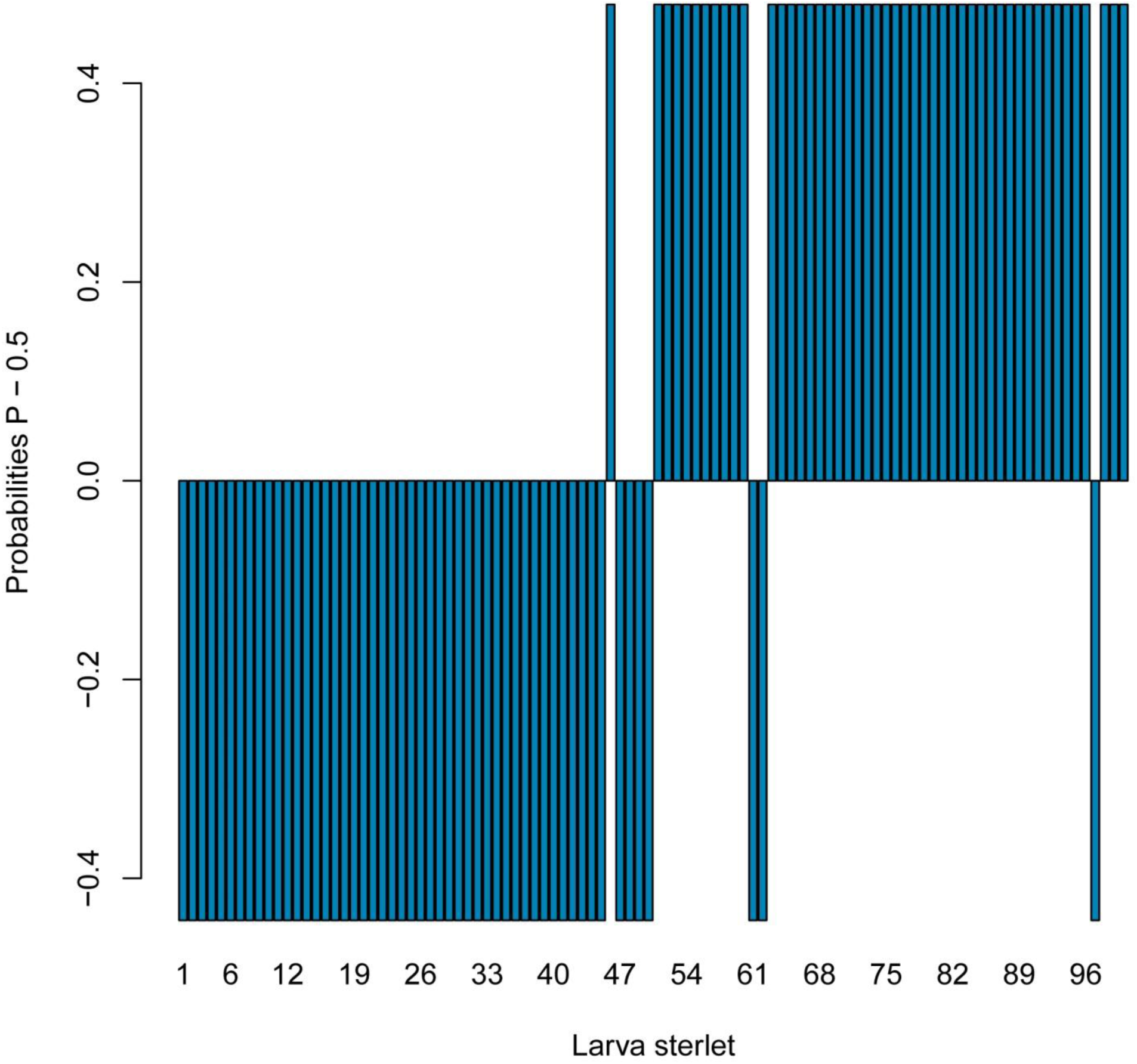
Estimating sex predicted probabilities from logistic regression for juveniles *Acipenser ruthenus* using decision tree model. Error of model – 4 %.

Also, we developed an easy-to-use in aquaculture sex determination system (Tabl. 5). Tried and tested, this system showed that the average points for males were 5.6 ± 0.1 (min = 5, max = 10), for females – 1.3 ± 0.2 (min = 0, max = 4). The observed differences were statistically significant (p <0.05).

**Table 5.** Sterlet sex determination system for easy-to-use in aquaculture

| Criterion | Variant of the criterion | Point |
| --- | --- | --- |
| Visual shape of the dorsal scute | Dorsal scute oblate, oval | 1 |
|  | Dorsal scute elongated, rounded | 0 |
| Visual shape of the dorsal scute teeth | Dorsal scute teeth sharp and elongated | 1 |
|  | Dorsal scute teeth short, rounded or no identified | 0 |
| Narrow scute distance | Narrow scute distance | 1 |
|  | Wide scute distance | 0 |

We have developed a protocol for sex determination of sterlet on the basis of our research (Supplementary Materials).

We studied only dorsal scutes. Study of other the sterlet scutes rows requires additional research. Preliminary results show that the lateral and abdominal starlet scutes also have sex-related structural differences.

We have not found that dorsal scutes of Siberian sturgeon have significant statistical differences between males and females (Fig. 49), but we found that lateral and abdominal scutes of Siberian sturgeon have significant statistical differences between males and females. Significant statistical differences remained in adults (2 years-old, length – 73.2±2.7 cm) and also in young (1 year-old, 34.8±1.9 cm) and juveniles (3 months-old, 11.5±4.2 cm) – future mature males and females.

**Fig. 49.**
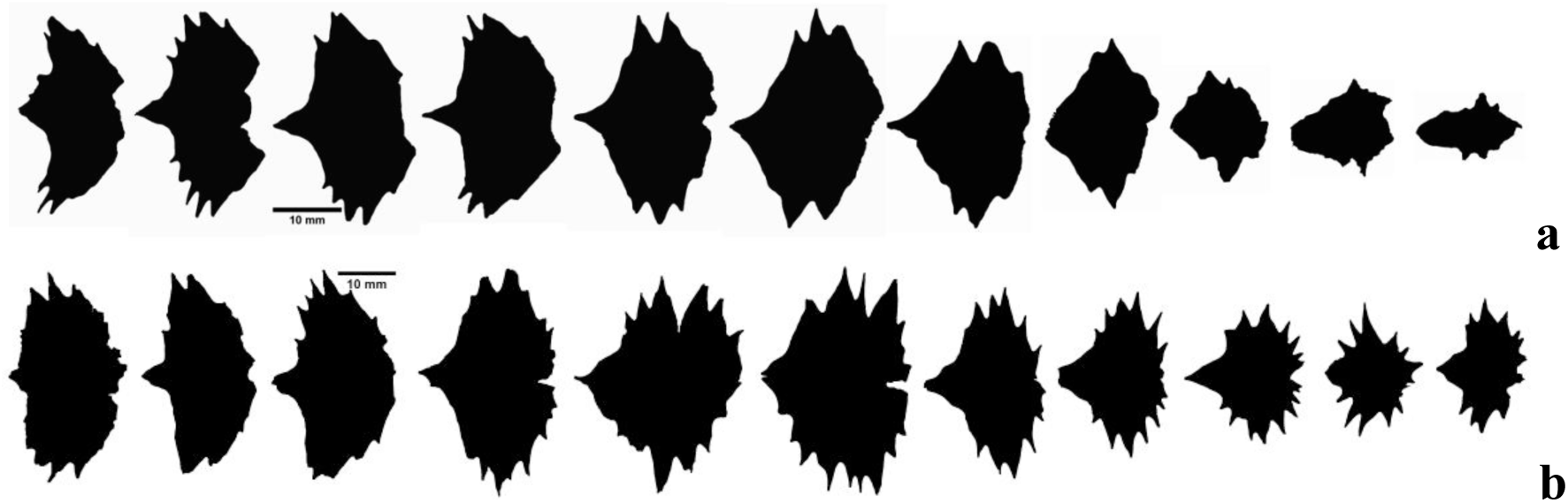
Contour samples of the typical dorsal scutes of adults *Acipenser baerii*. a – the dorsal scutes of males; b – the dorsal scutes of females. For creating the scute contours was using binary process of ImageJ software.

Male lateral scutes of Siberian sturgeon had a narrow and oblate shape; the scutes teeth were more pointed and elongated relative to the scute height; the female lateral scutes had a more rounded shape, the scutes teeth were more rounded and a little lengthened relative to the scute height (Fig. 50).

**Fig. 50.**
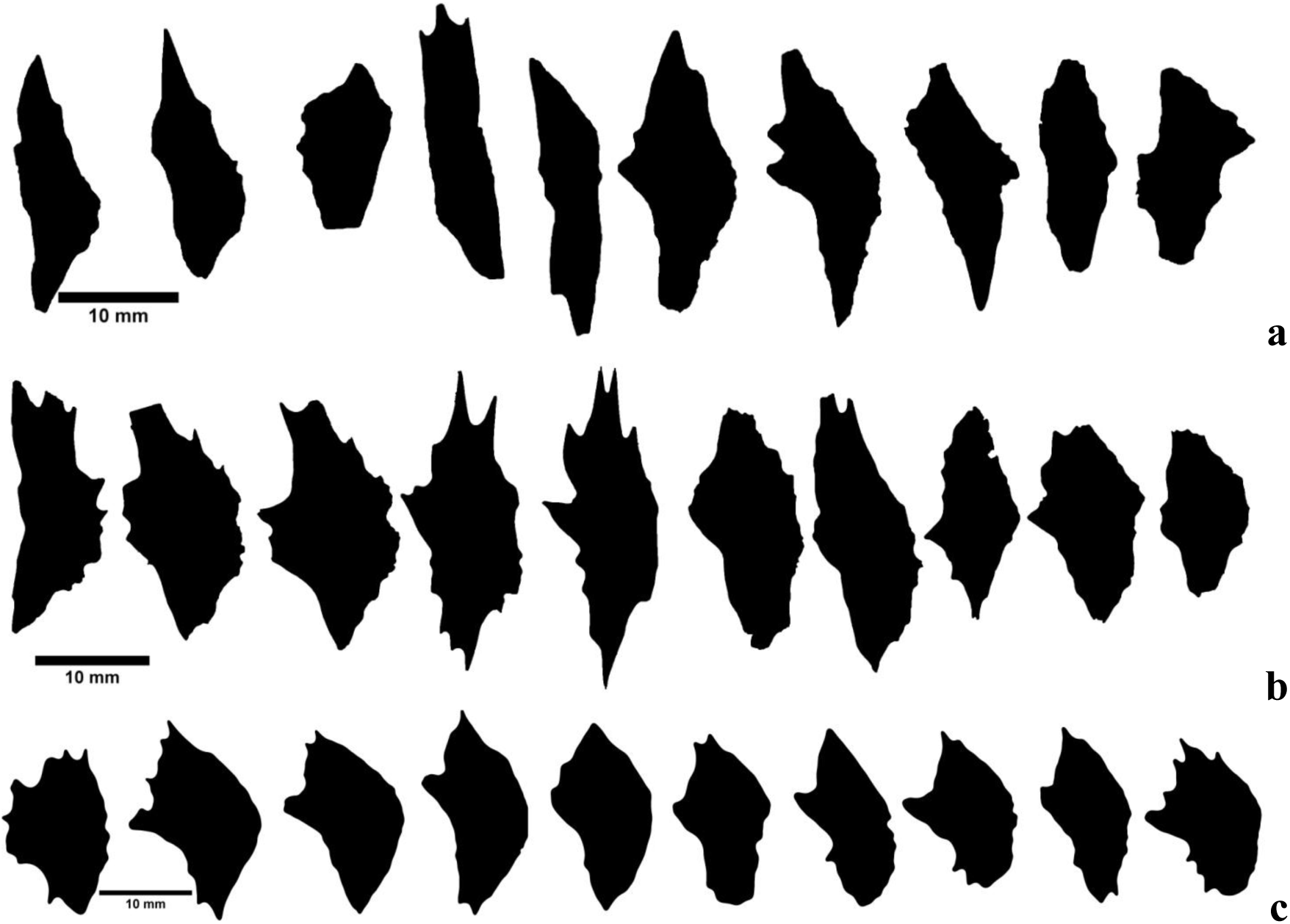
Contour samples of the typical lateral scutes of adult *Acipenser baerii*. a, b – the lateral scutes of male; c – the lateral scutes of female. For creating the scute contours was using binary process of ImageJ software.

Lateral and abdominal juvenile scutes of future mature males of Siberian sturgeon had a large number of pointed and elongated scutes teeth (Fig. 51, 52, 53, 54).

**Fig. 51.**
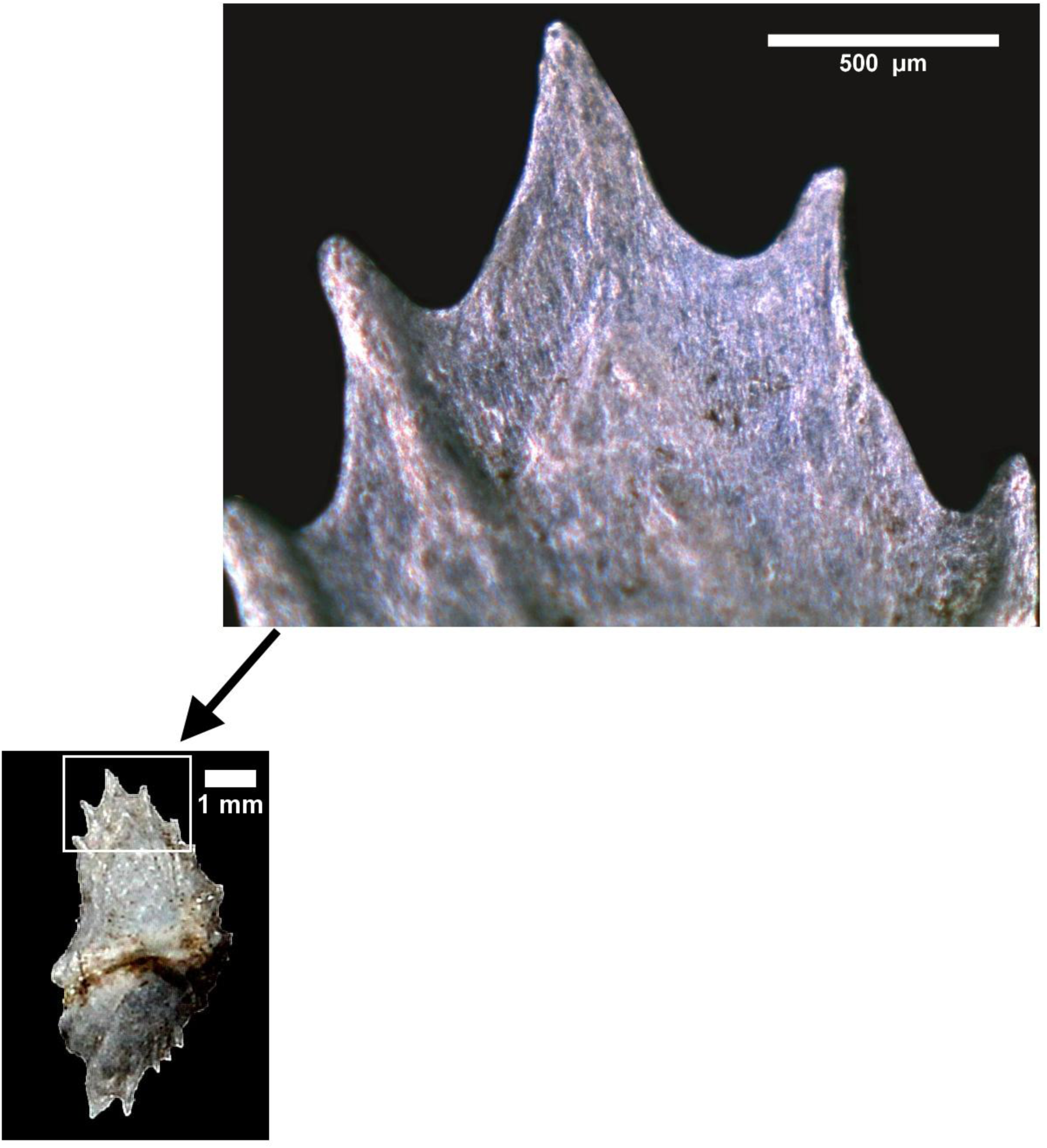
Typical lateral scute of male juvenile *Acipenser baerii*. Backgrounds were cleaned and contrast was enhanced using Adobe Photoshop.

**Fig. 52.**
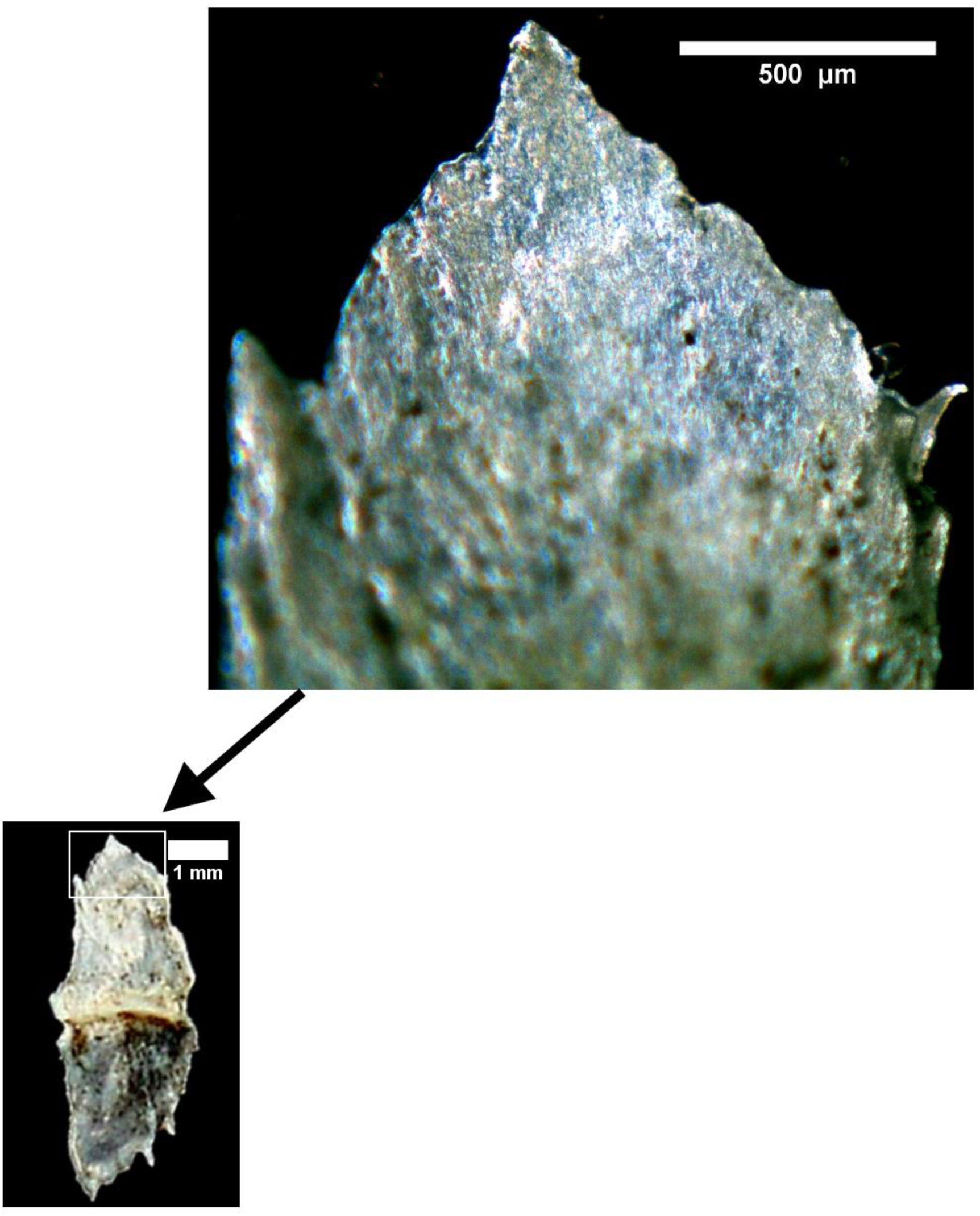
Typical lateral scute of female juvenile *Acipenser baerii*. Backgrounds were cleaned and contrast was enhanced using Adobe Photoshop.

**Fig. 53.**
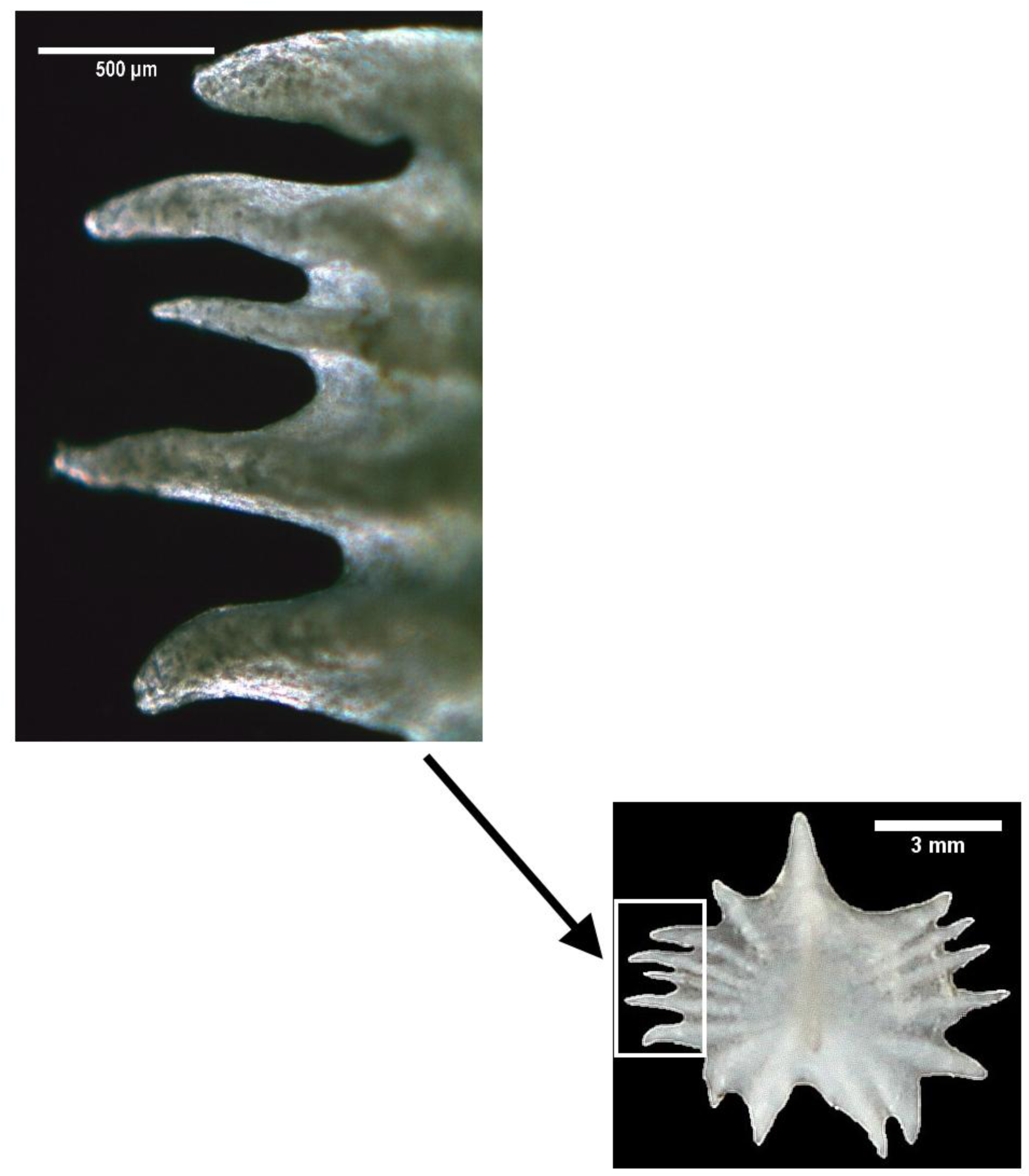
Typical abdominal scute of male juvenile *Acipenser baerii*. Backgrounds were cleaned and contrast was enhanced using Adobe Photoshop.

**Fig. 54.**
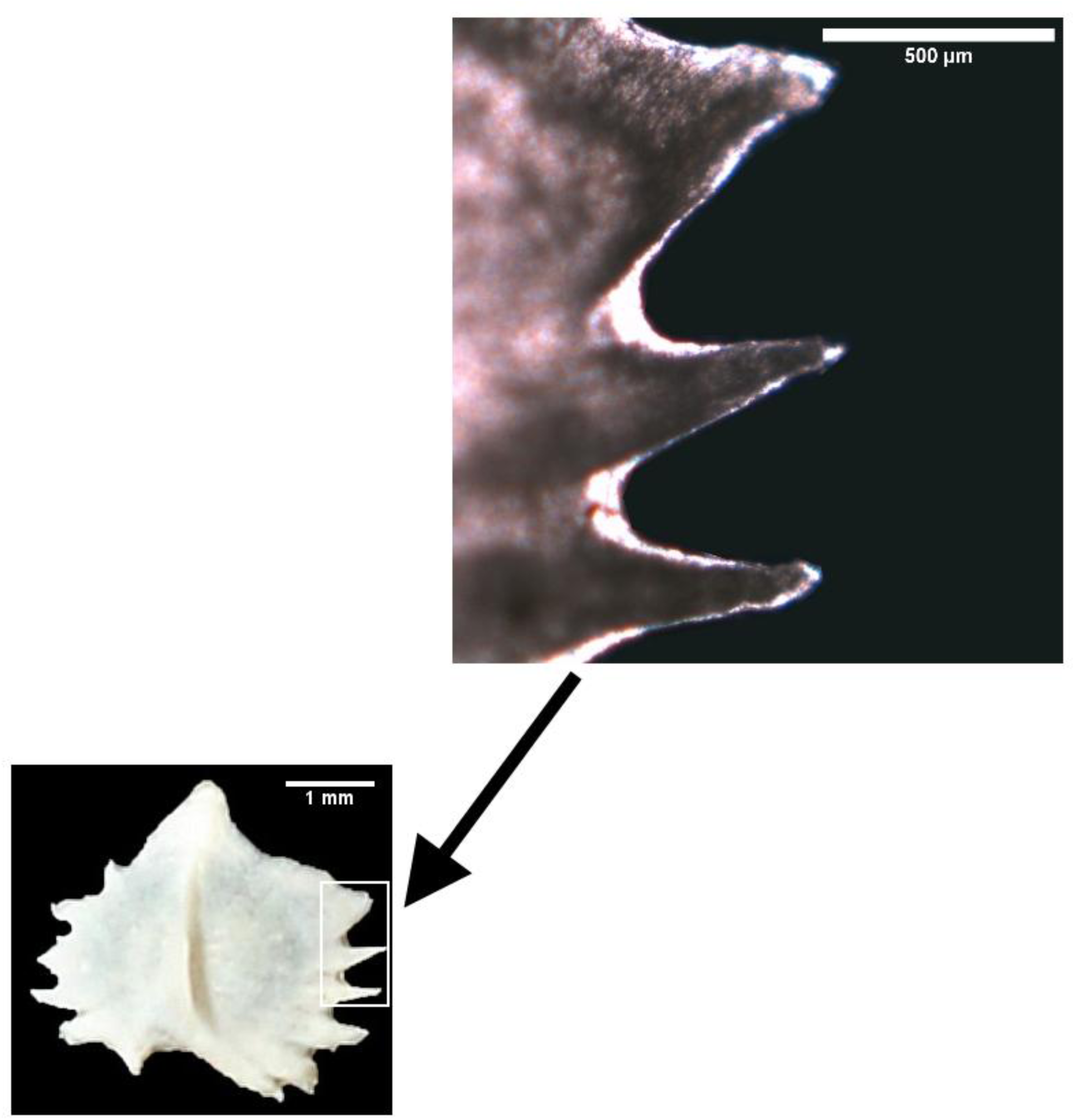
Typical abdominal scute of female juvenile *Acipenser baerii*. Backgrounds were cleaned and contrast was enhanced using Adobe Photoshop.

## CONCLUSION

We first found that sturgeon scutes have different forms (structure) depending on the sex. These relationships maintained in sturgeons of different ages and species. Thus, we affirm that sturgeons have external morphological differences depending on the sex, which was previously denied and considered impossible. Sex differences in the scutes forms are determined from a very early age (from 3 months). The observed relationship in the scutes forms can increase the economic efficiency of caviar aquaculture, because it is possible to identify males at earlier age to reject and not to spend expensive feed and electricity for male keeping. In addition, the earlier sex determination opens promising areas for the development of sturgeon specialized technologies, following the example of animal breeding. Early sturgeon sex determination would allow the stocking of rivers with the correct sex ratio and will contribute to more effective work to restore sturgeon wild populations.

Found dependencies using machine learning method open a perspective for creation of sex determination equipment using the artificial intelligence for image recognition based on neural networks.

Our studies were limited only to two sturgeon species. The question remains about the sex depending on scute structure in the other sturgeon species. For example, studies by (Wuertz et al., 2011) show that between juveniles of European (*A. sturio*) and Atlantic (*A. oxyrinchus*) sturgeons, there were differences in the scute structure, that were preserved in sexually mature individuals (Thieren et al., 2015). As is known, 25 from 27 sturgeon species have 5 scute rows. This allows us to hope for a positive result in detecting the sex depending on scute structure in other sturgeon species (for example, in White sturgeon, Atlantic sturgeon, Shovelnose sturgeon (*Scaphirynchus platorynchus*), Pallid sturgeon (*Scaphirynchus albus*), Alabama sturgeon (*Scaphirynchus suttkusi*) and other sturgeons inhabited in the US and Canada). We invite other researchers to participate in these studies.

## ACKNOWLEDGMENTS

The study was supported by projects of Belarusian State Agricultural Academy and Ministry of Agriculture of Belarus. I thank S.V. Rogovtsov, K.L. Shumsky, L.O. Atroschenko, A.V. Volynets, N. A. Surovets, V.V. Kalinin, L.E. Dubravsky, and A.V. Nekrulov (Belarusian State Agricultural Academy) for their help in the collection of samples. I am especially grateful to my wife, A.S. Barulina, for her accidental observation of sex-related differences in the structure of dorsal scutes of sterlets, which provided the concept of this study.

