## Supplementary Materials for "Scute structure reveals sex of juvenile and young sturgeon in caviar aquaculture technology"

Nikolai V. Barulin

### PROTOCOL FOR STERLET SEX DETERMINATION

Sex determination can be carried out on sterlet sturgeon when reaching a length of 40 cm. Special workplace should be arranged. The operator's workplace should include: special (metal or wooden) table for viewing the fish; fish tanks, which must be located in the immediate vicinity of the operator. Fish is placed on the stomach (head left or right relative to the operator). Fish must be kept in relative immobility during the whole process of sex determination, which can last from a few seconds to several minutes. We recommend not to feed the fish for 2-3 days before sex determination. We recommend to use anesthesia to reduce the fish stress. To view dorsal scutes, recommended to use a well-lit workplace, additionally using a manual or table magnifier with cross-polarized illumination, 5x magnification and 70 mm of lens diameter. In addition, you may need a white-light LED flashlight with a light output of 2000 lm and a headband binocular magnifier with backlighting in difficult cases.

Digital SLR camera can also be used in macro mode with enhanced flash, followed by fish tagging and individual analysis of images in ImageJ software or in any other graphic editor with the ability to adjust brightness and contrast, as well as inverting colors. We recommend to cover the head of fish with a cloth to prevent the fish eyes from the possible harmful effects of bright light.

To sex determination, it is necessary to examine the first five dorsal scutes (counting is from the head, the first scute tightly adjacent to the skull bone plates is not taken into account). It is necessary to pay attention to three main parameters:

1. Visual shape of the dorsal scute. One point go to each scute with oblate and oval shape (Fig. 1a). Point is not go for scute with elongated and rounded shape (Fig. 1b). Oblate and oval shape of scute should be characterized by: scute width more than scute length; conditional lines drawn through the scute endpoints formed an isosceles triangle (Fig. 2); scute blade separated from the scute at a less acute angle.

2. Visual shape of the dorsal scute teeth. One point go to each scute if it has a lot of sharp and elongated teeth (Fig. 1c). Point is not go for scute with short, rounded teeth or in their absence (Fig. 1d).

3. Narrow scutes distance. One total point go to scutes if they have narrow distance. Criterion for this parameter is the average distance between first five dorsal scutes, which should be less than ½ the average blade length of the dorsal scute (Fig. 3). This criterion is determined only in adult fishes.

Sterlet sturgeon is diagnosed as male, if the number of points are 5 or more, with a total score of 4 or less, sterlet sturgeon is diagnosed as female.

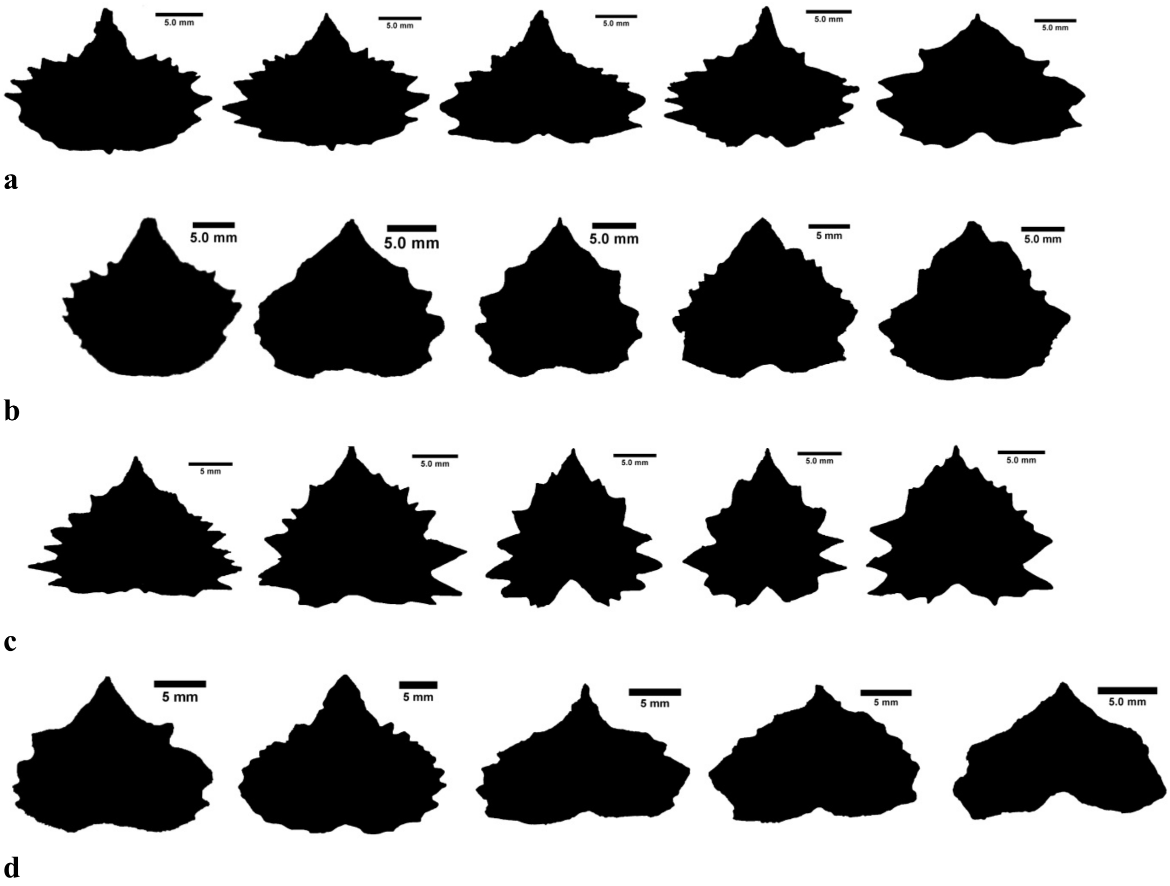


**Fig. 1.** Contour samples of the typical dorsal scute shapes of *Acipenser ruthenus* for sex determination. a – the dorsal scute is oblate, oval (1 point go to each scute); b – the dorsal scute is elongated, rounded (0 point); c – the dorsal scute teeth are sharp and elongated (1 point go to each scute); d – the dorsal scute teeth are short, rounded or not identified.

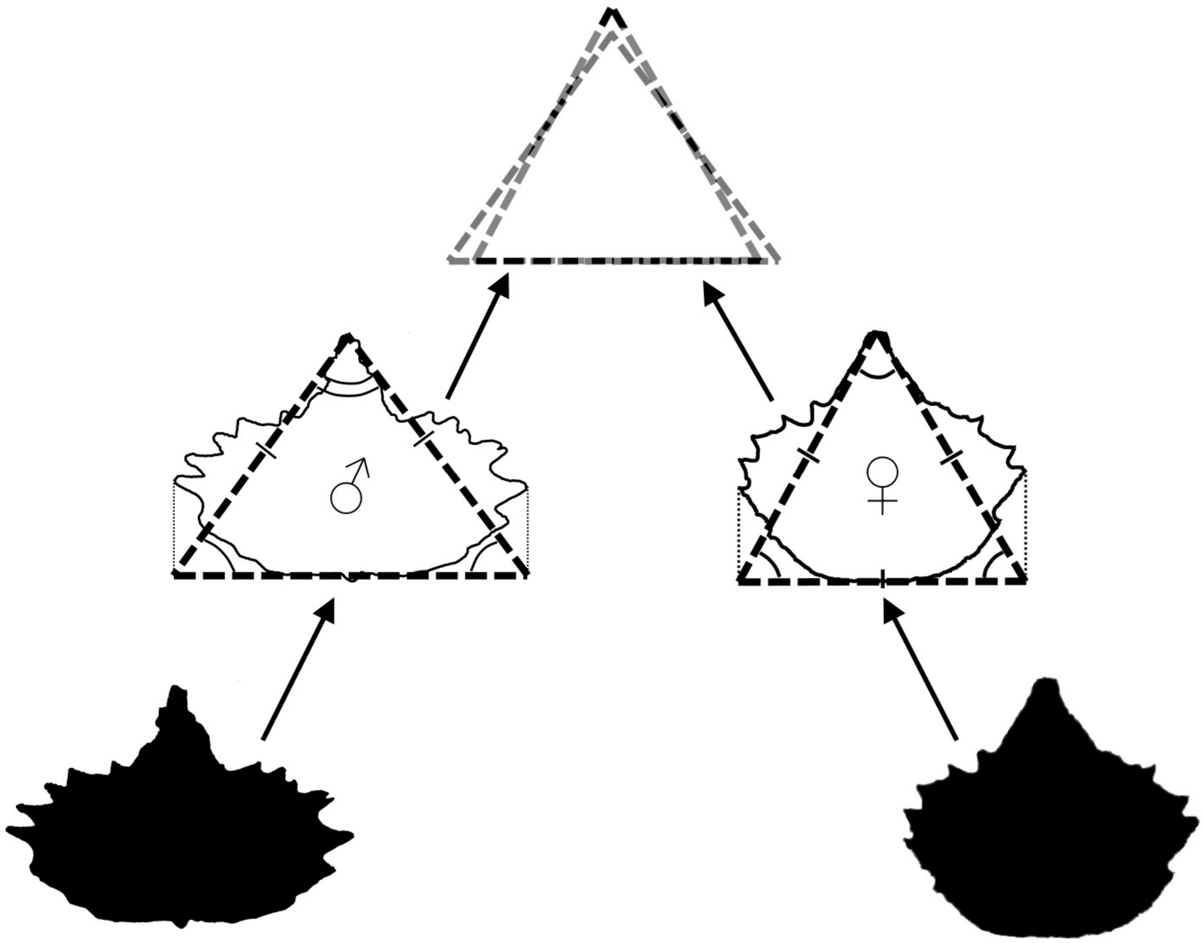


**Fig. 2.** Typical male scutes of *Acipenser ruthenus* had a broader and oblate (oval) shape; conditional lines drawn through the scute endpoints formed an isosceles triangle. Female scutes of *Acipenser ruthenus* had a more rounded shape, conditional lines drawn through the scute endpoints formed an equilateral triangle.

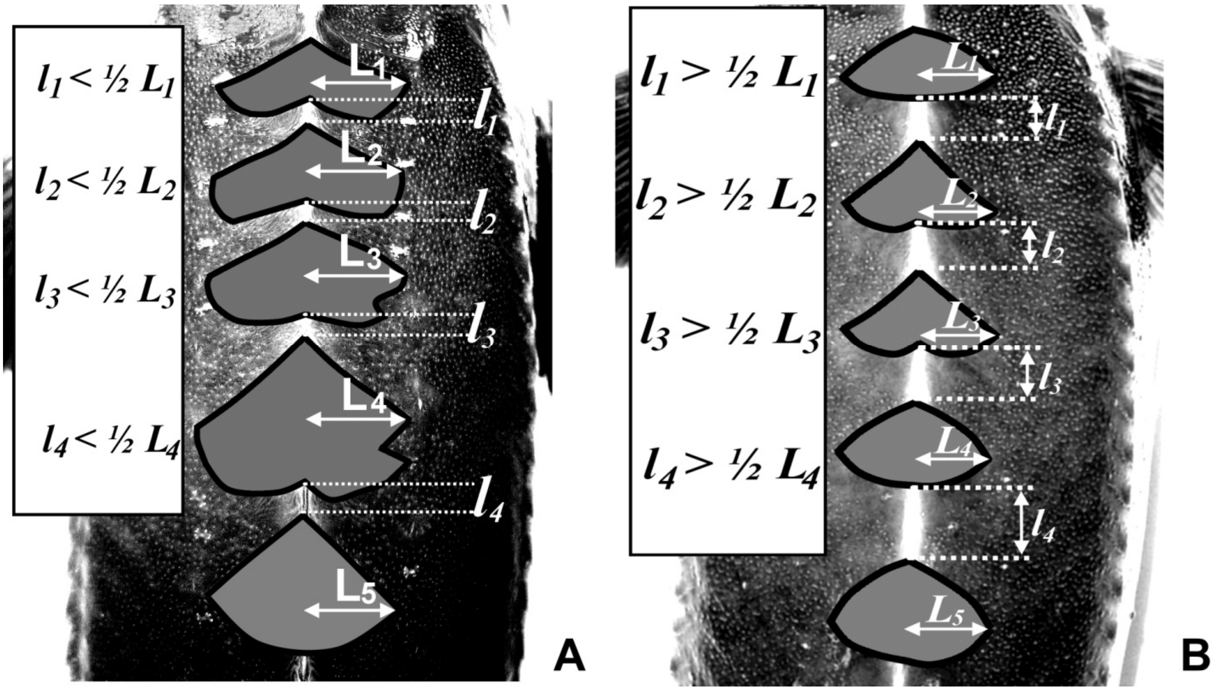


**Fig. 3.** Example of narrow scute distance. Criterion for this parameter is the average distance (*l*) between the first to fifth dorsal scutes, which should be less than ½ the average blade length (*L*) of dorsal scute (A)). This criterion is determined only in adult fishes.
